# Genome size is positively correlated with extinction risk in herbaceous angiosperms

**DOI:** 10.1101/2023.09.10.557053

**Authors:** Marybel Soto Gomez, Matilda J.M. Brown, Samuel Pironon, Pavel Veselý, Petr Bureš, Tammy L. Elliott, František Zedek, Jaume Pellicer, Félix Forest, Eimear Nic Lughadha, Ilia J. Leitch

## Abstract

- Angiosperms with large genomes experience nuclear-, cellular- and organism-level constraints that may limit their phenotypic plasticity and ecological niche. These constraints have been documented to vary across lineages, life-history strategies, ecogeographic patterns and environmental conditions. Therefore, we test the hypotheses that extinction risk is higher in large-genomed compared to small-genomed species, and that the effect of genome size varies across three selected covariates: life form, endemism, and climatic zones.
- We collated genome size and extinction risk information for a representative sample of angiosperms comprising 3,250 species, which we analyzed alongside life form, endemism and climate variables using a phylogenetic framework.
- Angiosperm genome size is positively correlated with extinction risk, a pattern driven by a signal in herbaceous but not woody species, regardless of climate and endemism. The influence of genome size is stronger in endemic herbaceous species, but is relatively homogenous across different climates. Beyond its indirect link via endemism and climate, genome size also influences extinction risk directly and significantly.
- Genome size may serve as a proxy for difficult-to-measure parameters associated with resilience and vulnerability in herbaceous angiosperms. Therefore, it merits further exploration as a useful biological attribute for understanding intrinsic extinction risk and augmenting plant conservation efforts.

## Introduction

Angiosperm genome size (nuclear DNA amount) is a highly variable character with long recognized implications for plant physiology, ecology and evolution (e.g., Sparrow & Miksche, 1961; Bennett, 1972; Novák *et al*., 2020). The remarkable diversity in angiosperm genome size spans ∼2,400-fold across the species documented to date, the largest range for any comparable group of eukaryotes (Pellicer *et al*., 2018). Mounting evidence has shown that genome size is involved in the scaling of organisms: from the subcellular level where it influences the duration of mitosis and meiosis (Bennett, 1971; Šímová & Herben, 2012; Zhukovskaya & Ivanov, 2022), to the cellular level where it determines minimum cell size and cell packing density (e.g., Roddy *et al*., 2020; Théroux-Rancourt *et al*., 2021), and the organismal level where it affects life-history strategies (e.g., Bennett, 1987; Veselý *et al*., 2012; Carta *et al*., 2022) and physiological parameters such as growth rate (e.g., Knight *et al*., 2005; Tenaillon *et al*., 2016; White *et al*., 2016) and photosynthetic efficiency (Beaulieu *et al*., 2007; Roddy *et al*., 2020). The cascading effects of genome size can in turn play a role in influencing where, when, and how plants grow and compete, thereby shaping community composition and distribution (Guignard *et al*., 2016; Bureš *et al*., 2022; Zhang *et al*., 2022). Taken together, these multiple lines of evidence raise the possibility that genome size may be associated with extinction risk in angiosperms, as originally argued by Vinogradov (2003).

Genome size diversity in angiosperms is largely driven by polyploidy followed by genome downsizing, and by the accumulation and removal of repetitive DNA (e.g., Grover & Wendel, 2010; Wendel, 2015). Despite the ubiquity of these two processes and the wide range of genome sizes observed, most angiosperms have small genomes (mode = 0.588 Gb/1C; Pellicer *et al*., 2018), leading to the hypothesis that larger genomes incur biological costs that drive selective pressure for genome downsizing (e.g., Leitch & Bennett, 2004; Knight *et al*., 2005; Wang *et al*., 2021). This hypothesised directional selection is partly influenced by genome size being positively correlated with cell size and inversely with cell division rate, so that species with large genomes are restricted to having large, slowly dividing cells with a lower maximum cell density (Francis *et al*., 2008; Roddy *et al*., 2020). Such cell-level effects can constrain phenotypic plasticity in large-genomed species by reducing the effective trait space (Faizullah *et al*., 2021). In turn this may exert selective pressure favouring smaller genomes, which have a lower minimum cell size (and hence potential for higher cell densities) that can either remain small or expand through processes like endopolyploidy and vacuole enlargement. Indeed, the wider phenotypic and ecological ranges of small-genomed species have been previously documented (e.g., Simonin & Roddy, 2018), whereas larger genomes tend to persist in conditions where selective pressures are relatively relaxed or compatible with the demands imposed by higher DNA amounts (e.g., Veselý *et al*., 2013; Bureš *et al*., 2022). These demands include meeting the biochemical costs of maintaining larger genomes, as nucleic acids are amongst the most nitrogen- and phosphorous-demanding molecules of the cell, leading to poor competitiveness of large-genomed taxa under nutrient-limiting conditions (Šmarda *et al*., 2013; Guignard *et al*., 2016; Peng *et al*., 2022). Overall, these observations point to several micro-and macro-evolutionary processes that impose selective disadvantages for large genomes, with potential downstream repercussions when confronted with the anthropogenic threats currently driving extinction risk (IPBES, 2019). For example, restricted phenotypic and physiological ranges may make large-genomed species less versatile in responding to protracted threats like climate change, while limits in ecological space, which frequently correspond with smaller geographic ranges (Slatyer *et al*., 2013), may increase extinction risk from stochastic and local-scale threats like land conversion.

A relationship between genome size and extinction risk across the tree of life was recovered with mixed support in the few large-scale studies that have examined this link. For example, no link was observed in amphibians (Pincheira-Donoso *et al*., 2023), possibly reflecting previous findings that the association between genome size and risk is complex and lineage-dependent across vertebrates (Vinogradov, 2004). Nevertheless, in angiosperms, large genomes were found to be maladaptive, and associated with extinction risk (Vinogradov, 2003) and rarity (Pandit *et al*., 2011). These broad patterns in plants were also reported at lower taxonomic levels, as in the Crassulaceae genus *Aeonium* (Brilhante *et al*., 2021). However, the relationship between angiosperm genome size and extinction risk has not been explicitly tested to date within a comprehensive evolutionary framework at a global scale, despite the considerable phylogenetic clustering of genome size across the angiosperm evolutionary tree (Carta *et al*., 2022). Moreover, few studies have explored the extent to which threat may be directly or indirectly associated with angiosperm genome size while considering its diverse covarying factors, which include genome evolution (e.g., polyploidy, repeat-sequence turnover; Wendel, 2015; Novák *et al*., 2020), physiology (e.g., photosynthetic efficiency, nutritional demands; Beaulieu *et al*., 2007; Guignard *et al*., 2016), life-history strategies (e.g., growth form; Bennett, 1987; Veselý *et al*., 2012; Carta *et al*., 2022), environment and geography (e.g., climate, range size; Bureš *et al*., 2022). The feasibility of such studies remains limited to date because of difficulties in obtaining consistently recorded information at broad taxonomic scales, hence the utility of resources like the World Checklist of Vascular Plants (WCVP; Govaerts *et al*., 2021) that compile relatively coarse but near complete datasets for multiple angiosperm characteristics. These include three genome size covariates –life form, climate zone, geographic distribution– that are independently associated with threat. Specifically, the scaling effects of genome size on organismal growth rate underlie the predominantly small genomes of annuals, as these species must complete their life cycle within a growing season (Bennett, 1987; Veselý *et al*., 2012). This constraint may be further exacerbated in environments that require rapid growth, such as temperate areas with short growing seasons, where annuals (therophytes) were found to be the second most threatened life form after hemicryptophytes (Le Roux *et al*., 2019). Environmental factors are also involved in the geographic patterns recently uncovered for angiosperm genome size, with narrower distributions found for larger-genomed than smaller-genomed taxa (Bureš *et al*., 2022), a noteworthy finding as range size is a well-documented correlate of past and contemporary extinctions (Gaston, 2003; Payne & Finnegan, 2007; Tanentzap, 2017).

Plants underpin life on earth, but two in five species are predicted to be threatened with extinction (Nic Lughadha *et al*., 2020), making it critical to preserve these elements of biodiversity alongside the ecosystem services they provide and unique evolutionary histories they represent (Antonelli *et al*., 2020). Effective species conservation requires information on their extinction risk, such as that provided by the International Union for Conservation of Nature (IUCN) Red List of Threatened Species (Red List; IUCN, 2022), the most authoritative source on the global conservation status of species. Red List assessments employ five criteria that incorporate current and temporal information on range size, and where available, population size and demographic change (IUCN, 2013). Genetic data are not explicitly considered in the Red List, but there is some evidence that criterion thresholds may reflect genetic diversity, and by extension, evolutionary potential in plants (Rivers *et al*., 2014; but see Schmidt *et al*., 2023 for vertebrates). Despite substantial progress in extending Red List coverage of plants (Bachman *et al*., 2018; 2019), only 18% of known species have been assessed to date (Bachman *et al*., 2023). Recent work to automate extinction risk assessment has achieved reasonable performance using a range of predictors, including geographic, environmental and morphological data (Pelletier *et al*., 2018; Walker *et al*., 2022). Nevertheless, there are outstanding knowledge gaps in identifying at-risk species and the factors that threaten them over time (Nic Lughadha *et al*., 2020). Therefore, if found to be associated with risk, genome size may be a useful genetic variable for understanding intrinsic vulnerability in angiosperms, for helping to prioritize Red List assessments and for enhancing the accuracy of predictive models.

Here we used a comprehensive evolutionary framework to test how genome size may influence extinction risk in angiosperms, leveraging the largest global datasets available for plant genome size, extinction risk, and newly released information on life form, climate zone and geographic distribution from WCVP (Govaerts *et al*., 2021). We included the latter three variables as representatives of a suite of processes previously associated with both genome size and extinction risk, and for which it was possible to obtain consistent angiosperm-wide data. In contrast to previous studies aiming to understand the impact of genome size on plant conservation through rarity patterns (Pandit *et al*., 2011; 2014) or using taxonomy as a proxy for evolutionary history (Vinogradov, 2003), we employed a phylogenetically-informed approach to test two hypotheses at a global scale: (i) angiosperm genome size and extinction risk are positively correlated, and (ii) the extent of risk in large-genomed species varies across life forms, climatic zones and range sizes. Finally, we identify a subset of species currently lacking a threat assessment but facing a potentially heightened extinction risk based on their genome size and covariate combinations.

## Materials and Methods

### Taxonomic reconciliation

We used WCVP as the taxonomic basis for reconciling sample names and taxonomic ranks across the various data sources employed here. Specifically, we collated (i) genome size data from the Plant DNA C-values Database (Leitch *et al*., 2019) and a newly published dataset (Bureš *et al*., 2022); (ii) global extinction risk data from the Red List (IUCN, 2022); (iii) life form, climate zone, and distribution data from WCVP; and (iv) phylogenetic information from a sampling of 100 angiosperm phylogenies (Forest, 2023).

We determined the taxonomic status of samples in the genome size dataset assembled here (see below), confirming whether these had names accepted in WCVP or non-accepted names requiring reconciliation. To do so, we used the R package rWCVP (Brown *et al*., 2023), using its “accepted plant name id” output to link species in the genome size dataset to available data for that species in the remaining datasets employed here (details in Methods **S1**).

### Genome size data

We constructed a genome size dataset comprising 15,158 species (Table **S1**). We maximized taxon sampling by collating data for 8,581 species from Leitch *et al*. (2019) with an additional 6,577 species from Bureš *et al*. (2022), after taxonomic reconciliation of each dataset to WCVP. We removed sample redundancy by selecting a single placeholder for species with multiple accessions, preferentially retaining those with the smallest genome size estimate. We adopted this conservative approach to minimize the potential of generating type I errors (i.e., false positives regarding our hypothesized effects of large genomes on extinction risk), which may arise from retaining the largest genome size estimate available for a species.

### Extinction risk, life form, climate zone, and endemism data

Considering the 15,158 species in the genome size dataset, extinction risk information was available for 3,394 of these, life form for 15,141; climate zone for 14,003; and distribution for 15,155. We focused most analytical effort on a subset of 3,250 species that were scored for all five variables.

Six of seven Red List Categories describing extinction risk were represented in the 3,250-species dataset (only the Extinct category was absent). Three of the categories in our sampling – Vulnerable (VU; n = 243), Endangered (EN; n = 287), Critically Endangered (CR; n = 166)– are collectively considered to be threatened in Red List criteria, and respectively face a high, very high and extremely high extinction risk (Mace *et al*., 2008). To qualify as such, species must meet at least one of five criteria concerning population size, geographic range and extinction probability, at a level corresponding to one of the three threatened categories (IUCN, 2013). A fourth Red List Category in our sampling –Extinct in the Wild (EW; n = 7)– is also included in Red List estimates of threatened species as EW taxa are mostly re-assessed as VU, EN or CR following rediscovery (Humphreys *et al*., 2019) or successful reintroduction (IUCN, 2019). The remaining categories in our sampling –Least Concern (LC; n = 2,388), Near Threatened (NT; n = 159)– comprise non-threatened species that were evaluated against Red List criteria and did not qualify at the level required to be considered threatened (IUCN, 2019).

To maintain viable sample sizes across data partitions in analyses that also included life form, climate zone, and distribution variables (see below and Table **S2**), we treated extinction risk as a binary response variable comprising threatened and non-threatened species across four varying threat thresholds. These dichotomizations were designed to capture the Red List definition of threatened as a point of reference, and to test the effects of shifting this threshold to account for differences in threat levels across categories. Thus, for the point of reference threshold, the non-threatened grouping comprised LC and NT species, and the threatened grouping comprised VU, EN, CR and EW species. For the three comparison dichotomizations, we lowered the threat threshold (by shifting NT species into the threatened grouping), increased the threshold (by shifting VU species into the non-threatened grouping), or polarized the threshold (by excluding NT species). The dichotomization of Red List Categories additionally facilitates comparison of our findings with those from previous plant and animal studies that used this approach, for example, when identifying Important Plant Areas (Darbyshire *et al*., 2017), predicting the probability of threat in species not yet assessed by the Red List (Pelletier *et al*., 2018), and characterizing the influence of genome size on amphibian extinction risk (Pincheira-Donoso *et al*., 2023).

We obtained life form, climate zone, and distribution data from WCVP, additionally collecting life form information from Bureš *et al*. (2022) when unavailable in WCVP. The life form classifications in both datasets follow the Raunkiær (1934) system, which we simplified following Humphreys *et al*. (2019), with modifications to produce two broad categories of biological significance to angiosperm genome size: herbaceous and woody (Beaulieu *et al*., 2010; Methods **S2**). We aggregated nine WCVP climate zones into four groupings: tropical, subtropical, temperate, and desert areas (Methods **S2**). We used WCVP geographical data to score species as endemic if their native distribution is restricted to a single botanical country (i.e., Level 3 of the TDWG World Geographic Scheme for Recording Plant Distributions; Brummitt *et al*., 2001); the remaining species were scored as non-endemic. This binary coding was strongly correlated with point-derived range size estimates, while overcoming the lack of point data for 500 species in our sampling (Methods **S2**, Fig. **S1**).

### Angiosperm phylogenies

We adapted the species-level angiosperm phylogenies of Forest (2023) to use as input in phylogenetically-informed statistical analyses. We used all 100 of these phylogenies to capture the uncertain phylogenetic placement of 935 (28.8%) species in our dataset that lacked phylogenetic data. The original phylogenies comprise all 329,798 angiosperm species recognized in WCVP (assembly details in Methods **S3**). Here we prepared reduced versions comprising only the 3,250 species in our dataset, with additional updates to fully bifurcate and rescale the phylogenies to a total height of 1.0 for statistical analyses (Methods **S3**).

### Data representativeness

We used four approaches to characterize the representativeness of the different data types encompassed by our 3,250-species dataset relative to angiosperms with available information, and to test for potential effects from imbalanced representation (Methods **S4**). First, we used the *D* statistic (Fritz & Purvis, 2010) to test whether the 993 genera represented in our dataset were phylogenetically clustered (i.e., concentrated in particular clades) or overdispersed (i.e., evenly distributed) across the 13,503 genera recognized in WCVP. Second, we used the 202,743 species in WCVP with available life form, climate zone and endemism information as a baseline for estimating a factor of representation for the equivalent data partitions in our sampling. Third, we assessed our coverage of known angiosperm genome sizes by comparing the distribution in our sampling to that of 15,167 species with available information (also collated here; Table **S1**).

Finally, we tested for possible effects from imbalanced proportions of non-threatened and threatened species by characterizing genome size differences between these two groups in our sampling, compared to 999 randomly down-sampled subsets with equal threat status proportions.

### Relationship between genome size and extinction risk

We performed an ANOVA followed by Tukey’s range test to determine whether average genome size differed significantly amongst individual Red List Categories. We also constructed phylogenetic generalized linear models (PGLMs) to characterize the association between genome size and extinction risk, and how it varies across life forms, climate zones and endemism. We modelled extinction risk as a binary response variable (i.e., threatened or non-threatened; see above) in 27 distinct logistic regressions that included genome size as the sole predictor, or together with life form, climate zone and endemism (Table **1**). We implemented all 27 models using the Red List definition of threatened as a point of reference for dichotomizing risk, and three varying threat thresholds for comparison. To account for phylogenetic uncertainty in our sampling, we ran 100 independent analyses for all models, each using a different input phylogeny.

**Table 1.**
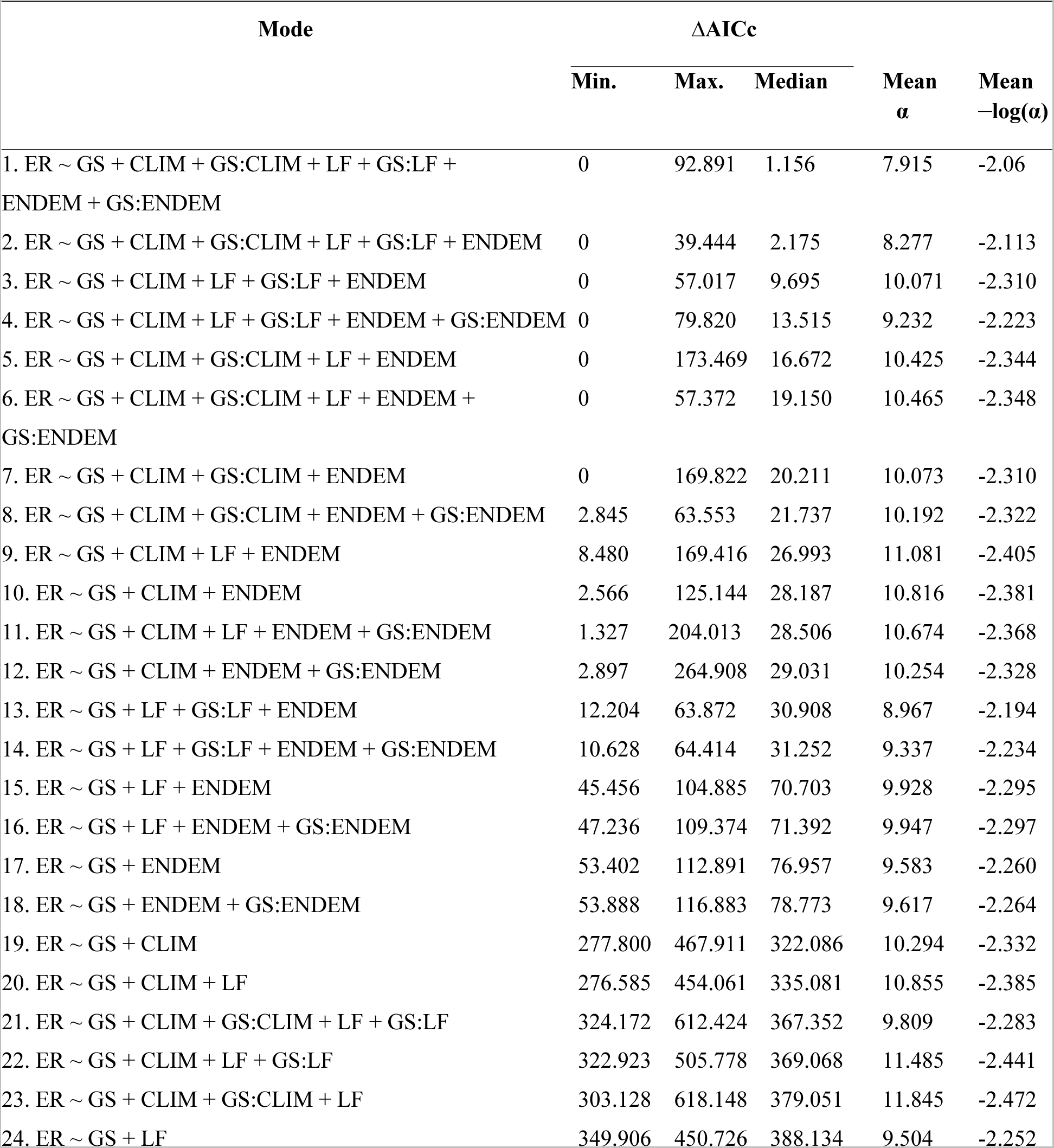
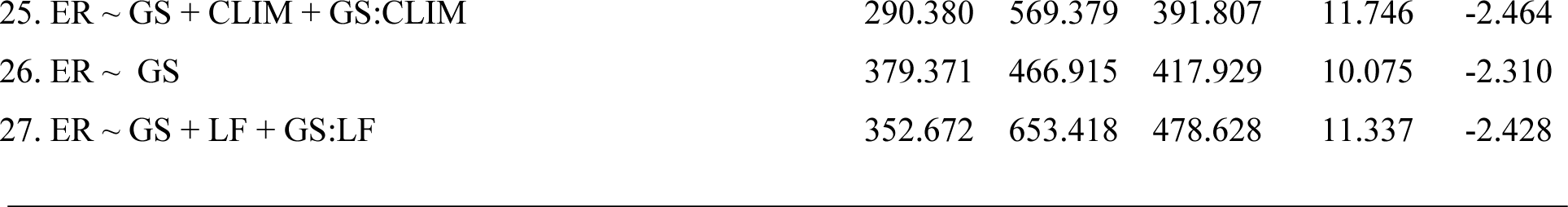
Description, selection, and phylogenetic signal of the 27 models tested in phylogenetic logistic regressions using the Red List definition of threat as a point of reference for dichotomizing the response variable into groups comprising non-threatened (i.e., Least Concern, Near Threatened) and threatened (i.e., Vulnerable, Endangered, Critically Endangered, Extinct in the Wild) species (three comparison threat thresholds are summarized in Table **S5**). Models represent all combinations possible when adding life form, climate zone and endemism as explanatory variables to a baseline model with genome size as the sole predictor. Models are arranged in decreasing order of best fit based on median ΔAICc values, obtained by first ranking models within each of the 100 variant phylogenies used as input in analyses, and then summarized by model across all trees (see Fig. **S2**). The first seven models (depicted in Fig. **2**) were found to be the best (i.e., ΔAICc = 0) in at least one phylogeny. Mean phylogenetic signal (α) was obtained by summarizing across the 100 separate analyses conducted for each model and then used to calculate –log(α). Abbreviations: ER = extinction risk, GS = genome size, CLIM = climate zone, LF = life form, ENDEM = endemism (proxy for range size).

| Mode | $\Delta AICc$ | | | Mean $\alpha$ | Mean $-\log(\alpha)$ |
| --- | --- | --- | --- | --- | --- |
|  | Min. | Max. | Median |  |  |
| 1. ER ~ GS + CLIM + GS:CLIM + LF + GS:LF + ENDEM + GS:ENDEM | 0 | 92.891 | 1.156 | 7.915 | -2.06 |
| 2. ER ~ GS + CLIM + GS:CLIM + LF + GS:LF + ENDEM | 0 | 39.444 | 2.175 | 8.277 | -2.113 |
| 3. ER ~ GS + CLIM + LF + GS:LF + ENDEM | 0 | 57.017 | 9.695 | 10.071 | -2.310 |
| 4. ER ~ GS + CLIM + LF + GS:LF + ENDEM + GS:ENDEM | 0 | 79.820 | 13.515 | 9.232 | -2.223 |
| 5. ER ~ GS + CLIM + GS:CLIM + LF + ENDEM | 0 | 173.469 | 16.672 | 10.425 | -2.344 |
| 6. ER ~ GS + CLIM + GS:CLIM + LF + ENDEM + GS:ENDEM | 0 | 57.372 | 19.150 | 10.465 | -2.348 |
| 7. ER ~ GS + CLIM + GS:CLIM + ENDEM | 0 | 169.822 | 20.211 | 10.073 | -2.310 |
| 8. ER ~ GS + CLIM + GS:CLIM + ENDEM + GS:ENDEM | 2.845 | 63.553 | 21.737 | 10.192 | -2.322 |
| 9. ER ~ GS + CLIM + LF + ENDEM | 8.480 | 169.416 | 26.993 | 11.081 | -2.405 |
| 10. ER ~ GS + CLIM + ENDEM | 2.566 | 125.144 | 28.187 | 10.816 | -2.381 |
| 11. ER ~ GS + CLIM + LF + ENDEM + GS:ENDEM | 1.327 | 204.013 | 28.506 | 10.674 | -2.368 |
| 12. ER ~ GS + CLIM + ENDEM + GS:ENDEM | 2.897 | 264.908 | 29.031 | 10.254 | -2.328 |
| 13. ER ~ GS + LF + GS:LF + ENDEM | 12.204 | 63.872 | 30.908 | 8.967 | -2.194 |
| 14. ER ~ GS + LF + GS:LF + ENDEM + GS:ENDEM | 10.628 | 64.414 | 31.252 | 9.337 | -2.234 |
| 15. ER ~ GS + LF + ENDEM | 45.456 | 104.885 | 70.703 | 9.928 | -2.295 |
| 16. ER ~ GS + LF + ENDEM + GS:ENDEM | 47.236 | 109.374 | 71.392 | 9.947 | -2.297 |
| 17. ER ~ GS + ENDEM | 53.402 | 112.891 | 76.957 | 9.583 | -2.260 |
| 18. ER ~ GS + ENDEM + GS:ENDEM | 53.888 | 116.883 | 78.773 | 9.617 | -2.264 |
| 19. ER ~ GS + CLIM | 277.800 | 467.911 | 322.086 | 10.294 | -2.332 |
| 20. ER ~ GS + CLIM + LF | 276.585 | 454.061 | 335.081 | 10.855 | -2.385 |
| 21. ER ~ GS + CLIM + GS:CLIM + LF + GS:LF | 324.172 | 612.424 | 367.352 | 9.809 | -2.283 |
| 22. ER ~ GS + CLIM + LF + GS:LF | 322.923 | 505.778 | 369.068 | 11.485 | -2.441 |
| 23. ER ~ GS + CLIM + GS:CLIM + LF | 303.128 | 618.148 | 379.051 | 11.845 | -2.472 |
| 24. ER ~ GS + LF | 349.906 | 450.726 | 388.134 | 9.504 | -2.252 |

|  |  |  |  |  |  |  |
| --- | --- | --- | --- | --- | --- | --- |
| 416 | 25. ER ~ GS + CLIM + GS:CLIM | 290.380 | 569.379 | 391.807 | 11.746 | -2.464 |
| 417 | 26. ER ~ GS | 379.371 | 466.915 | 417.929 | 10.075 | -2.310 |
| 418 | 27. ER ~ GS + LF + GS:LF | 352.672 | 653.418 | 478.628 | 11.337 | -2.428 |
| 419 |  |  |  |  |  |  |

We conducted PGLMs using the phyloglm function in the R package phylolm (Ho & Ané, 2014a,b) with genome sizes log10-transformed. We first characterized the effect of genome size alone on extinction risk using a baseline model (model 26, Table **1**) with default settings across the four varying threat thresholds for dichotomizing the response variable. This model produced bimodally distributed estimates for one threat threshold, which required additional parameter optimization (Methods **S5).** We then modelled extinction risk across the remaining 26 model formulae (Table **1**), which represent all the combinations possible when adding life form, climate zone and endemism (as additive or interacting explanatory variables) to the baseline model. Non-continuous variables analyzed in phyloglm require a reference category: we selected herbaceous (for life form), desert (for climate zone), and endemic (for endemism). We adjusted the “btol” parameter in cases when the linear predictor was reached during optimization procedures, but otherwise used default settings. We performed model selection using AICc (Akaike Information Criterion corrected for small sample sizes; Burnham & Anderson, 2002); full details in Fig. **S2**, Methods **S6**. For the final model set, we again used phyloglm to run 2,000 bootstrap replicates (Ives & Garland, 2010), obtaining 95% confidence intervals for model variables.

We used the findings from PGLMs to (i) estimate the difference in probability of threat between thresholds set for very small and large genomes (Fig. **S3**), and (ii) conduct a post-hoc identification of species in the genome size dataset that lack a Red List assessment but face a potentially heightened extinction risk based on their genome size and covariate combinations. Following the criteria of Leitch *et al*. (1998) for categorizing angiosperm genome size, we considered very small genomes to be ≤1.18 Gb/1C, which is ≤2 times the modal 1C-value documented for angiosperms (0.588 Gb; Pellicer *et al.,* 2018), and large genomes to be ≥11.76 Gb/1C, which is ≥20 times the mode. Whereas selecting a biologically relevant threshold for very small genomes is difficult because of the relatively low physiological costs they impose (Simonin & Roddy, 2018), our large genome threshold is comparable to 10 Gb/1C –where fundamental shifts in genome evolutionary dynamics start occurring (Novák *et al*., 2020)– and to 8 Gb/1C –where the diversity of guard cell length and vein density start decreasing (Simonin & Roddy, 2018)–.

### Direct and indirect effects of genome size on extinction risk

We used confirmatory phylogenetic path analyses (PPA; von Hardenberg & Gonzalez-Voyer, 2013) to test whether angiosperm extinction risk is directly or indirectly associated with genome size, life form and endemism across climate zones. We constructed and tested four “causal models” (i.e., directed acyclic graphs; Fig. **S4**) using the R package phylopath (van der Bijl, 2018) with the method “logistic_MPLE” (details in Methods **S7**). We used the Red List threat threshold to obtain non-threatened (i.e., LC, NT) and threatened (i.e., VU, EN, CR, EW) groupings, and performed separate analyses using 100 different phylogenies as input to account for phylogenetic uncertainty. We used ΔCICc (C-statistic information criterion corrected for small sample sizes) to rank and discard causal models with ΔCICc >2 (van der Bijl, 2018). We then bootstrapped the best causal models using 500 replicates to obtain 95% confidence intervals. Coefficients were considered to be significant if their confidence intervals excluded zero (e.g. Guo *et al*., 2019).

## Results

### Species representativeness in the genome size dataset

The representativeness of the 3,250-species dataset differed across sampled variables when compared to angiosperm diversity, but small and large genomes were sampled in sufficient numbers across all threat-lifeform-climate-endemism combinations to permit confident estimates of genome size effects and interactions with these covariates (Table **S3**). Our sampling contained an overrepresentation of non-endemic species, particularly for temperate herbaceous and woody species (by factors of 1.94 and 4, respectively; Table **S4**). Endemic species were underrepresented across climate zones by factors of 0.28–0.62 for herbs and 0.41–0.91 for woody species (Table **S4**).

The 993 genera represented in our dataset were not strongly clustered across the 13,503 angiosperm genera recognized in WCVP (*D* = 0.665, range = 0.652–0.679, *p* < 0.001 for 0< *D* <1 across 100 variant phylogenies), indicating that our sampling was not strongly biased taxonomically (Fig. **S5**). Though slightly underrepresented for large genomes, our sampling (Gb/1C range = 0.08–73.01, mean = 3.54, median = 1.08; Table **S3**) covered most of the genome size diversity documented for angiosperms (Gb/1C range = 0.064–149.0, mean = 3.94, median = 1.37; Table **S1**, Fig. **S6**). Finally, using the point of reference threshold to dichotomize extinction risk, we found that the difference in mean genome size between threatened (n = 703) and non-threatened (n = 2,547) species in our dataset was nearly identical to the mean difference across 999 randomly down-sampled subsets that equalized the proportions of the two threat groupings (Fig. **S7**). This indicates that our analyses were likely unaffected by the relatively imbalanced sampling of non-threatened and threatened species.

### Probability of threat as a function of genome size

Average genome size differed significantly amongst Red List Categories (ANOVA: F = 13.1, df = 5, 3244, *p* <0.001; Fig. **1a**). Tukey’s range test showed that LC species were significantly different from EN and CR species; NT species also grouped with EN and CR, while VU species were intermediate (Fig. **1a**; note that we omitted EW species from this test due to a small sample size of seven). A baseline phylogenetic logistic regression model showed that genome size and extinction risk were significantly and positively correlated using the Red List threat threshold for dichotomizing extinction risk (Fig. **1b**). This relationship was consistent with the three comparison risk dichotomizations: only the intercept term (and not the slope) differed substantially across models, as expected when varying the threat threshold (Table **2**). Using the point of reference threshold, the probability of extinction risk increased by 0.28 from small-genomed to large-genomed species (Fig. **1b**).

**Figure 1.**
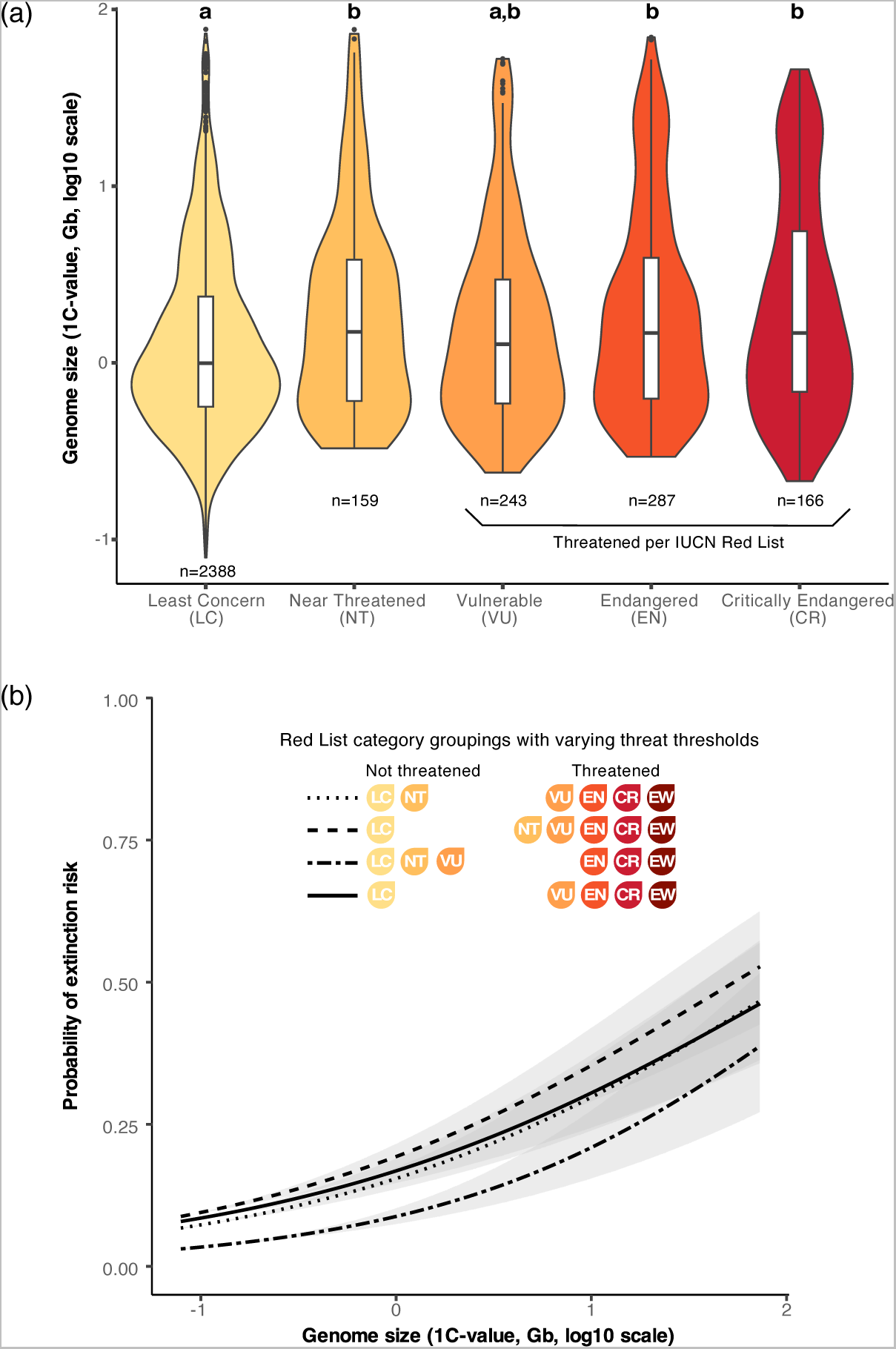
Characterization of the relationship between angiosperm genome size and extinction risk based on the 3,250-species dataset assembled here. (**a**) Distribution of genome sizes across Red List Categories (omitting Extinct in the Wild, EW, due to a small sample size of seven species). Significance was tested using ANOVA and Tukey’s range test. **(b)** Phylogenetic logistic regression curves predicting the probability of threat as a function of genome size using the Red List definition of threat as a point of reference for aggregating species into non-threatened or threatened groupings (shown as a dotted line) and using three additional threat thresholds for comparison (shown as solid, dashed and double-dashed lines). Curves represent mean coefficients summarized across the 100 different phylogenies used as input for each variant threat threshold; the shaded areas indicate 95% confidence intervals from bootstrap analyses.

**Table 2.** Consensus coefficients for a baseline phylogenetic logistic regression model characterizing angiosperm extinction risk as a function of genome size across four varying threat thresholds for dichotomizing the response variable (depicted in Fig. **1b**). The point of reference threshold reflects the Red List definition of threat for grouping species into non-threatened (i.e., Least Concern, Near Threatened) and threatened (i.e., Vulnerable, Endangered, Critically Endangered, Extinct in the Wild) groupings. The three comparison binarizations represent threat thresholds that are lower (by shifting Near Threatened species into the threatened grouping), higher (by shifting Vulnerable species into the non-threatened grouping) or polarizing (by excluding Near Threatened species altogether). Values represent mean coefficients from 100 separate analyses conducted for each model using different phylogenies as input.

|  |  |  |  |  |  |
| --- | --- | --- | --- | --- | --- |
| 431 |  |  |  |  |  |
| 432 |  | **Point of reference** | **Lower threat** | **Higher threat** | **Polarizing** |
| 433 |  | **threat threshold** | **threshold** | **threshold** | **threat threshold** |
| 434 |  |  |  |  |  |
| 435 | Intercept | -1.703 | -1.434 | -2.336 | -1.605 |
| 436 | Slope | 0.841 | 0.826 | 1.001 | 0.778 |
| 437 |  |  |  |  |  |

### Probability of threat as a function of genome size, life form, climate zone, and endemism

Model fit improved in phylogenetic logistic regressions that included genome size alongside life form, climate zone, and endemism as predictors of extinction risk, regardless of the threat threshold used to dichotomize the response variable (Tables **1**, **S5**). Of the 27 models tested, we retained two best models when using the Red List definition of threat as a point of reference threat threshold (Fig. **2**). One best model included genome size and the remaining three variables as both additive and interaction terms; the second model differed by lacking a genome size-endemism interaction term (Table **1**, Fig. **2**). We retained three best models in regressions using higher and polarizing threat thresholds for comparison to the point of reference, and one best model when using a lower threshold (Table **S5**, Figs. **S8**–**S10**). Coefficients were approximately normally distributed and yielded consistent predictions across the 100 analyses conducted for each of the best models across the four varying threat thresholds (Table **S6**, Figs. **S11**–**S19**).

**Figure 2.**
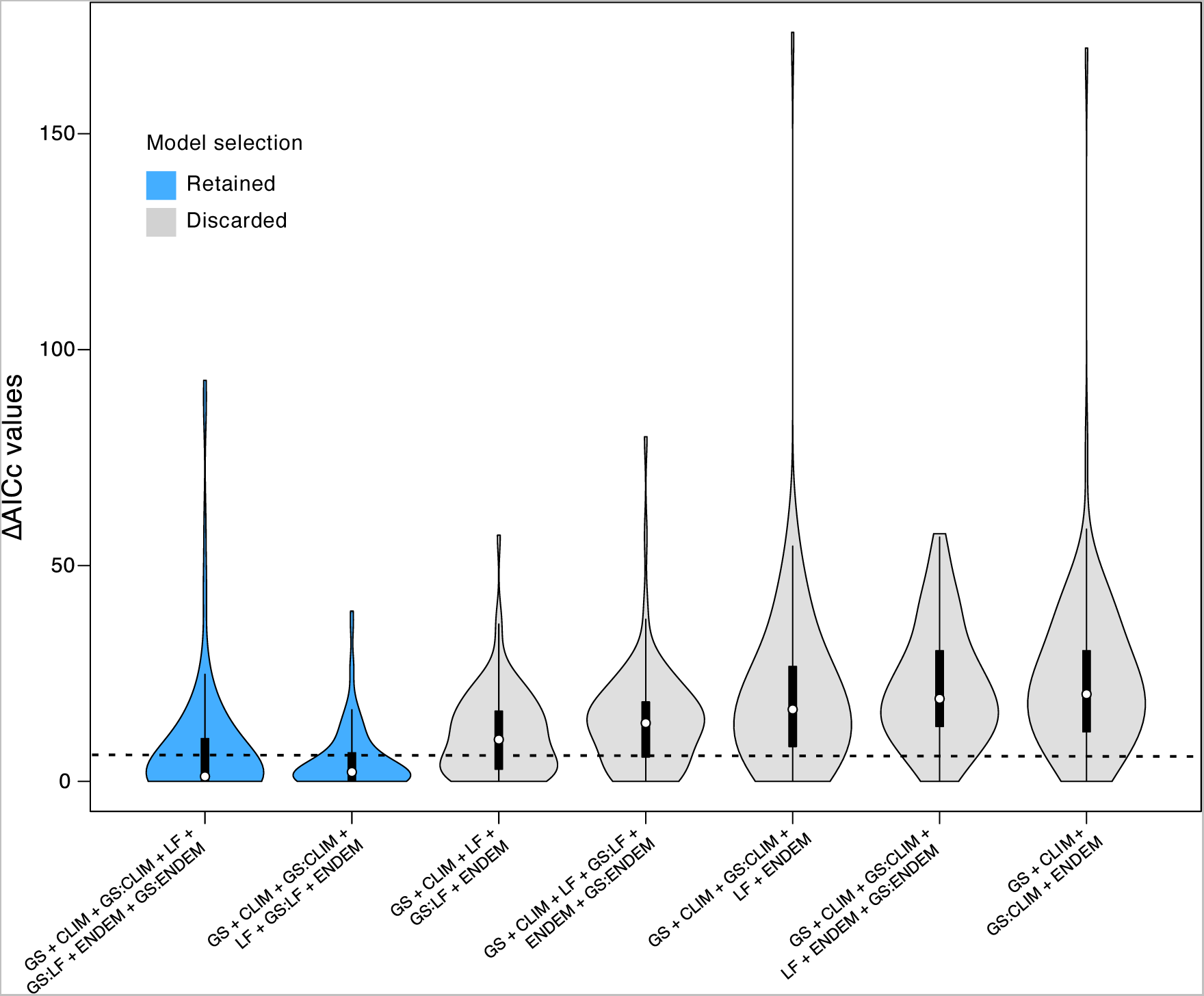
Model selection based on ΔAICc values estimated from phylogenetic logistic regressions using the Red List definition of threat as a point of reference for dichotomizing the response variable into a non-threatened group (comprising Least Concern and Near Threatened species) and a threatened group (comprising Vulnerable, Endangered, Critically Endangered and Extinct in the Wild species); model selection for three comparison threat groupings is shown in Figs. **S8**–**S10**. ΔAICc values were calculated by first ranking all 27 tested models within each of the 100 different phylogenies used as input in analyses, and then summarizing by model across all trees (as shown in Fig. **S2**). The plot shows the distribution of ΔAICc values for the seven (of 27) models that had the best AICc value (i.e., ΔAICc = 0) in at least one of the phylogenies (see Table **1** for support values of the remaining models). Models are ordered by increasing median ΔAICc values. The dashed line at ΔAICc = 6 indicates that models with a median ΔAICc below this cut-off were retained. Abbreviations: GS = genome size, CLIM = climate zone, LF = life form, ENDEM = endemism (proxy for range size).

All best models showed that genome size and extinction risk were positively correlated in both endemic and non-endemic herbaceous species across climate zones (Figs. **3, S20**–**22**). Averaging predictions across the best models for the point of reference threat threshold, this relationship was strongest for herbaceous species that are endemic to a single botanical country, where the threat probability increased by 52.2–66.4% between very small (≤1.18 Gb/1C) and large (≥11.76 Gb/1C) genomes across climate zones (Table **3**, Fig. **3**). For non-endemic herbs, the threat probability increased by 27.8–54.7% (Table **3**, Fig. **3**). However, in woody species, the threat probability remained nearly constant with increasing genome size, regardless of climate and endemism (Fig. **3**). We found consistent patterns for both herbaceous and woody species using the three comparison threat thresholds (Figs. **S20**–**S22**), which differed more substantially from the point of reference threshold in the intercept term than in the slope (Table **S7**), as with the baseline model.

**Figure 3.**
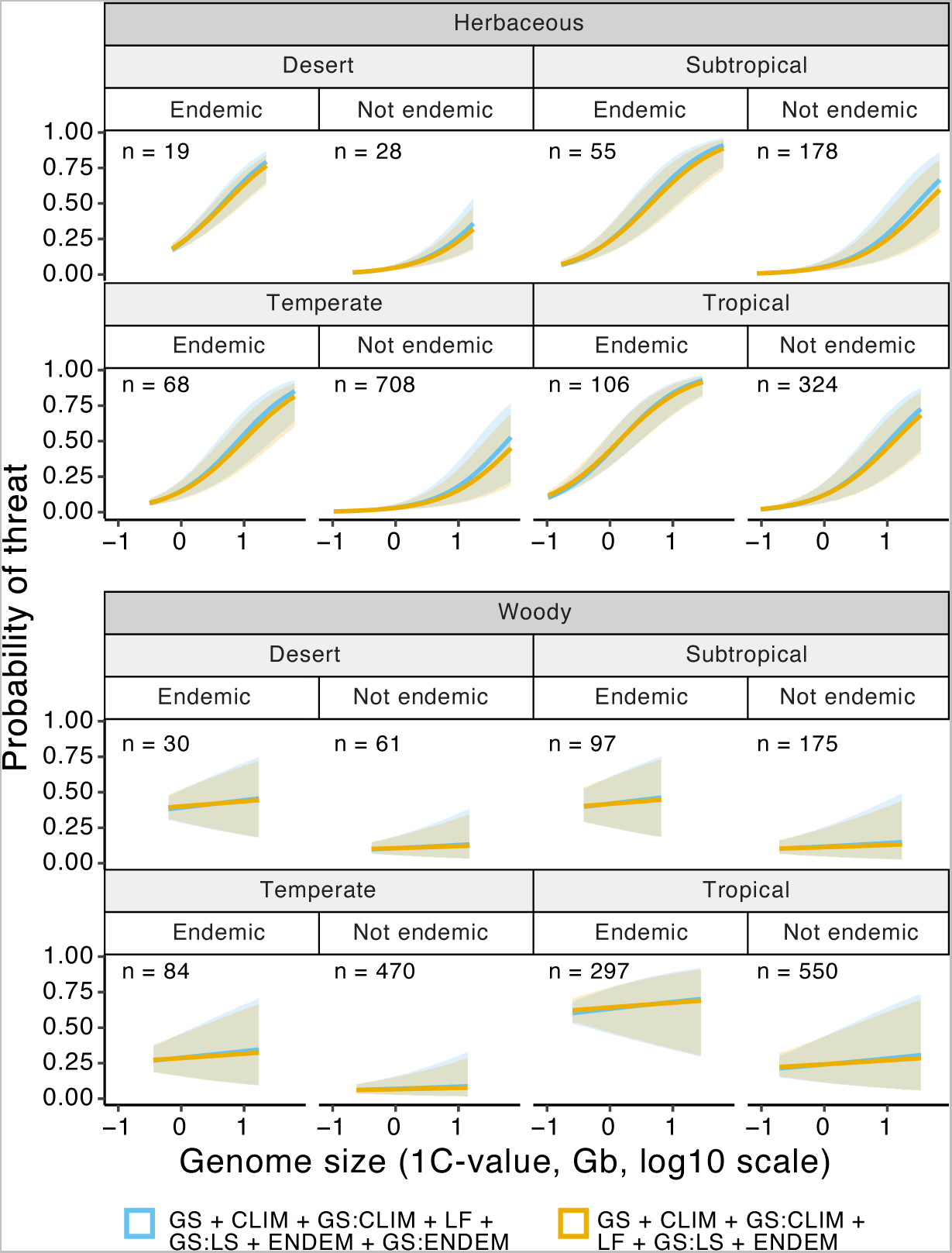
Phylogenetic logistic regression curves predicting the probability of threat in angiosperms as a function of genome size, life form, climate zone and endemism in the two best models selected when using the Red List definition of threat as a point of reference for grouping Least Concern and Near Threatened species as non-threatened and Vulnerable, Endangered, Critically Endangered and Extinct in the Wild species as threatened (see Fig. 2). Curves represent mean coefficients from phylogenetic logistic regressions, summarized across the 100 different phylogenies used as input in individual regressions conducted for each of the two best models (the latter are shown in blue and orange). Shaded areas indicate the mean 95% confidence intervals from bootstrap analysis for each model. The start and end points of the curves and confidence intervals indicate the minimum and maximum genome sizes represented across the different variable partitions (also given in Table **S3**). Abbreviations: n = sample size, GS = genome size, CLIM = climate zone, LF = life form, ENDEM = endemism.

**Table 3.** Mean coefficients and threat probabilities summarized across the two best models found in phylogenetic logistic regressions using the Red List definition of threat as a point of reference for dichotomizing the response variable into a non-threatened group (comprising Least Concern and Near Threatened species) and a threatened group (comprising Vulnerable, Endangered, Critically Endangered and Extinct in the Wild species), shown in Fig. **3**. Probabilities represent averaged estimates across very small-genomed (≤1.18 Gb/1C) and large-genomed (≥11.76 Gb/1C) herbaceous angiosperms sampled here, expressed as percentages. Mean 95% confidence intervals (CIs) from bootstrap analysis are shown in brackets. The Δ threat probability was calculated as the difference in the probability of threat between the two genome size categories (as depicted in Fig. **S3**). Coefficients and threat probabilities for three comparison threat thresholds are summarized in Table **S7**.

|  |  | Probability of threat |  |  |
| --- | --- | --- | --- | --- |
| Endemism/<br>Climate | Coefficient<br>estimates | Very small<br>genome<br>(95% CIs) | Large<br>genome<br>(95% CIs) | $\Delta$<br>threat<br>probability |
| <u>Endemic</u> |  |  |  |  |
| Desert | 1.86 | 21.0 (17.6–25.8) | 73.2 (59.2–81.9) | 52.2 |
| Subtropical | 1.89 | 14.8 (10.4–21.9) | 81.3 (62.8–90.6) | 66.4 |
| Temperate | 1.84 | 10.8 (7.1–16.4) | 71.2 (48.8–84.2) | 60.4 |
| Tropical | 1.23 | 26.0 (20.3–34.2) | 89.0 (77.1–94.4) | 62.9 |
| <u>Not-endemic</u> |  |  |  |  |
| Desert | 1.68 | 3.0 (2.2–4.5) | 30.8 (16.5–47.0) | 27.8 |
| Subtropical | 1.71 | 2.6 (1.4–4.9) | 46.2 (21.3–69.3) | 43.6 |
| Temperate | 1.66 | 1.6 (0.8–2.9) | 33.0 (13.1–56.0) | 31.4 |
| Tropical | 1.05 | 6.2 (3.7–10.9) | 61.0 (34.8–79.7) | 54.7 |

### Direct and indirect effects of genome size on extinction risk

Considering that the four varying thresholds for dichotomizing extinction risk produced consistent results across logistic regressions, we focused on the point of reference threshold to test four causal models in confirmatory phylogenetic path analyses (Fig. **S4**). We found one best causal model for subtropical and tropical species, and two best models for temperate species (Fig. **S23**). We omitted findings for desert species as these analyses did not converge (likely due to a small sample size; Table **S3**) and were therefore unreliable. Causal model one was common to all three of the reported climates; it included a direct link between genome size and extinction risk, in addition to indirect links through life form and endemism (Figs. **4**, **S4a**). Causal model three, exclusive to temperate species, differed by lacking a life form-extinction risk link (Figs. S4c, S23, S24).

**Figure 4.**
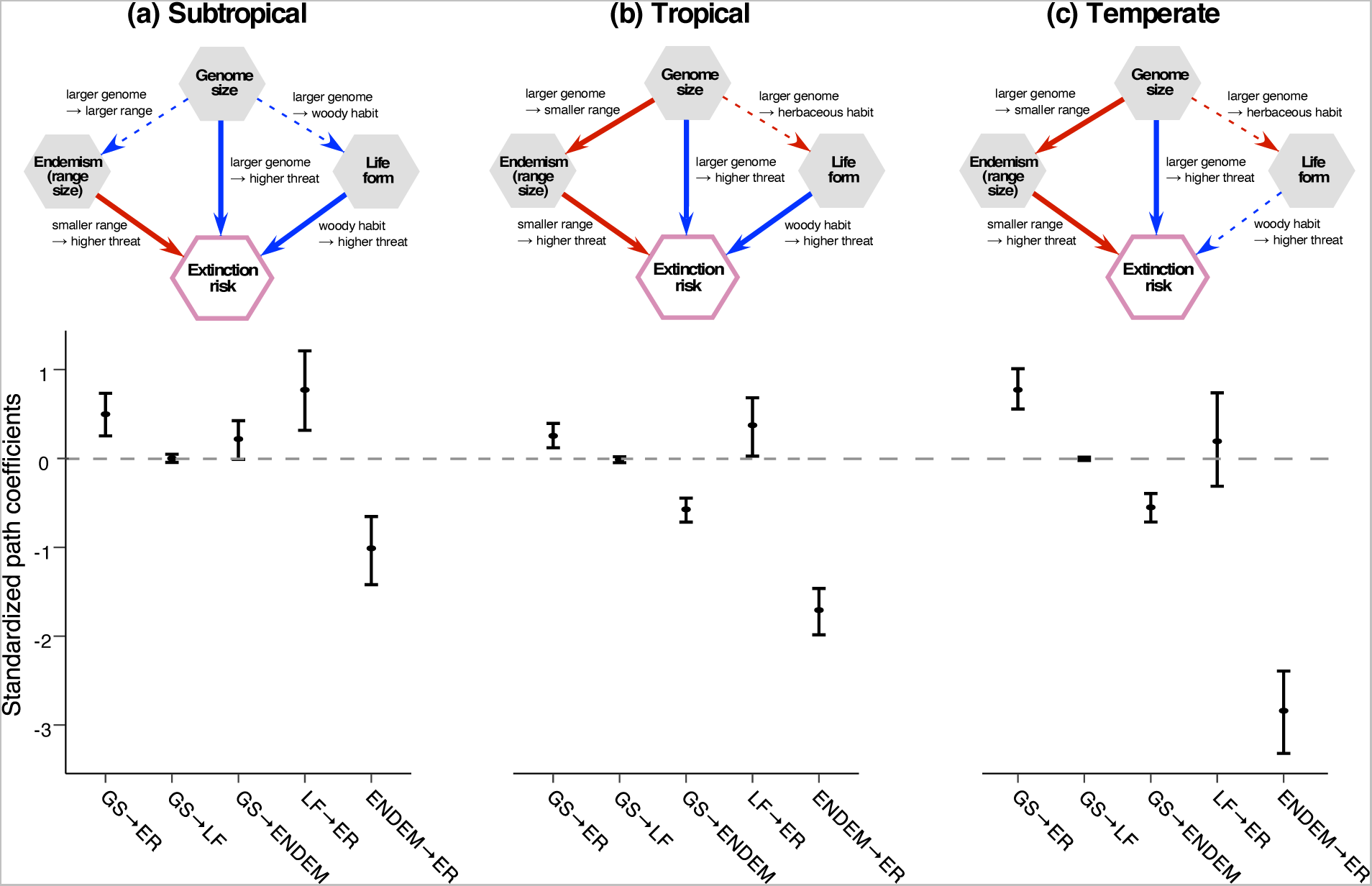
Schematics and coefficients for the best causal model obtained from confirmatory phylogenetic path analyses for **(a)** subtropical, **(b)** tropical and **(c)** temperate species using the Red List definition of threat as a point of reference for grouping Least Concern and Near Threatened species as non-threatened, and Vulnerable, Endangered, Critically Endangered and Extinct in the Wild species as threatened (desert species are excluded as analyses for this data subset did not converge). The schematics represent causal model one (shown in Fig. **S4a**), the single best model found for subtropical and tropical species based on ΔCICc model selection (Fig. **S23**); for the additional best model found for temperate species (shown in Fig. **S4c**), see Fig. **S24**. Positive links between variables are indicated by blue arrows and negative links by red arrows; solid and dashed arrows indicate significant and non-significant links, respectively. Plots show the mean standardized path coefficients and 95% confidence intervals (the latter obtained from bootstrap analyses) for links present in the models above, after summarising across the 100 different phylogenies used as input in individual path analyses for each climate. Confidence intervals fully above or below the dashed line (coefficient = 0) indicate significant coefficients. Abbreviations: GS = genome size, ER = extinction risk, LF = life form, ENDEM = endemism (proxy for range size).

All best causal models showed that increases in genome size were significantly associated with higher extinction risk across climate zones (Figs. **4**, **S24**). This link was strongest in temperate species (coefficients = 0.772 and 0.764 for causal models one and three, respectively), followed by subtropical (coefficient = 0.498) and tropical (coefficient = 0.255) species (Figs. **4, S24**). Genome size increases were also significantly associated with endemism (i.e., smaller range sizes) in tropical and temperate species (Fig. **4b**-**c**), but not in subtropical species (Fig. **4a**). Nevertheless, in all three climate zones, single-country endemics were significantly associated with increased extinction risk. This endemism-risk link was the most influential across climates and best causal models, with coefficients ranging from −1.01 to −2.839 (Figs. **4**, **S24**). The genome size-life form link was not significant across the best models (Figs. **4**, **S24**). Although we found significant links between life form and risk for subtropical and tropical species, which may merit further exploration, these fall outside of the genome size-related hypotheses tested here.

### Species with a potentially heightened risk of extinction

As genome size is positively correlated with extinction risk in herbaceous angiosperms (Figs. **3, S20**–**S22**), we filtered the 11,764 sampled species that lacked a Red List assessment to identify 816 large-genomed herbs (≥11.76 Gb/1C; Fig. **5**). This group is expected to contain a higher proportion of threatened than non-threatened species based on their genome size and life form. Of the 816 species identified (Table **S8**), 729 (89.3%) were monocots from 14 families and 87 (10.7%) were eudicots from 11 families (Fig. **5**). In comparison, monocots represent 4,484 (38.1%) of the 11,764 unassessed species in our sampling and 806 (85.6%) of the 942 unassessed large-genomed species, while eudicots represent 7,189 (61.1%) and 132 (14%) of these species, respectively (Table **S1**). Of the large-genomed herbs identified, 231 (28.3%) are single-country endemics (Fig. **5**).

**Figure 5.**
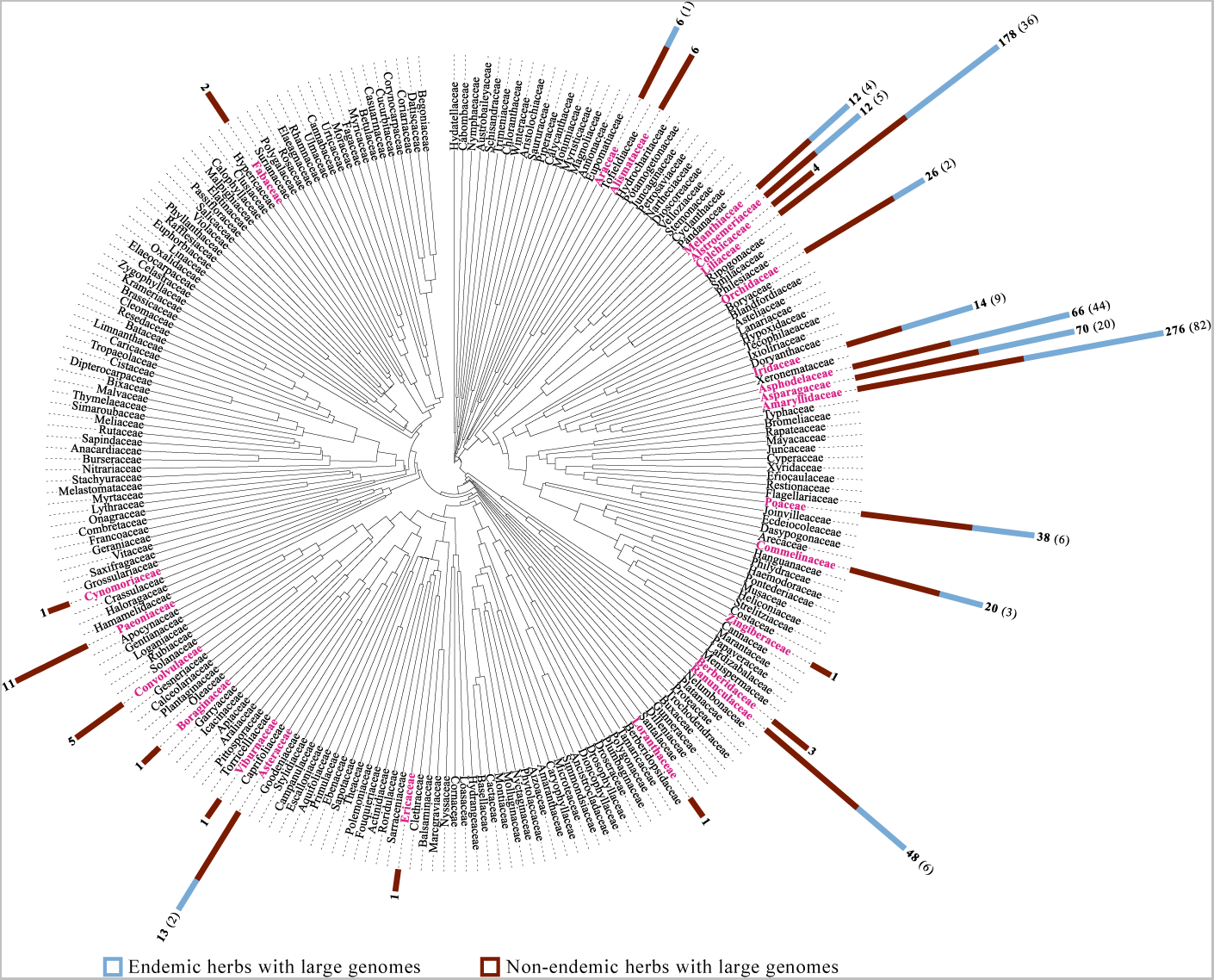
Summary phylogeny of 237 angiosperm families that contain species for which genome size information is available, but a Red List threat assessment is lacking. A randomly selected phylogeny is depicted (out of the 100 different species-level trees used in analyses), with a single terminal per family. Pink terminals indicate priority families that contain herbaceous species with large (i.e., ≥11.76 Gb/1C) genomes. The total number of unassessed large-genomed herbs in these families is represented by the full length of the bar graphs (red and blue areas) and indicated in bold font; the blue area of each bar and the values in brackets indicate the proportion of large-genomed herbs that are endemic. Note that species numbers are converted using a log10 scale for projection as bar graphs. For the full list of 816 species in these priority families see Table **S8**.

## Discussion

### Genome size and extinction risk are positively correlated across angiosperms

Our findings from phylogenetically-informed logistic regressions and path analyses provide unambiguous support for our hypothesis that genome size is positively correlated with extinction risk in angiosperms (Figs. **1**, 4). This is consistent with Vinogradov’s (2003) report of significantly larger mean genome sizes in globally threatened species than in those not assessed as threatened. Our study and that of Vinogradov are congruent despite differences in their overall approaches, notably including the underlying threat data. Here we used the Red List, which comprises 54,263 assessments and employs quantitative criteria (IUCN, 2022), whereas Vinogradov (2003) used the UNEP-WCMC Species Database that predated the 1994 Red List (fide Walter & Gillett, 1998) and instead relied on qualitative criteria to assess ∼34,000 species. Our use of the Red List uncovered that genome size does not increase gradually across categories denoting increasing risk; instead, it displays a binary pattern with significantly smaller genomes in LC compared to NT, EN and CR species (Fig. **1a**). The point where genome size begins playing a significant role in influencing extinction risk is unclear, but likely lies within the size ranges encountered in NT and VU species (Fig. **1a**). We captured this uncertainty by using four varying threat thresholds in logistic regressions, which all produced consistent results (Figs. **1b**, **3**, **S20**–**S22**), suggesting that the broad categorization of extinction risk applied here and in Vinogradov (2003) accurately captures its association with genome size.

Large genome size may thus represent a biological attribute associated with an increasing intrinsic susceptibility to extinction that additionally interacts with a range of (anthropogenic) threats to shape species risk (Vazquez & Lucifora, 2023). The effects of this intrinsic susceptibility were previously described in the “large genome constraint” hypothesis proposed by Knight *et al*. (2005), who found tentative support for evolutionary, biogeographic and phenotypic correlates of genome size that may contribute to large-genomed species being “trimmed” from the angiosperm tree of life. Evolutionary correlates included significantly lower species richness (measured as number of species per genus), underpinned by the confounding effects of higher extinction and/or lower speciation rates, but nevertheless indicative of costs imposed by large genomes over evolutionary time and not as an exclusively recent occurrence (Vinogradov, 2003). Biogeographic correlates are reflected in more restricted range sizes and ecological distributions of large-genomed species (e.g., Knight & Ackerly, 2002; Pandit *et al*., 2014; Bureš *et al*., 2022). The narrower environmental tolerances documented for large genomes are likely shaped by diverse phenotypic correlates, including a lower variation in cell sizes and packing densities, which have been shown to limit metabolic variation and therefore the ability to optimize performance across a range of environments (e.g., Roddy *et al*., 2020). Using range size as a proxy for effective population size (e.g., Gaston, 2003), the “mutational hazard hypothesis” proposed by Lynch and Conery (2003) may provide an underlying mechanism for the relatively restricted ranges of large-genomed species, whereby mutation frequency increases alongside genome size due to a higher availability of target DNA. Despite posing a selective cost from mutational hazards, large genomes may become fixed in small populations due to the prevalence of genetic drift over natural selection, potentially increasing extinction risk over time. However, more direct measures of population size (e.g., census data) are required for confirming this hypothesis, as support for it has been equivocal in angiosperms (Bureš *et al*., 2022) or conflicting in animals (e.g., Yi & Streelman, 2005; Roddy *et al*., 2021; Pincheira-Donoso *et al*., 2023).

These genome size correlates, in conjunction with angiosperms generally having smaller genomes than expected given the ubiquity of repetitive DNA and polyploidy (e.g., Wang *et al*., 2021), provide persuasive arguments for the potentially maladaptive consequences of large genomes (Vinogradov, 2003; Knight *et al*., 2005). However, genome size alone does not fully explain extinction risk, as illustrated by the presence of both threatened and non-threatened species with large genomes in our sampling (Table **S3**). Moreover, the contrasting responses to environmental stress conditions documented for large-genomed angiosperms across different lineages and growth forms (e.g., Faizullah *et al*., 2021; Feng *et al*., 2022; Zhang *et al*., 2022) make it difficult to provide a single explanation for why species with large genomes are more likely to be threatened.

### The relationship between genome size and extinction risk varies across life forms

Our second main finding is that the genome size and extinction risk relationship is driven by a signal in herbaceous, but not woody species (Fig. **3**). Despite woody angiosperms having generally smaller genomes and lower overall size variance compared to herbs (Table **S3**; Carta *et al.,* 2022), our dataset contains enough genome size variation in woody species that we would expect to detect a relationship with extinction risk if indeed it exists. Rather, the lack of signal in woody species may be associated with the scaling effects of cell size, as small genomes are typically associated with small stomata that can close rapidly, a potential advantage in tall plants for permitting greater conductance through long xylem pathways while reducing hydraulic dysfunction, particularly under drought conditions (e.g., Hetherington & Woodward, 2003; Beaulieu *et al*., 2010). Thus, large genomes are predicted to be removed by strong selection in trees (Beaulieu *et al*., 2010), enabled by their typically large effective population sizes (Petit & Hampe, 2006), as predicted by the mutational hazard hypothesis (Lynch & Conery, 2003). Selective pressures may be reinforced by the generally long generation times of woody species and relatively uniform environments experienced over their lifetime, two factors that are expected to reduce dynamism in genome evolution (Levin & Wilson, 1976). These forces could in turn constrain genome growth in woody species to a point where genome size does not increase extinction risk, which is instead driven by extrinsic factors such as habitat loss or degradation (Newton & Oldfield, 2008). In contrast, genetic drift rather than selection may shape genome size dynamics in the generally smaller population sizes of herbaceous angiosperms, which are additionally likelier to experience a range of variable local environments (Leitch & Leitch, 2012).

Comparing our findings for woody angiosperms with gymnosperms (the other major vascular plant lineage containing woody species) may be of interest. However, we expect this to provide limited insights to support or refute our hypotheses given the contrasts between these two lineages concerning genome dynamics and profiles, including genome size ranges, chromosome numbers and polyploidy frequency (Leitch & Leitch, 2012).

### Genome size is associated with extinction risk both directly and indirectly via range size

We found that genome size and range size (using endemism as a proxy) are interwoven in their effects on extinction risk. First, path analyses showed that some of the effect of range size on extinction risk can be indirectly attributed to genome size (Fig. **4**). This is somewhat expected given (i) the generally small range sizes of large-genomed species compared to the diverse range sizes of small-genomed species (Bureš *et al*., 2022); and (ii) the fact that range size underpins Red List criterion B, designed to identify risk in populations with restricted distributions (in combination with additional population-level metrics; IUCN, 2013). Second, logistic regressions showed that genome size has a stronger effect on the extinction risk of species that are endemic to a single botanical country than non-endemic species (Fig. **3**). In-silico modelling provides a potential mechanism for this finding, whereby genome expansion drives extinction risk in small populations by increasing the lethal mutational burden (LaBar & Adami, 2020). Restricted ranges may also interact with major anthropogenic impacts like land conversion and species overexploitation to further exacerbate risk.

Path analyses showed that a large genome size, beyond its indirect link via range size, additionally has direct and significant effects on angiosperm extinction risk, likely underpinned by the nuclear-, cellular-, and organism-level constraints imposed by large genomes (Fig. **4**). Although data for these constraints are not available at the scale of this study and therefore were not explicitly included in our models, our findings highlight the possibility that genome size may be a useful proxy for parameters that are difficult to measure but associated with intrinsic risk in angiosperms and aligned with Red List criteria. These parameters may include maximum photosynthetic rate, water-use efficiency, and nutrient demand (Guignard *et al*., 2016; Faizullah *et al*., 2021; Schley *et al*., 2022).

### Genome size influences extinction risk relatively uniformly across climates

In contrast to life form and range size, we found weaker support for climatic heterogeneity in the genome size and extinction risk relationship. In both logistic regressions and path analyses (Figs. **3**, 4), the effect of genome size was smallest in tropical species. Other studies documented on average smaller genomes for tropical than temperate species (e.g., Levin & Funderburg, 1979; Bureš *et al*., 2022), perhaps arising from selection against larger genomes that have a competitive disadvantage in environments requiring rapid growth to trap sunlight due to slower rates of cell division. The advantages of smaller genomes may therefore partly explain our findings of a more limited role for genome size in the tropics, where other factors, such as differences in the global distribution of threat types, are likely to influence extinction risk. For example, the main threats documented in the Red List for terrestrial vertebrates differed between tropical areas, where agriculture and logging are more pervasive, compared to temperate areas, where pollution and invasive species are the dominant documented threats (Harfoot *et al*., 2021). Whether these patterns accurately reflect the relative strength of threats globally, or bias in how threat is recorded in the Red List, they may contribute to explaining why genome size-related threats may appear less important in the tropics when facing swift anthropogenic drivers of extinction like land conversion, which would be only indirectly associated with genome size through its effects on range size (Figs. **3**, **4**). In contrast, intrinsic factors like genome size may underlie our findings of relatively larger effect sizes in temperate species (Fig. **4**), where the cascading effects of large genomes may constrain plant responses to threats like pollution (e.g., Temsch *et al*., 2010).

### Implications for conservation

Our main finding of concurrent increases in genome size and extinction risk in herbaceous angiosperms (Figs. **1**, **3**, **4**), has both theoretical and practical implications for conservation. In addition to guiding fundamental research to understand the underlying causes of differential extinction risk, it may also prove useful for informing decisions relevant to plant conservation. For example, we identified 816 herbaceous species lacking a Red List assessment but belonging to a pool likely containing a higher proportion of at-risk species based on their large genomes and life form (Fig. **5**). Considering genome size in the context of predictors with high importance for estimating plant extinction risk, such as range size (Pelletier *et al*., 2018; Walker *et al*., 2022), may prove useful for understanding the relative risk of species with similar geographic and ecological characteristics but dissimilar genome size. Additionally, it may serve as motivation for addressing underrepresentation in the Red List of herbaceous angiosperms (Table **S4**), particularly monocots (Nic Lughadha *et al*., 2020), a group with an exceptionally high number of large-genomed herbs (Fig. **5**; Pellicer *et al*., 2018). Conversely, genome size may be useful information for fast tracking the identification of LC species (e.g., Bachman *et al*., 2020), considering that small-genomed herbs have a 15–43% lower threat probability than large-genomed ones (Table **3**, Fig. **3**). Extending the number of species with both genome size data and a Red List assessment, while maximising taxonomic and geographic representativity, will enhance our understanding of the role of genome size in extinction risk and its potential for informing conservation strategies. Additionally, future studies characterizing how the impacts of genome size vary across different threat types may help to explicitly model the influence of anthropogenic activities on angiosperm extinction risk.

## Acknowledgements

The authors thank Marie Henniges for assistance with phylopath analyses, Cecile Ané for advice on phyloglm, Barbara Holland and Ben Halliwell for their suggestions on phylogenetic comparative methods, and Barnaby Walker for providing scripts to standardise life form data. This research was supported by a Future Leader Fellowship (RBG, Kew) to M. Soto Gomez; J. Pellicer benefited from a Ramón y Cajal grant (Ref: RYC-2017-2274) funded by both MCIN/AEI/ 10.13039/501100011033 and “ESF Investing in your future”.

## Author contributions

M.S.G, I.L., E.N.L. and M.J.M.B. designed the research. P.V., P.B, T.L.E. and F.Z. provided genome size and life form data. F.F. provided angiosperm phylogenies for analyses. M.S.G and M.J.M.B. performed data analysis, interpretation and visualization with input from S.P and E.N.L. M.S.G. led the writing of the manuscript with substantial input from I.L., E.N.L., J.P. All authors participated in manuscript writing and editing.

## Data availability statement

The dataset assembled for this study is available in the supplementary materials (Table S1).

## Supporting Information

The following Supporting Information is available for this article:

**Supplementary Methods S1.** Dataset assembly and taxonomic reconciliation with the World Checklist of Vascular Plants.

**Supplementary Methods S2.** Adjustments to life form, climate zone and geographic information.

**Supplementary Methods S3.** Preparation of angiosperm phylogenies

**Supplementary Methods S4.** Characterization of species representativeness in the genome size dataset.

**Supplementary Methods S5.** Exploration of likelihood space for a baseline phylogenetic logistic regression model.

**Supplementary Methods S6.** Selection of best models from phylogenetic logistic regressions.

**Supplementary Methods S7.** Direct and indirect effects of genome size on extinction risk

**Table S1** (separate Excel table). Complete dataset used across analyses, comprising angiosperm genome size, life form, climatic zone and endemism status (proxy for range size).

**Table S2** (separate Word table). Summary of sample size in individual Red List categories across data partitions in the 3,250-species dataset.

**Table S3** (separate Word table). Summary statistics on sample size and genome size for the 3,250-species dataset.

**Table S4** (separate Word table). Evaluation of species representativeness across data partitions in the 3,250-species dataset.

**Table S5** (separate Word table). Summary of logistic regression models, fit and phylogenetic signal across varying threat thresholds for dichotomizing extinction risk.

**Table S6** (separate Word table). Consensus coefficients for best models identified in phylogenetic logistic regressions using varying threat thresholds for dichotomizing extinction risk.

**Table S7** (separate Excel table). Mean threat probabilities summarized across best models from phylogenetic logistic regressions using varying threat thresholds to dichotomize extinction risk.

**Table S8** (separate Word table). List of candidate species for prioritizing threat assessments.

**Figure S1.** Comparison of point-based and binary representations of species ranges.

**Figure S2.** Schematic of the model selection pipeline applied to phylogenetic logistic regressions.

**Figure S3.** Schematic of the methods used to estimate a mean probability of threat for very small- and large-genomed herbaceous species across best models from logistic regressions.

**Figure S4.** Competing causal models tested in confirmatory phylogenetic path analyses.

**Figure S5.** Phylogenetic distribution of the genera represented in the 3,250-species dataset.

**Figure S6.** Distribution of genome sizes in the 3,250-species dataset and a larger sampling of angiosperms.

**Figure S7.** Distribution of mean genome size differences in the 3,250-species dataset and 999 down-sampled subsets with equal proportions of threatened and non-threatened species.

**Figure S8.** Model selection for phylogenetic logistic regressions using a higher threat threshold than the Red List to dichotomize extinction risk.

**Figure S9.** Model selection for phylogenetic logistic regressions using a polarizing threat threshold (by excluding Near Threatened species) to dichotomize extinction risk.

**Figure S10.** Model selection for phylogenetic logistic regressions using a lower threat threshold than the Red List to dichotomize extinction risk.

**Figure S11. C**oefficients distribution for one of two best models from phylogenetic logistic regressions using the Red List definition of threatened to dichotomize extinction risk.

**Figure S12.** Coefficients distribution for a second of two best models from phylogenetic logistic regressions using the Red List definition of threatened to dichotomize extinction risk.

**Figure S13.** Coefficients distribution for the best model from phylogenetic logistic regressions using a lower threat threshold than the Red List to dichotomize extinction risk.

**Figure S14.** Coefficients distribution for one of three best models from phylogenetic logistic regressions using a higher threat threshold than the Red List to dichotomize extinction risk.

**Figure S15.** Coefficients distribution for a second of three best models from phylogenetic logistic regressions using a higher threat threshold than the Red List to dichotomize extinction risk.

**Figure S16.** Coefficients distribution for a third of three best models from phylogenetic logistic regressions using a higher threat threshold than the Red List to dichotomize extinction risk.

**Figure S17.** Coefficients distribution for one of three best models from phylogenetic logistic regressions using a polarizing threat threshold (omitting Near Threatened species) to dichotomize extinction risk.

**Figure S18.** Coefficients distribution for a second of three best models from phylogenetic logistic regressions using a polarizing threat threshold (omitting Near Threatened species) to dichotomize extinction risk.

**Figure S19.** Coefficients distribution for a third of three best models from phylogenetic logistic regressions using a polarizing threat threshold (omitting Near Threatened species) to dichotomize extinction risk.

**Figure S20.** Curves predicting threat probability in a best model common to phylogenetic logistic regressions using a lower, higher and polarizing threshold to dichotomize extinction risk.

**Figure S21.** Curves predicting threat probability in a best model common to phylogenetic logistic regressions using a higher, polarizing and point of reference threshold to dichotomize extinction risk.

**Figure S22.** Curves predicting threat probability in a best model common to phylogenetic logistic regressions using a higher and polarizing threshold to dichotomize extinction risk.

**Figure S23.** Support for competing causal models tested in confirmatory phylogenetic path analyses across different climates using the Red List definition of threatened to dichotomize extinction risk.

**Figure S24.** Schematic and coefficients for one of two best causal models for temperate species using the Red List definition of threatened to dichotomize extinction risk.

