## Supplementary Materials for "Genome size is positively correlated with extinction risk in herbaceous angiosperms"

**Table S8** (separate Word table). List of candidate species for prioritizing threat assessments.

**Supplementary Methods**

**Methods S1**

*Dataset assembly and taxonomic reconciliation with the World Checklist of Vascular Plants*

We used the World Checklist of Vascular Plants (WCVP; Govaerts *et al.*, 2021) as the taxonomic basis for reconciling sample names and taxonomic ranks across the various data sources employed in this study. Specifically, we collated (i) genome size data from the Plant DNA C-values Database (Leitch *et al.*, 2019) and a newly published dataset (Bureš *et al.*, 2022); (ii) global extinction risk data from the Red List (IUCN, 2022); (iii) life form, climate zone, and distribution data from WCVP; and (iv) phylogenetic information from a sampling of 100 species-level angiosperm phylogenies (Forest, 2023).

We first determined the taxonomic status of samples in the two source datasets for genome size, confirming whether these had names accepted in WCVP or non-accepted names that required reconciliation. To do so, we used the R package rWCVP (Brown *et al.*, 2023), with the full taxon names in the genome size datasets as input, including author information when available. We assembled an initial genome size dataset comprising 11,278 taxa from the Plant DNA C-values database (Leitch *et al.*, 2019); we then used the “wcvp_accepted_plant_name_id” information provided by rWCVP to identify and add 7,261 taxa from Bureš *et al.* (2022) that were absent from the initial dataset. We then used the “match_type” output from rWCVP to identify 16,565 taxa with identical or nearly identical matches to accepted taxon names in the WCVP, considering taxon authority information when it was available. The “match types” that rWCVP produced for these taxa were “matched in WCVP with author” or “matched in WCVP without author” for identical matches; nearly identical matches were recorded as “phonetically matched (strong)” or “fuzzy matched (strong)”. We retained these taxon names without modification for species. However, for infraspecific taxa (which represented subspecies, varieties, nothovarieties and nothospecies) we replaced their “wcvp_accepted_plant_name_id” with the “parent_plant_name_id” provided by rWCVP, effectively raising these samples to the species rank. Working at the species rather than infraspecific level allowed us to maximize the number of overlapping taxa across the various data sources employed here for genome size, extinction risk, life form, climate zone, endemism, and phylogenetic information.

We individually assessed the remaining 1,974 taxon names in the initial genome size dataset, as the “match types” from rWCVP for these were “fuzzy matched (weak)”, “phonetically matched (weak)”, “multiple fuzzy matches found”, “multiple matches found”, “no fuzzy match with similarity >0.75 found”, or “no phonetic match with similarity >0.75 found”. We retained 551 taxa that we were able to confidently assign to accepted species in WCVP. For taxa with multiple matches (resulting from synonymy or fuzzy matching), in many cases we were able to select a single name by visually matching the taxon author information from the original datasets with that of the accepted name in the WCVP; author information was not matched by rWCVP in these cases due to substantial spelling or abbreviation differences. When taxon author information was unavailable, we either selected the WCVP accepted name that exactly matched the original name (if such a match was recovered by rWCVP), or excluded the species from further consideration if there was insufficient information to choose amongst the multiple matches.

The provisional 17,116-taxon dataset obtained from taxonomic reconciliation with rWCVP underwent further filtering to (i) retain a single placeholder for species represented by multiple accessions (by retaining the accession with the smallest genome size; see main text), and (ii) remove species not found in the 100 phylogenies that we used as input in analyses. This resulted in a final 15,158-taxon dataset that included the original and WCVP-accepted name for all species (Table **S1**). Finally, we used the “accepted plant name id” output from rWCVP to link taxa in the genome size dataset to available data for that taxon in the WCVP, Red List, and angiosperm phylogenies employed here; the latter two datasets were also taxonomically reconciled to WCVP (see Brown *et al.,* 2023, for the Red List and Forest, 2023, for the phylogenies).

**Methods S2**

*Manual adjustments to life form, climate zone and geographic information*

We standardized life form, climate zone and geographic information to facilitate incorporation of these data in the statistical analyses conducted here. We used WCVP as a source of data for the three variables, additionally obtaining life form information from Bureš *et al.* (2022) when unavailable in WCVP. The original life form classifications follow the system of Raunkiær (1934) with species in the genome size dataset representing 204 different categories that we rescored as either “herbaceous” or “woody” by following the methods of Humphreys *et al.* (2019), with the following modifications. We first used the standardization developed in that study to rescore species as “annual”, “epiphytic”, “herbaceous perennial, or “woody perennial”. We further simplified these four categories into two after manual curation and rescoring (i) annual species as herbaceous (except for three out of 1,939 species that we rescored as woody); (ii) epiphytic species as herbaceous (except for 87 out of 1,005 species that we rescored as woody); (iii) herbaceous perennials as herbaceous (except for one out of 7,174 species that we rescored as woody); and all 5,023 woody perennials as woody. Life form information was not available for the remaining 17 species in the dataset. The original life form information and final dichotomization are provided in Table **S1** for all species.

Species in the genome size dataset represented nine different climate categories that we aggregated into four groupings corresponding to desert areas (with two categories rescored: desert and dry shrubland, desert or dry shrubland), subtropical areas (with one category rescored: subtropical), tropical areas (with four categories rescored: wet tropical, seasonally dry tropical, montane tropical, subtropical and tropical), and temperate areas (with one category rescored: temperate). We excluded species categorized as subalpine or subarctic from analyses due to the low availability of Red List assessments for this climate subset (n = 16). The original and final climate categories are provided in Table **S1** for all species.

The species geographic information that we obtained from WCVP corresponds to Level 3 of the TDWG World Geographic Scheme for Recording Plant Distributions (Brummitt *et al.*, 2001). We scored species as endemic if their native distribution is restricted to a single botanical country and the remaining species were scored as non-endemic. This binary coding was strongly correlated with extent of occurrence (EOO) estimates (Fig. **S1**), which we obtained from Bures *et al.* (2023) for 2,750 of the 3,250 species in our sampling with available occurrence data in the Gobal Biodiversity Information Facility (GBIF). This suggests that our endemic vs. non-endemic coding constitutes a reliable proxy for range size while overcoming the lack of point data for 500 species in our sampling.

**Methods S3**

*Preparation of angiosperm phylogenies*

We adapted the species-level angiosperm phylogenies of Forest (2023) to use as input in phylogenetically-informed statistical analyses. We used all 100 of these phylogenies to capture the uncertain phylogenetic placement of 935 species (28.8%) in our 3,250-species dataset that lacked phylogenetic data. The original phylogenies comprise all 329,798 angiosperm species recognized in WCVP across 417 families and 64 orders (version 6; Govaerts *et al*., 2021). Briefly, they were assembled by Forest (2023) by first using the GBMB phylogeny of Smith and Brown (2018) as a molecular backbone that was estimated in the latter study using a variable suite of markers (obtained from GenBank) and time-calibrated using 590 secondary constraints (obtained from Magallón *et al.*, 2015). Then, species recognized by WCVP but absent from the GBMB phylogeny due to unavailable sequence data were added in 100 separate imputations using the “scenario 2” option in the R package V.PhyloMaker (Jin & Qian, 2019), thereby obtaining 100 different trees (Forest, 2023).

Here we used the keep.tip function in the R package ape v.5.6-2 (Paradis & Schliep, 2018) to prepare reduced versions of the phylogenies comprising only the 3,250 species in our dataset. We used the bifurcatr function in the R package PDcalc (Nipperess & Wilson, 2020) to resolve polytomies present in the original phylogenies. The final phylogenies were fully bifurcating, ultrametric and rescaled to a total height of 1.0 for statistical analyses (following Ho & Ané, 2014a).

**Methods S4**

*Characterization of species representativeness in the genome size dataset and testing for effects of imbalanced sampling*

We used four approaches to characterize the representativeness of the different data types contained in our sampling of 3,250 species relative to angiosperms with available information, and to test for potential effects of imbalanced representation. First, we used the *D* statistic (Fritz & Purvis, 2010), implemented in the R package caper (Orme *et al.*, 2018), to test whether the sampling in the genome size dataset was phylogenetically clustered (i.e., taxonomically biased) or overdispersed (i.e., randomly sampled) with respect to angiosperms as a whole. It was computationally intractable to obtain species-level estimates for the 329,798 angiosperms recognized by WCVP. Therefore, we applied the *D* statistic at the genus level by scoring the 993 genera represented in the genome size dataset as “1” and the remaining 12,510 angiosperm genera recognized by WCVP as “0” (for a total of 13,503 genera). We performed the calculation on all 100 of the Forest (2023) phylogenies, after pruning these to retain a single representative from each of the 13,503 angiosperm genera. The *D* statistic equals 1 if a binary trait is randomly distributed across a phylogeny and decreases towards 0 as the trait approaches a Brownian model of evolution (Fritz & Purvis, 2010).

Second, we used the 202,753 species in WCVP with available life form, climate zone and distribution information as a baseline for estimating a factor of representation for each of the 16 data partitions in our dataset, which we obtained by grouping species according to these three characteristics. We calculated factors as the proportion of species in each partition of the genome size dataset, divided by the proportion of species in the equivalent partition in WCVP. For individual partitions in the genome size dataset, a factor of 1 indicates that the same number of species would be expected in a random sample of 3,250 angiosperms; factors of 2 and 0.5 respectively indicate double and half the number of expected species.

Third, we assessed our coverage of angiosperm genome sizes documented to date by comparing the distribution in our 3,250-species dataset to that of 15,167 angiosperms with available information (i.e., the full genome size dataset assembled here before filtering for species assessed by the Red List; Table **S1**). Finally, we tested whether imbalanced proportions of non-threatened and threatened species in our sampling may have influenced our results. To do so, we estimated the mean genome size difference between non-threatened and threatened species in our dataset and compared this to the mean difference in 999 subsets of our dataset that were randomly down-sampled without replacement to equalize non-threatened and threatened proportions with 500 species per threat group. We used the Red List definition of threat for dichotomizing extinction risk in this analysis, aggregating Least Concern and Near Threatened species into a non-threatened group and Vulnerable, Endangered, Critically Endangered and Extinct in the Wild species into a threatened group (we did not perform this test on the three comparison dichotomizations of extinction risk).

**Methods S5**

*Exploration of likelihood space for a baseline phylogenetic logistic regression model*

We used a baseline model in phylogenetic logistic regressions to characterize extinction risk in angiosperms as a function of genome size, using four varying threat thresholds to dichotomize the response variable. The model using the Red List definition of threat for grouping species into a non-threatened group (by aggregating the Least Concern and Near Threatened categories) and a threatened group (by aggregating the Vulnerable, Endangered, Critically Endangered and Extinct in the Wild categories), resulted in bimodally distributed estimates across the 100 different phylogenies used as input in individual analyses, with intercepts ranging from -4.3 to -2.7 and slopes from 0.19 to 0.38. This was indicative of a likelihood surface containing multiple local optima. Therefore, we explored likelihood space thoroughly by initiating this analysis using 25 different starting points in a gridded approach spanning intercepts from -5 to -1 (at 1-unit intervals) and slopes from 0.1 to 0.5 (at 0.1-unit intervals). We then selected the model with the highest penalized log-likelihood averaged across the 100 separate runs based on different phylogenies.

**Methods S6**

*Selection of best models from phylogenetic logistic regressions*

We performed model selection using AICc, a variation of the Akaike Information Criterion that corrects for small sample sizes (Burnham & Anderson, 2002), as depicted in Fig. **S2**. For each of the 100 phylogenies across the four threat thresholds used here for dichotomizing the response variable, we obtained 27 AIC values from phyloglm, representing the distinct logistic models (Table **1**). We first converted these AIC to AICc values using the function AICc in the R package wiqid (Meredith, 2020). We then ranked the 27 models within each phylogeny by calculating ∆AICc, which is the difference between the best model for a given tree (∆AICc = 0) and the remaining 26 models for that same tree. We obtained a median ∆AICc for each model by summarizing across all 100 phylogenies for each threat threshold. Finally, we selected a set of best models after firstly discarding those with a median ∆AICc >6, and secondly those that were more complex versions of models with a lower ∆AICc (following Richards *et al.*, 2011). For the final model set, we again used phyloglm to run 2,000 bootstrap replicates (Ives & Garland, 2010), obtaining 95% confidence intervals for model variables across all phylogenies.

**Methods S7**

*Direct and indirect effects of genome size on extinction risk*

We used confirmatory phylogenetic path analyses (PPA; von Hardenberg & Gonzalez-Voyer, 2013) to test whether angiosperm extinction risk is directly or indirectly associated with genome size, life form and endemism across different climate zones. We constructed four “causal models” (i.e., directed acyclic graphs; Fig. **S4**) and applied the R package phylopath (van der Bijl, 2018) with the method “logistic_MPLE” to test these in a framework suitable for continuous (i.e., genome size) and binary (i.e., life form, endemism) data. For these analyses, we used the point of reference threat threshold to obtain non-threatened (i.e., LC, NT) and threatened (i.e., VU, EN, CR, EW) groupings. Life form was coded as herbaceous (0) and woody (1), and endemism as endemic (0) and non-endemic (1). To incorporate climate zones (the remaining variable, comprising four categories) we partitioned the data into desert, subtropical, tropical and temperate species, and analysed each subset separately. We also performed separate analyses using 100 different phylogenies as input to account for phylogenetic uncertainty, resulting in a total of 400 PPAs (i.e., 100 analyses for each of the four climate subsets). For each PPA, we used ∆CICc (C-statistic information criterion corrected for small sample sizes) to rank the four competing causal models, discarding those with ∆CICc >2 (van der Bijl, 2018). The best causal models for each PPA were then bootstrapped using 500 replicates to obtain 95% confidence intervals. Coefficients were considered to be significant if their confidence intervals excluded zero and non-significant otherwise (e.g., Guo *et al.*, 2019).

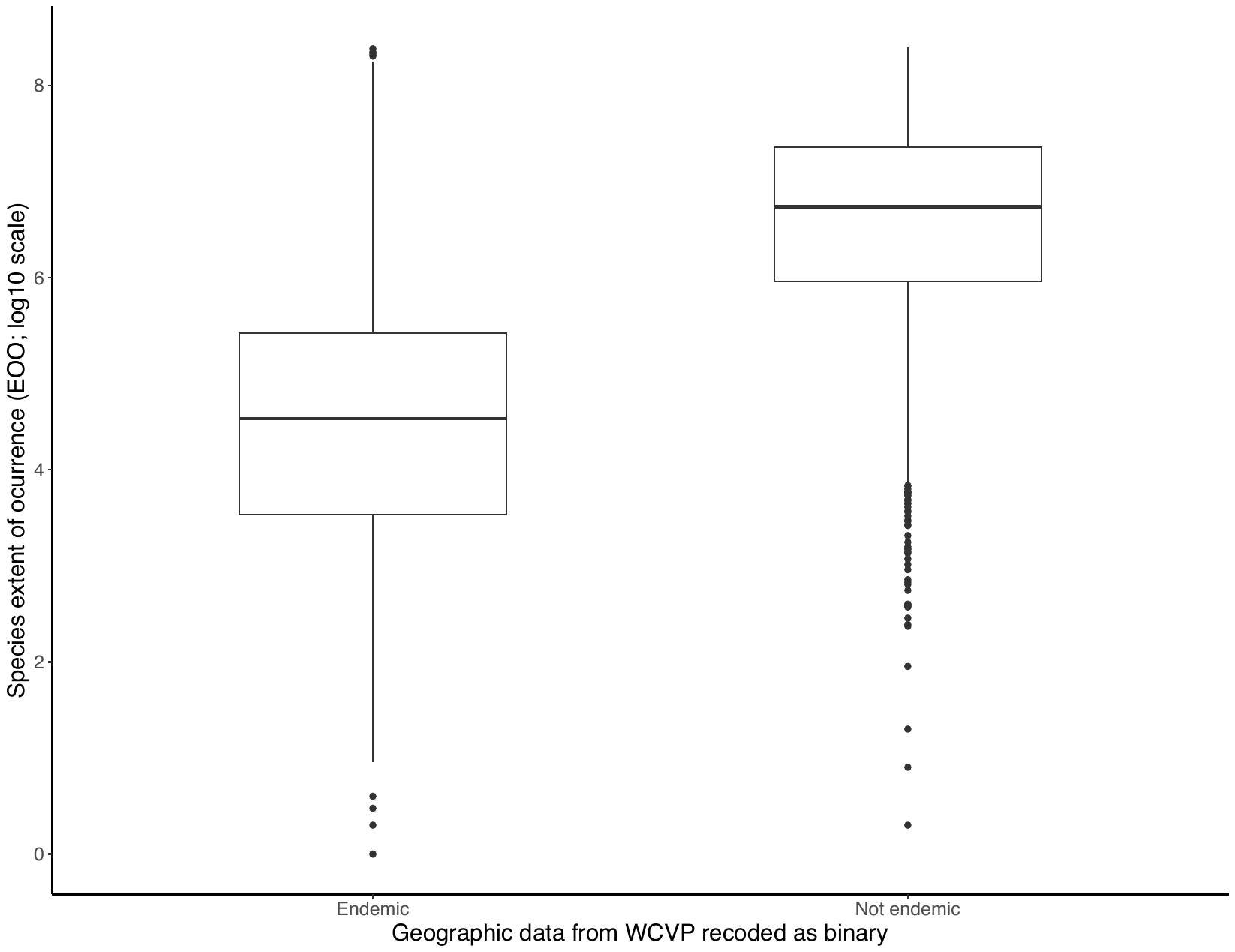

**Figure S1.** Comparison of the binarization applied here to geographic data from WCVP scoring species as being endemic to a single botanical country or non-endemic, with point-derived estimates of extent of occurrence (EOO). The comparison includes 2,750 species in the genome size dataset for which point estimates were available.

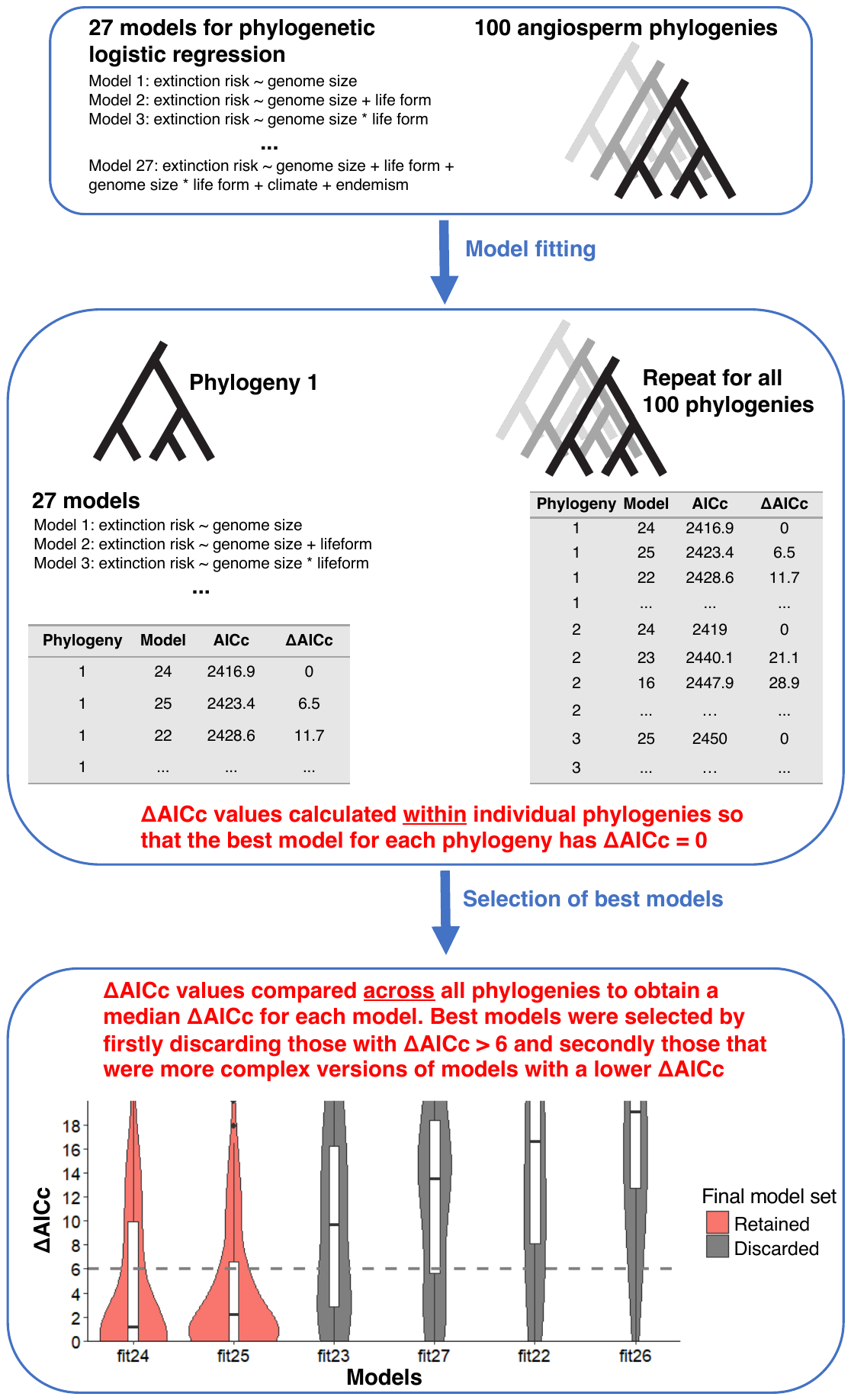

**Figure S2.** Schematic of the ΔAICc-based model selection pipeline applied to phylogenetic logistic regressions. These steps were applied individually to the four varying threat thresholds used to dichotomize the response variable.

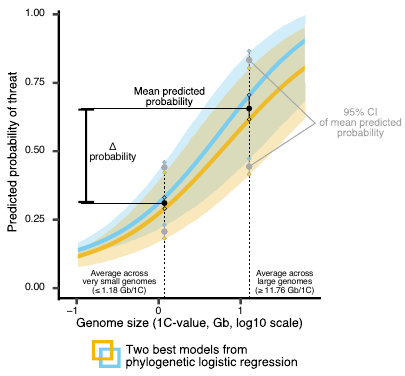

**Figure S3**. Schematic of the methods used to estimate a consensus mean probability of threat (and 95% confidence intervals) for very small- and large-genomed herbaceous species across best models identified in phylogenetic logistic regressions. This example depicts a case when two best models were found; the same procedure was applied when three best models were found. The Δ probability of threat was calculated as the difference between the averaged threat across the very small genome range vs. the large genome range. The values resulting from these calculations are shown in Tables **3 and S7**.

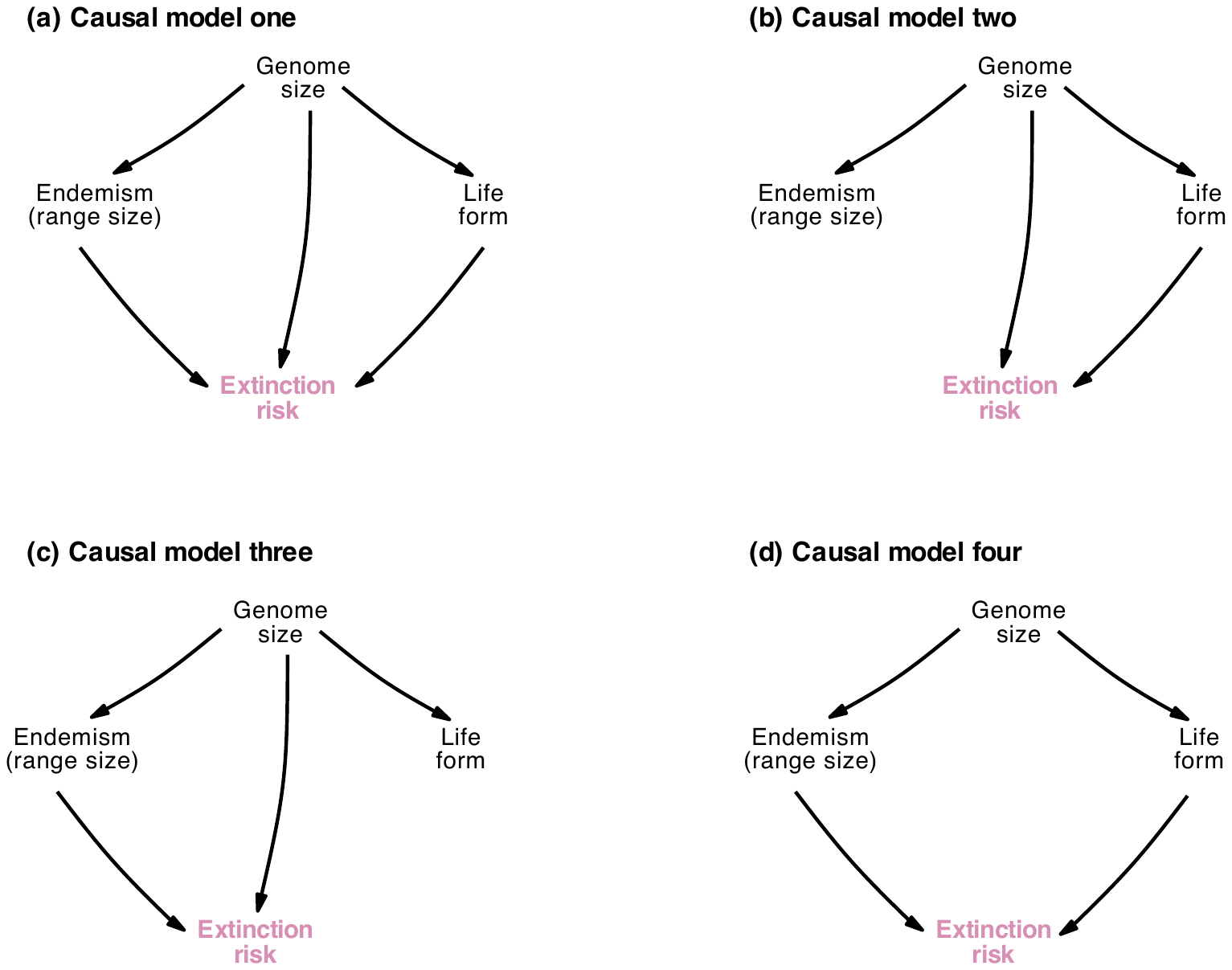

**Figure S4**. Competing causal models (i.e., directed acyclic graphs) tested using confirmatory phylogenetic path analysis. These were ranked in individual path analyses using 100 different phylogenies as input, across four separate data subsets comprising subtropical, tropical, temperate or desert species (see Figs. **4**, **S23**, **S24**).

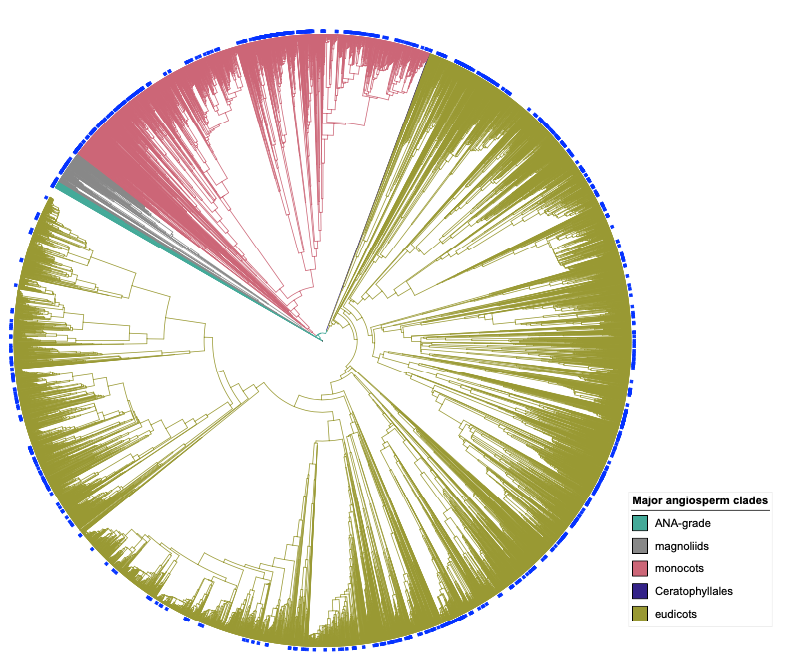

**Figure S5.** Phylogenetic distribution of the 993 angiosperm genera represented in the 3,250-species genome size dataset assembled here, shown by solid blue bars on the outside of the phylogenetic tree. A randomly selected phylogeny is depicted (out of the 100 different trees used in analyses) after pruning it to retain a single placeholder for each of the 13,503 genera it represents. Major angiosperm clades are indicated by colored branches.

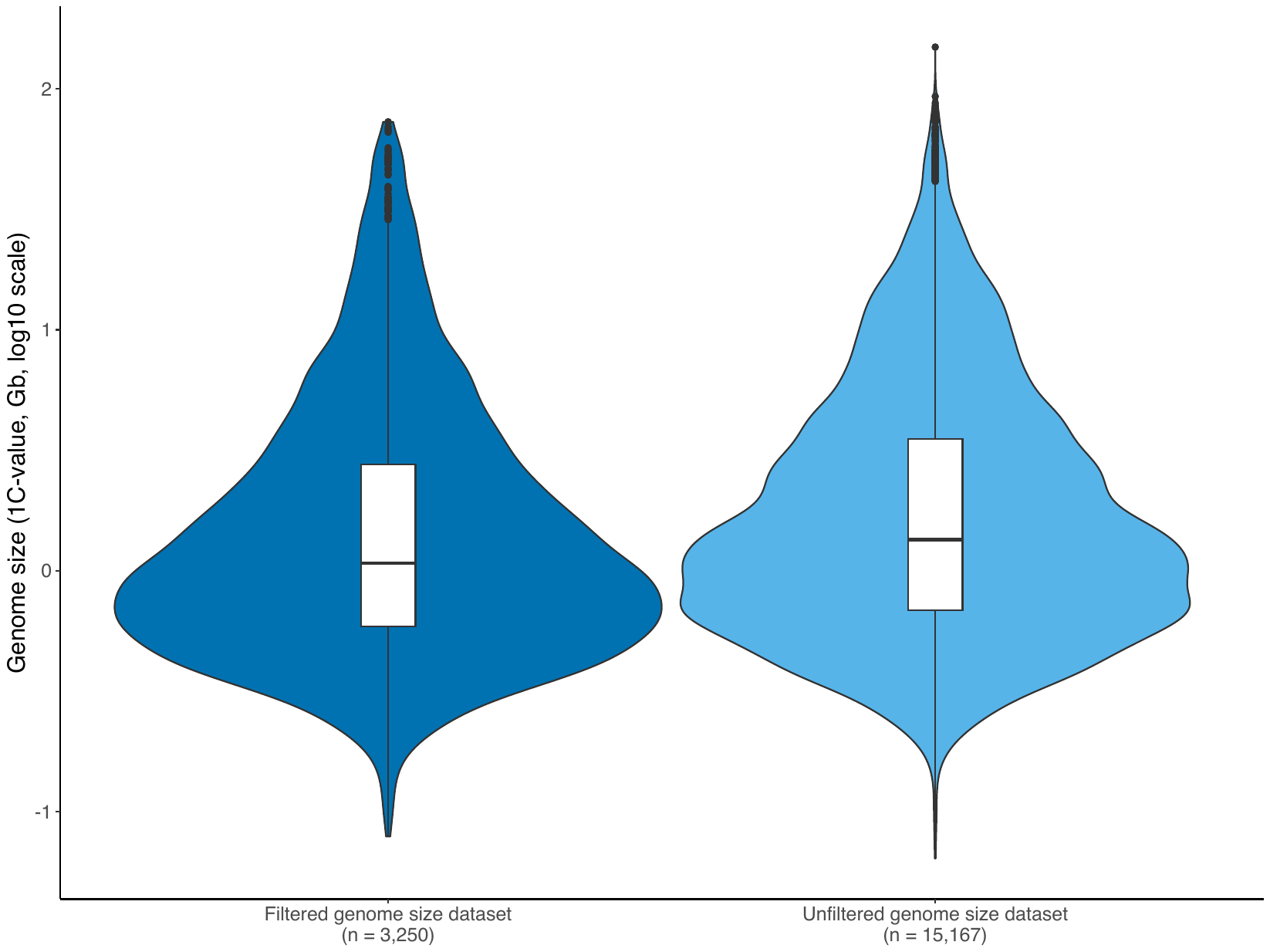

**Figure S6.** Distribution of genome sizes in the 3,250-species dataset assembled here after filtering out species lacking Red List assessments, compared to the unfiltered dataset comprising 15,167 species (Table **S1**).

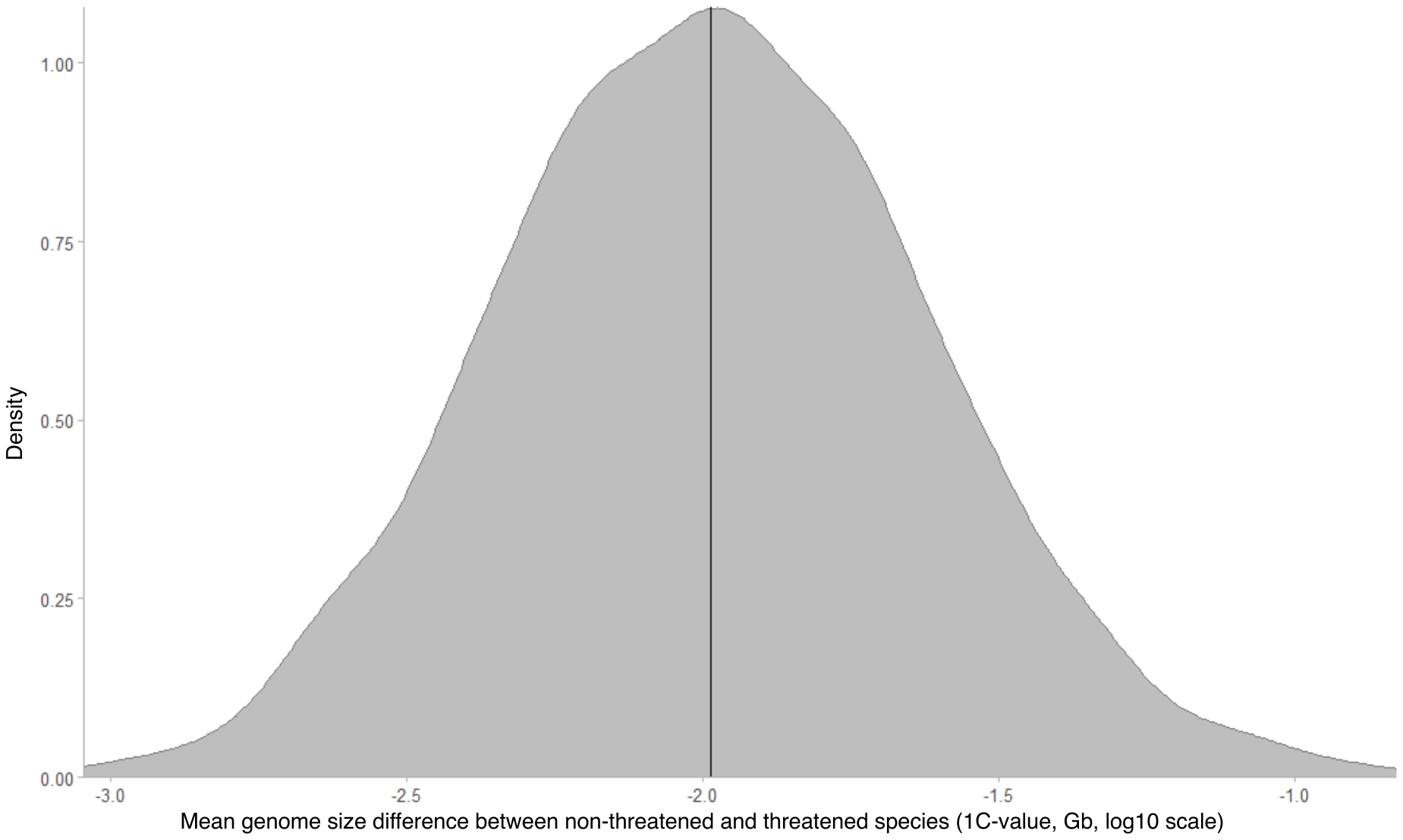

**Figure S7.** Distribution of mean genome size differences between threatened and non-threatened species in the 3,250-species dataset assembled here (indicated by a horizontal black line), compared to mean differences in 999 subsets that were randomly down-sampled without replacement to equalize the proportion of the two threat groupings to 500 species each. The non-threatened grouping comprises Least Concern and Near Threatened species, and the threatened grouping comprises Vulnerable, Endangered, Critically Endangered and Extinct in the Wild species, following the Red List definition of threatened.

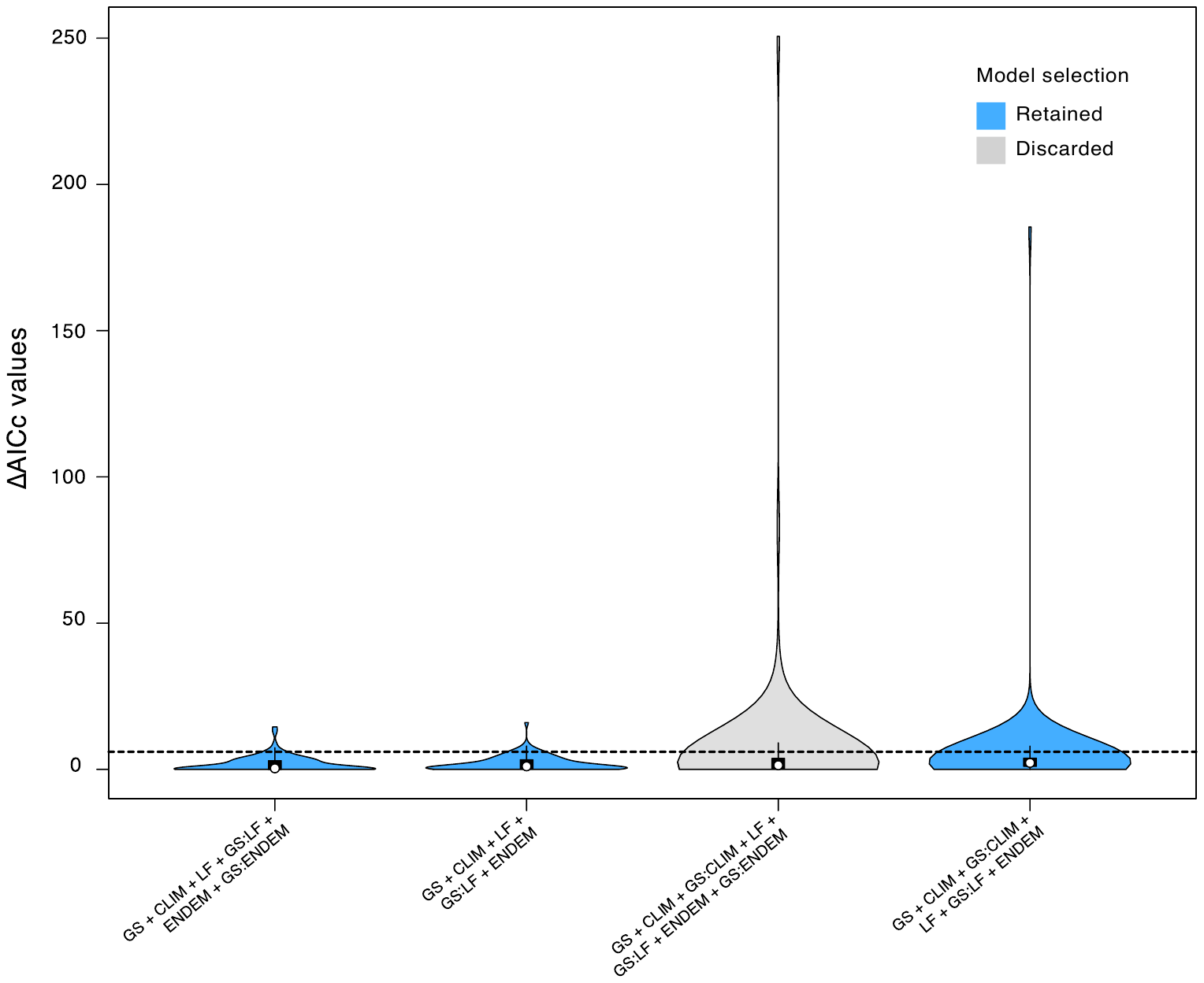

**Figure S8.** Model selection based on ∆AICc values estimated from phylogenetic logistic regressions using a higher threat threshold than the Red List for dichotomizing the response variable into a non-threatened group (comprising Least Concern, Near Threatened, Vulnerable species) and a threatened group (comprising Endangered, Critically Endangered, Extinct in the Wild species); model selection for the point of reference threshold that follows the Red List is shown in Fig. **2**. ∆AICc values were calculated by first ranking all 27 tested models within each of the 100 different phylogenies used as input in analyses, and then summarizing by model across all trees (as shown in Fig. **S2**). The plot shows the distribution of ∆AICc values for the four (of 27) models that had the best AICc value (i.e., ∆AICc = 0) in at least one of the phylogenies (see Table **S5** for support values of the remaining models). Models are ordered by increasing median ∆AICc values. The dashed line at ∆AICc = 6 indicates that models with a median ∆AICc below this cut-off were retained. Abbreviations: GS = genome size, CLIM = climate zone, LF = life form, ENDEM = endemism (proxy for range size).

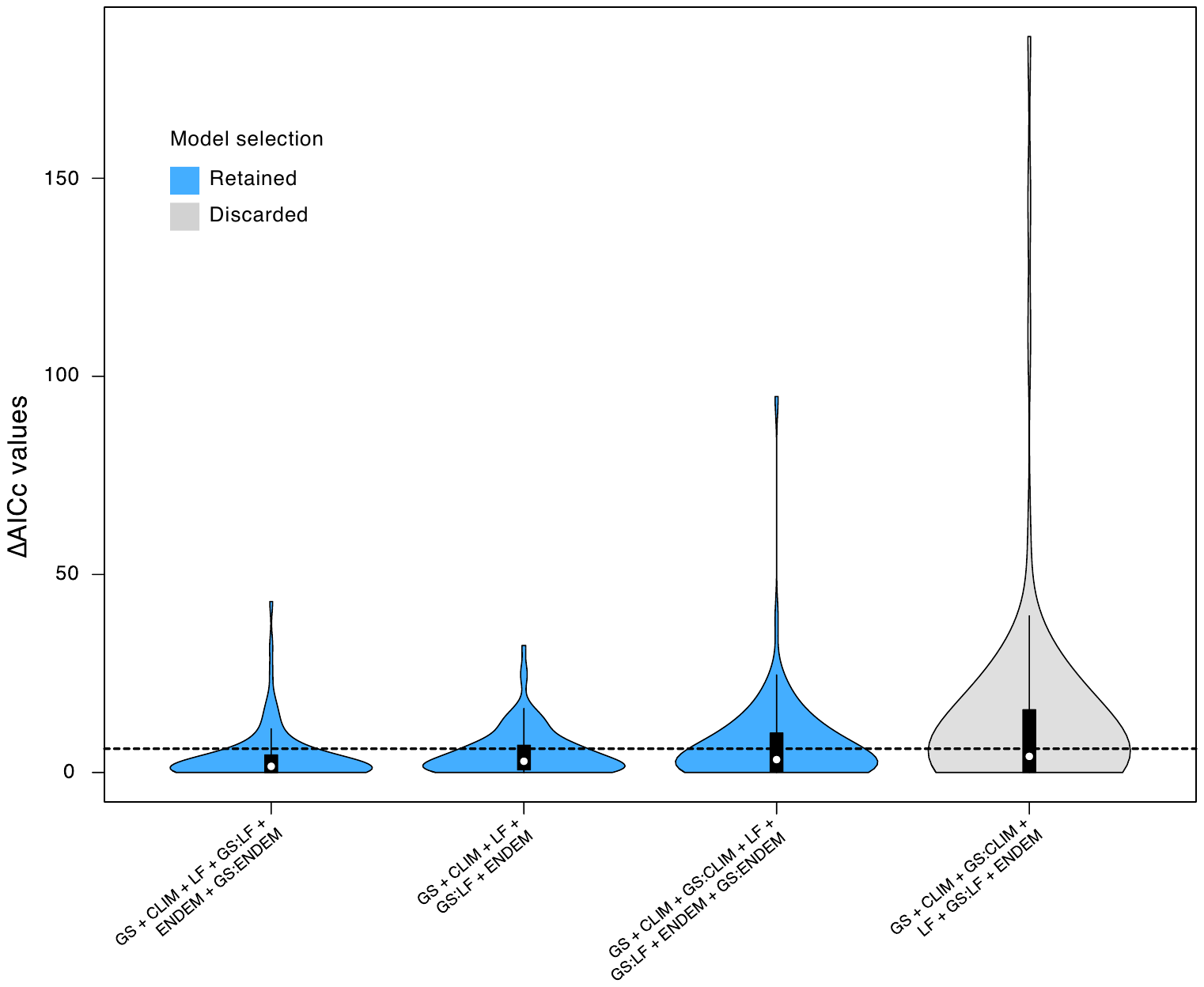

**Figure S9.** Model selection based on ∆AICc values estimated from phylogenetic logistic regressions using a polarizing threat threshold (relative to the Red List) for dichotomizing the response variable into a non-threatened group (comprising Least Concern species) and a threatened group (comprising Vulnerable, Endangered, Critically Endangered, Extinct in the Wild species) while excluding Near Threatened species; model selection for the point of reference threshold that follows the Red List is shown in Fig. **2**. ∆AICc values were calculated by first ranking all 27 tested models within each of the 100 different phylogenies used as input in analyses, and then summarizing by model across all trees (as shown in Fig. **S2**). The plot shows the distribution of ∆AICc values for the four (of 27) models that had the best AICc value (i.e., ∆AICc = 0) in at least one of the phylogenies (see Table **S5** for support values of the remaining models). Models are ordered by increasing median ∆AICc values. The dashed line at ∆AICc = 6 indicates that models with a median ∆AICc below this cut-off were retained. Abbreviations: GS = genome size, CLIM = climate zone, LF = life form, ENDEM = endemism (proxy for range size).

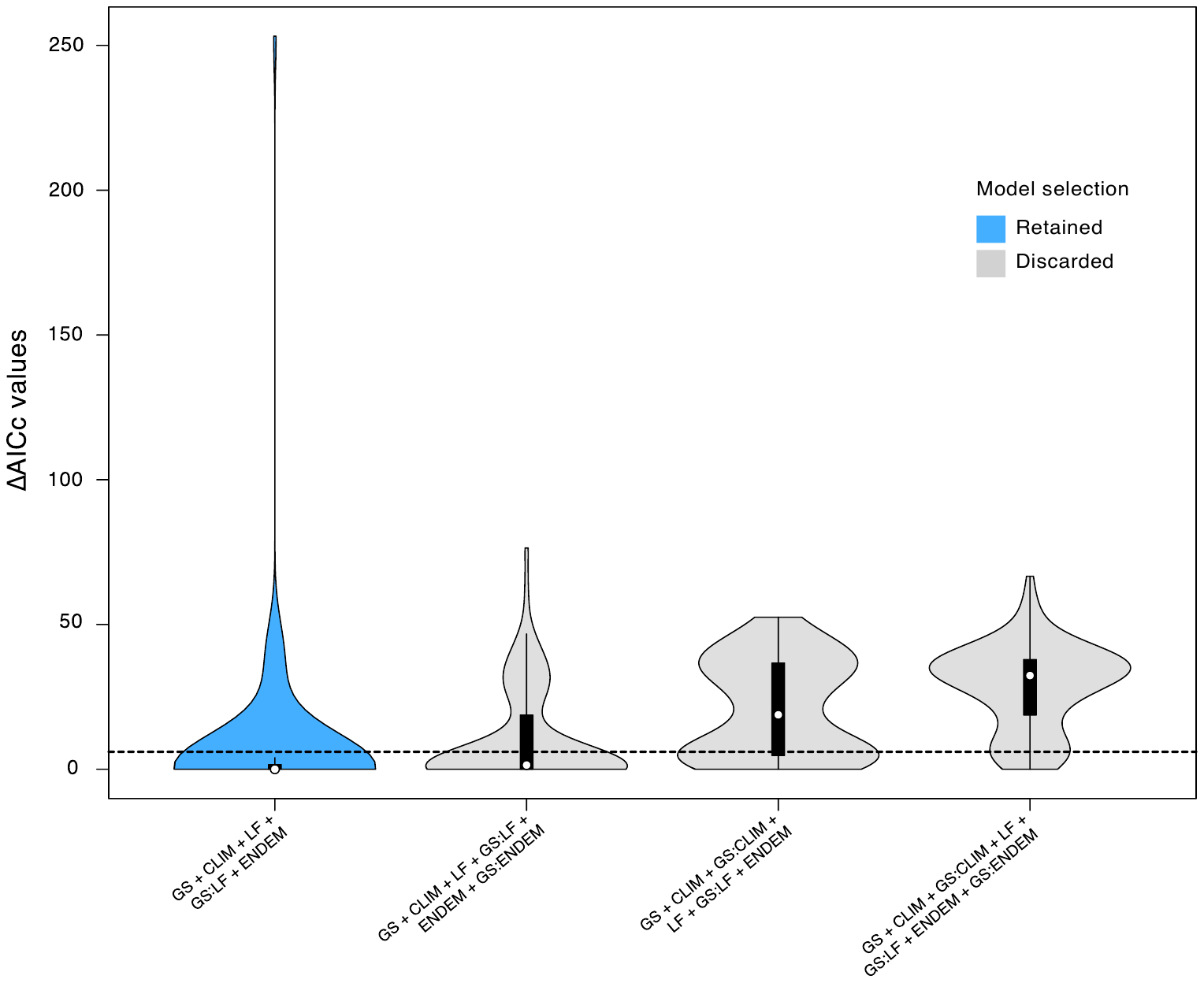

**Figure S10.** Model selection based on ∆AICc values estimated from phylogenetic logistic regressions using a lower threat threshold than the Red List for dichotomizing the response variable into a non-threatened group (comprising Least Concern species) and a threatened group (comprising Near Threatened, Vulnerable, Endangered, Critically Endangered, Extinct in the Wild species); model selection for the point of reference threshold that follows the Red List is shown in Fig. **2**. ∆AICc values were calculated by first ranking all 27 tested models within each of the 100 different phylogenies used as input in analyses, and then summarizing by model across all trees (as shown in Fig. **S2**). The plot shows the distribution of ∆AICc values for the four (of 27) models that had the best AICc value (i.e., ∆AICc = 0) in at least one of the phylogenies (see Table **S5** for support values of the remaining models). Models are ordered by increasing median ∆AICc values. The dashed line at ∆AICc = 6 indicates that models with a median ∆AICc below this cut-off were retained. Abbreviations: GS = genome size, CLIM = climate zone, LF = life form, ENDEM = endemism (proxy for range size).

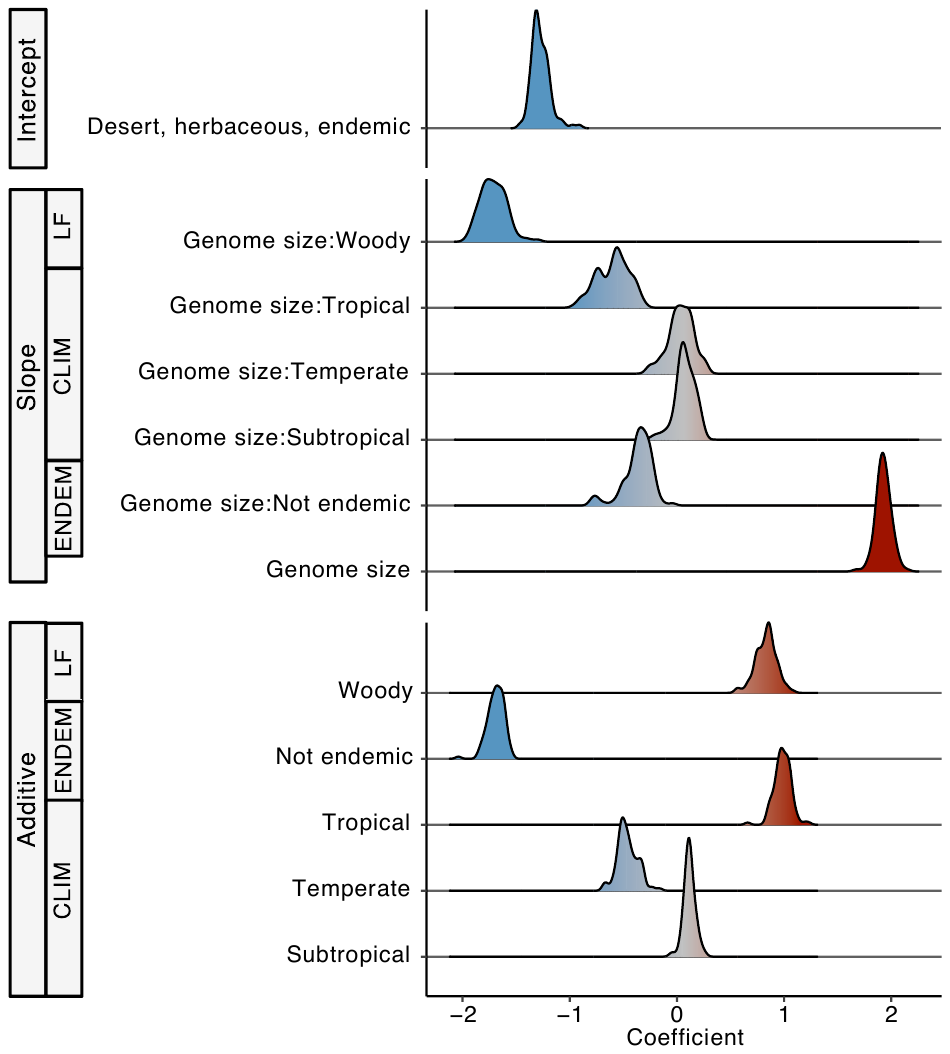

**Figure S11.** Distribution of coefficients for one of two best models (i.e., extinction risk ~ GS + CLIM + GS:CLIM + LF + GS:LF + ENDEM + GS:ENDEM) identified in phylogenetic logistic regressions using 100 different phylogenies as input and the Red List definition of threat as a point of reference for dichotomizing the response variable into a non-threatened grouping (comprising Least Concern, Near Threatened species) and a threatened grouping (comprising Vulnerable, Endangered, Critically Endangered, Extinct in the Wild species). Plots show the coefficient frequency distributions of all the model terms across all 100 of the conducted analyses. These coefficients are summarized in Table **S6** along with their 95% confidence intervals from bootstrap analysis; they were also used to estimate the blue curves in Fig. **3** for predicting the probability of threat in angiosperms based on this model. Abbreviations: GS = genome size, CLIM = climate zone, LF = life form, ENDEM = endemism (proxy for range size).

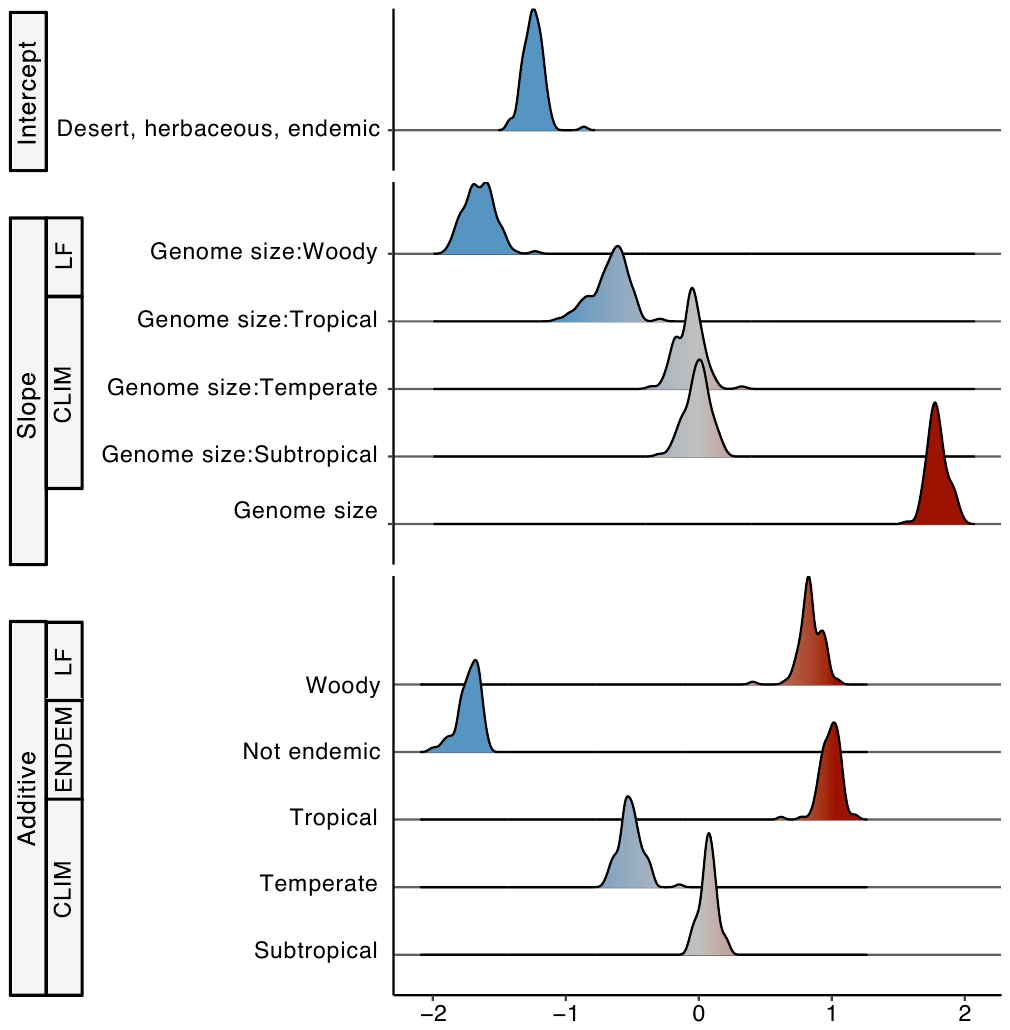

Coefficient

**Figure S12.** Distribution of coefficients for a second of two best models (i.e., extinction risk ~ GS + CLIM + GS:CLIM + LF + GS:LF + ENDEM) identified in phylogenetic logistic regressions using 100 different phylogenies as input and the Red List definition of threat as a point of reference for dichotomizing the response variable into a non-threatened grouping (comprising Least Concern, Near Threatened species) and a threatened grouping (comprising Vulnerable, Endangered, Critically Endangered, Extinct in the Wild species). Plots show the coefficient frequency distributions of all the model terms across all 100 of the conducted analyses. These coefficients are summarized in Table **S6** along with their 95% confidence intervals from bootstrap analysis; they were also used to estimate the orange curves in Fig. **3** for predicting the probability of threat in angiosperms based on this model. Abbreviations: GS = genome size, CLIM = climate zone, LF = life form, ENDEM = endemism (proxy for range size).

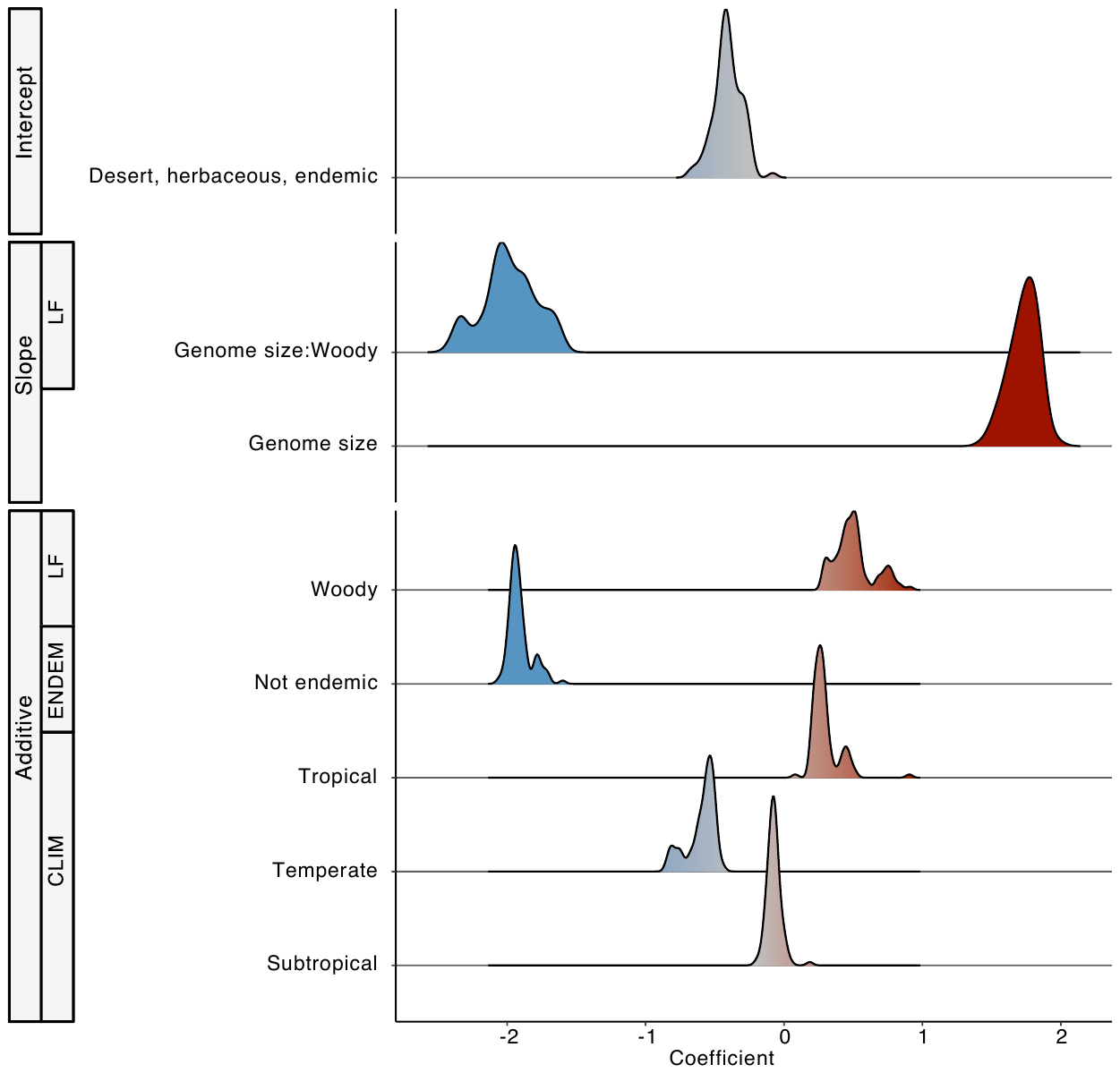

**Figure S13.** Distribution of coefficients for the single best model (i.e., extinction risk ~ GS + CLIM + LF + GS:LF + ENDEM) identified in phylogenetic logistic regressions using 100 different phylogenies as input and a lower threat threshold than the Red List for dichotomizing the response variable into a non-threatened grouping (comprising Least Concern species) and a threatened grouping (comprising Near Threatened, Vulnerable, Endangered, Critically Endangered, Extinct in the Wild species). Plots show the coefficient frequency distributions of all the model terms across all 100 of the conducted analyses. These coefficients are summarized in Table **S6** along with their 95% confidence intervals from bootstrap analysis; they were also used to estimate the dashed curves in Fig. **S20** for predicting the probability of threat in angiosperms based on this model. Abbreviations: GS = genome size, CLIM = climate zone, LF = life form, ENDEM = endemism (proxy for range size).

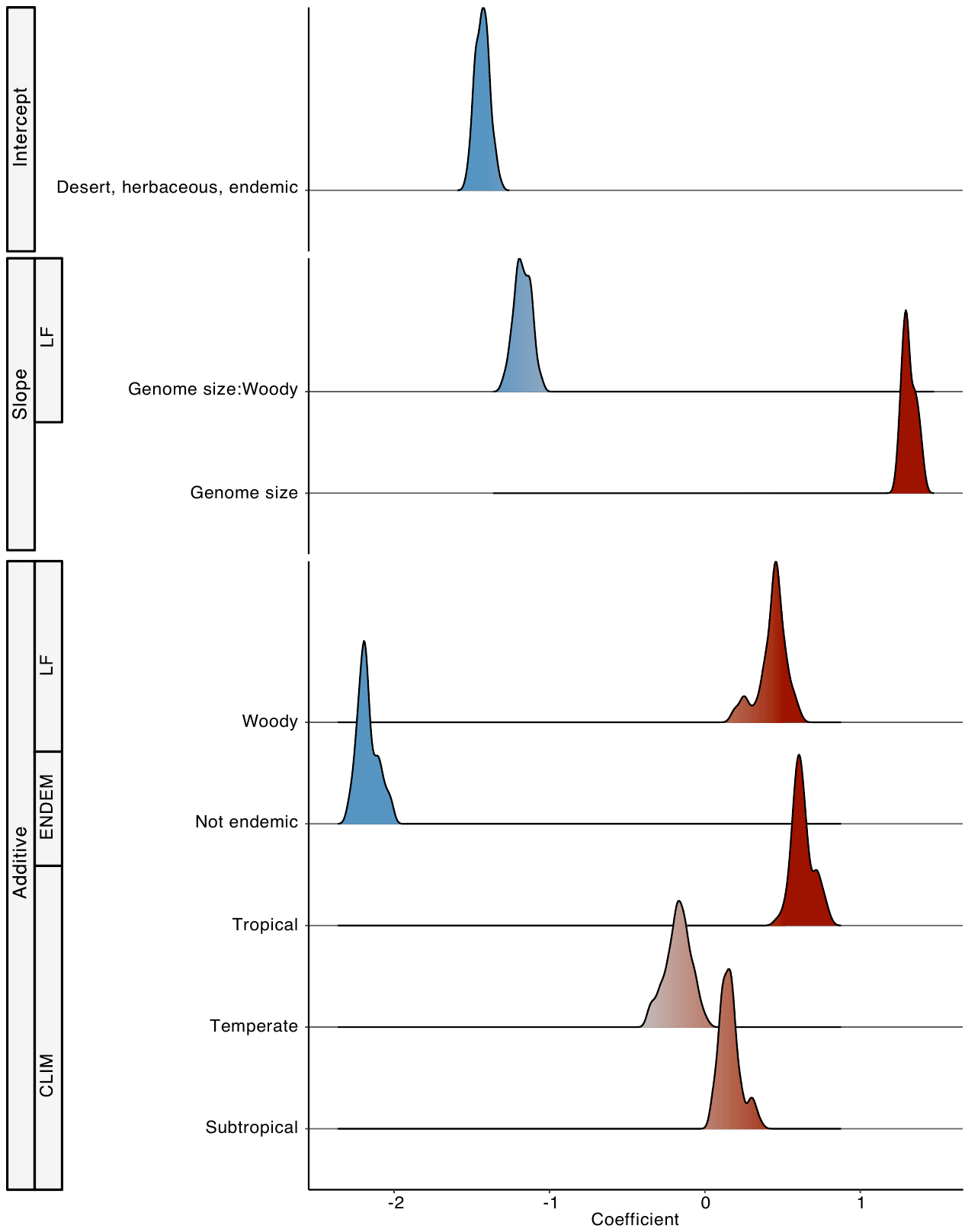

**Figure S14.** Distribution of coefficients for one of three best models (i.e., extinction risk ~ GS + CLIM + LF + GS:LF + ENDEM) identified in phylogenetic logistic regressions using 100 different phylogenies as input and a higher threat threshold than the Red List for dichotomizing the response variable into a non-threatened grouping (comprising Least Concern, Near Threatened, Vulnerable species) and a threatened grouping (comprising Endangered, Critically Endangered, Extinct in the Wild species). Plots show the coefficient frequency distributions of all the model terms across all 100 of the conducted analyses. These coefficients are summarized in Table **S6** along with their 95% confidence intervals from bootstrap analysis; they were also used to estimate the double-dashed curves in Fig. **S20** for predicting the probability of threat in angiosperms based on this model. Abbreviations: GS = genome size, CLIM = climate zone, LF = life form, ENDEM = endemism (proxy for range size).

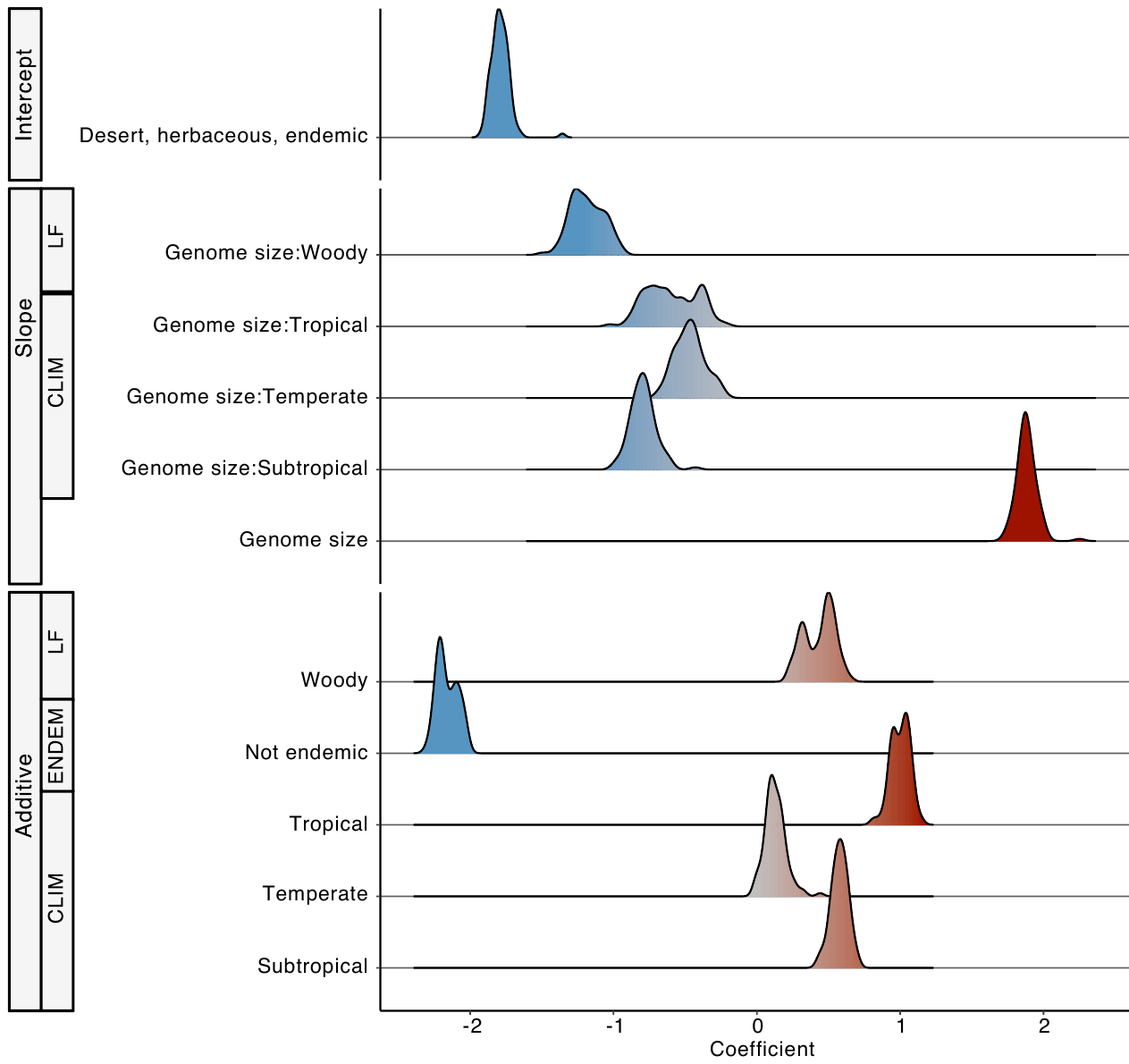

**Figure S15.** Distribution of coefficients for a second of three best models (i.e., extinction risk ~ GS + CLIM + GS:CLIM + LF + GS:LF + ENDEM) identified in phylogenetic logistic regressions using 100 different phylogenies as input and a higher threat threshold than the Red List for dichotomizing the response variable into a non-threatened grouping (comprising Least Concern, Near Threatened, Vulnerable species) and a threatened grouping (comprising Endangered, Critically Endangered, Extinct in the Wild species). Plots show the coefficient frequency distributions of all the model terms across all 100 of the conducted analyses. These coefficients are summarized in Table **S6** along with their 95% confidence intervals from bootstrap analysis; they were also used to estimate the double-dashed curves in Fig. **S21** for predicting the probability of threat in angiosperms based on this model. Abbreviations: GS = genome size, CLIM = climate zone, LF = life form, ENDEM = endemism (proxy for range size).

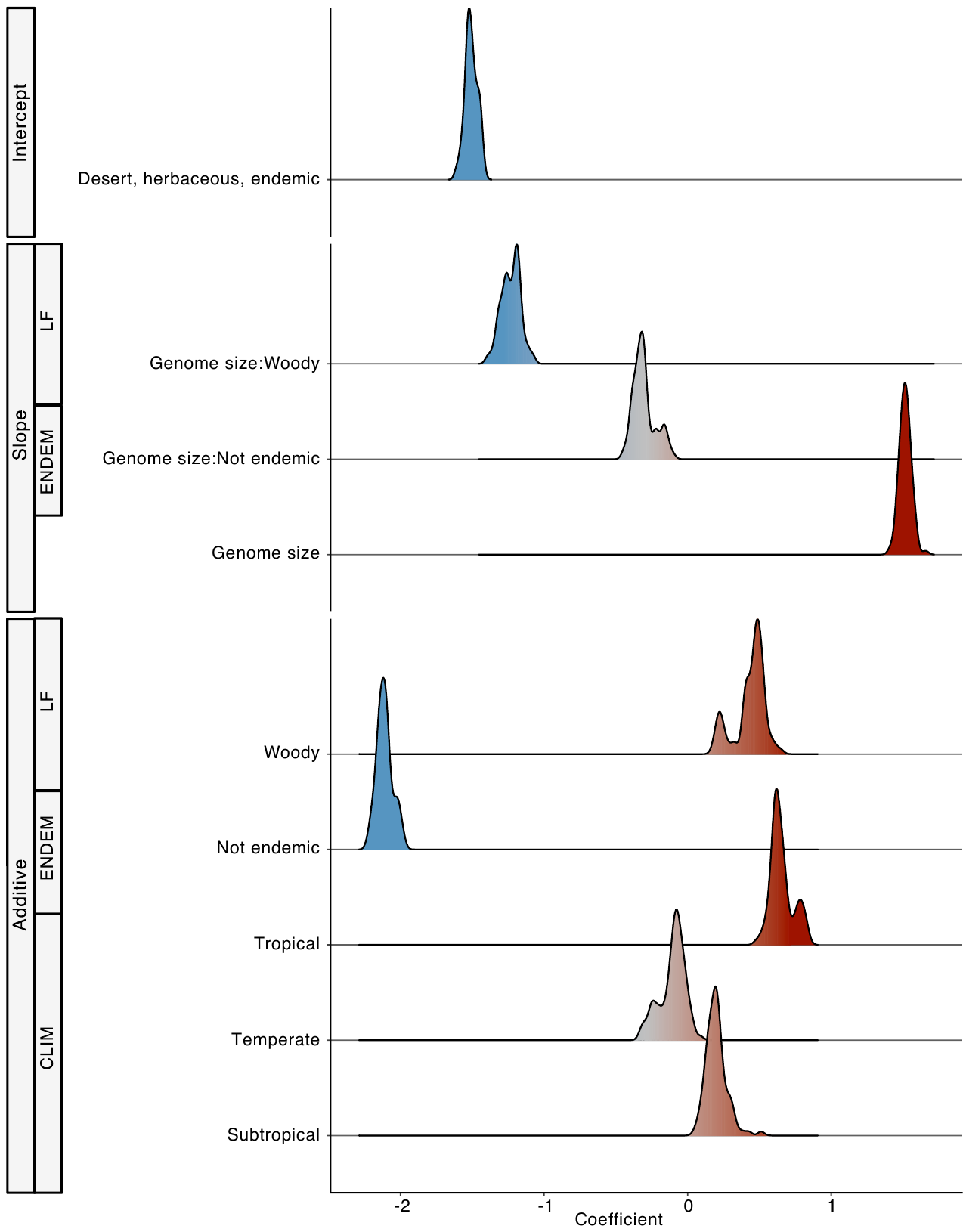

**Figure S16.** Distribution of coefficients for a third of three best models (i.e., extinction risk ~ GS + CLIM + LF + GS:LF + ENDEM + GS:ENDEM) identified in phylogenetic logistic regressions using 100 different phylogenies as input and a higher threat threshold than the Red List for dichotomizing the response variable into a non-threatened grouping (comprising Least Concern, Near Threatened, Vulnerable species) and a threatened grouping (comprising Endangered, Critically Endangered, Extinct in the Wild species). Plots show the coefficient frequency distributions of all the model terms across all 100 of the conducted analyses. These coefficients are summarized in Table **S6** along with their 95% confidence intervals from bootstrap analysis; they were also used to estimate the double-dashed curves in Fig. **S22** for predicting the probability of threat in angiosperms based on this model. Abbreviations: GS = genome size, CLIM = climate zone, LF = life form, ENDEM = endemism (proxy for range size).

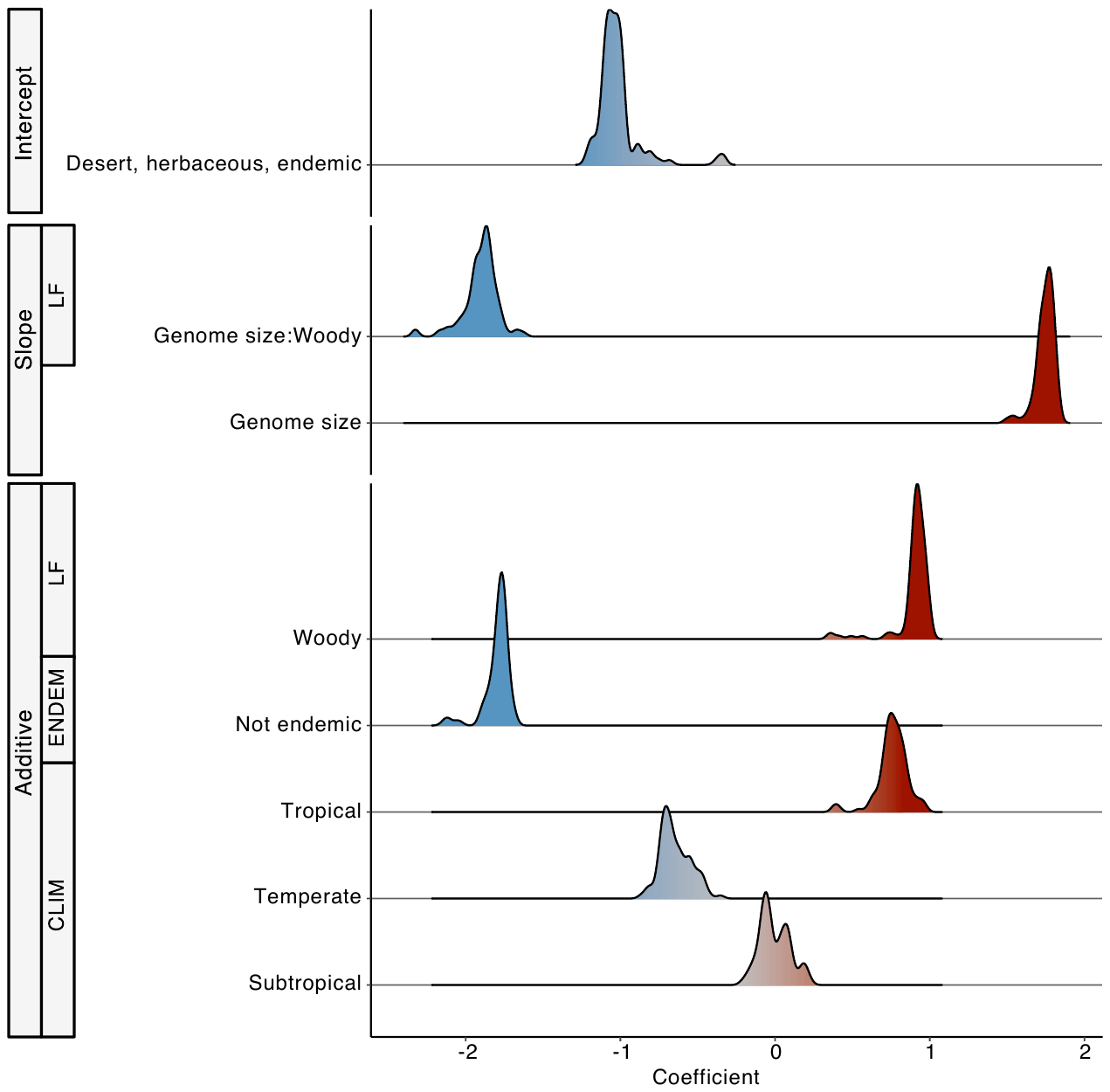

**Figure S17.** Distribution of coefficients for one of three best models (i.e., extinction risk ~ GS + CLIM + LF + GS:LF + ENDEM) identified in phylogenetic logistic regressions using 100 different phylogenies as input and a polarizing threshold (compared to the Red List) for dichotomizing the response variable into a non-threatened grouping (comprising Least Concern species) and a threatened grouping (comprising Vulnerable, Endangered, Critically Endangered, Extinct in the Wild species), while excluding Near Threatened species. Plots show the coefficient frequency distributions of all the model terms across all 100 of the conducted analyses. These coefficients are summarized in Table **S6** along with their 95% confidence intervals from bootstrap analysis; they were also used to estimate the solid curves in Fig. **S20** for predicting the probability of threat in angiosperms based on this model. Abbreviations: GS = genome size, CLIM = climate zone, LF = life form, ENDEM = endemism (proxy for range size).

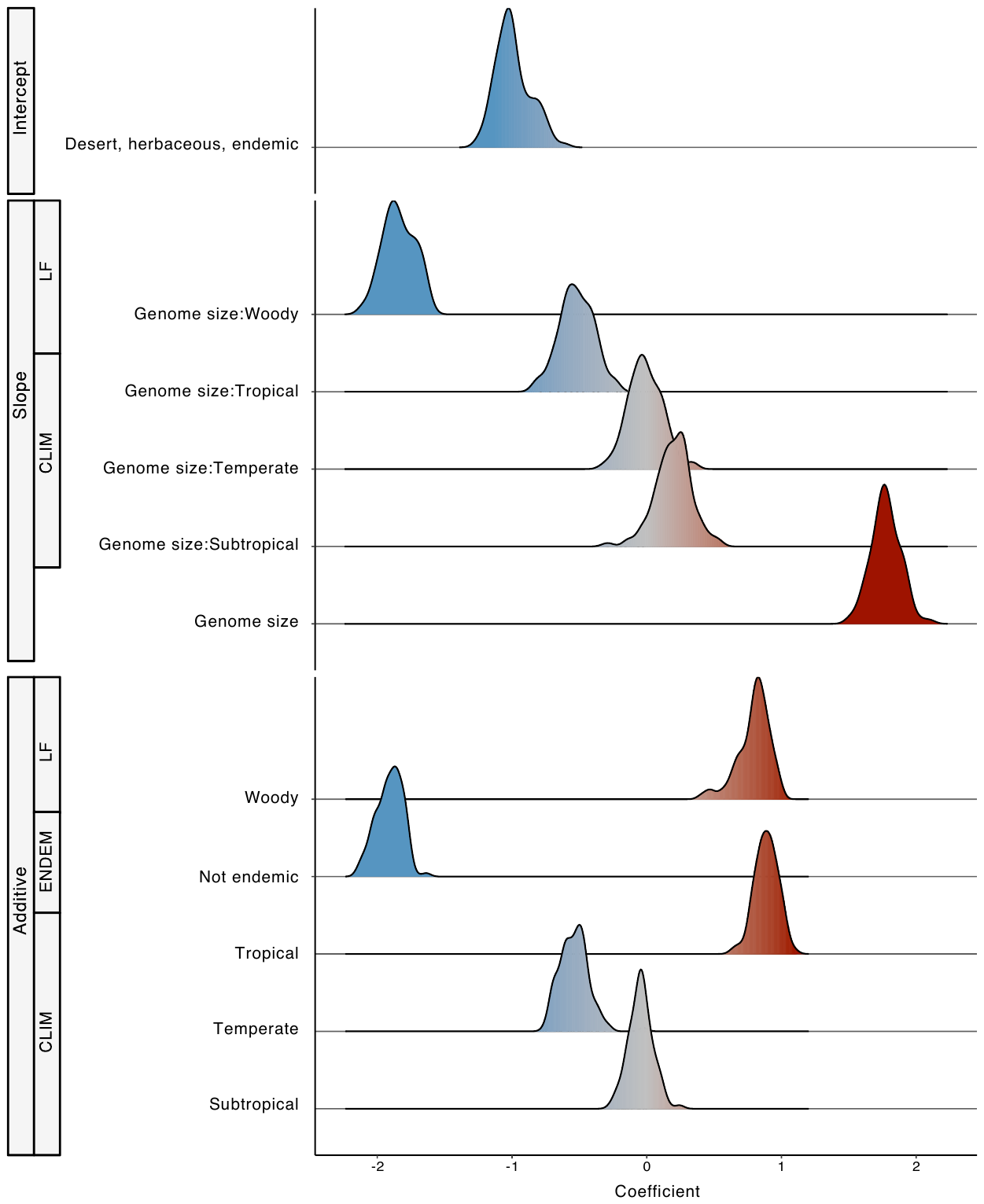

**Figure S18.** Distribution of coefficients for a second of three best models (i.e., extinction risk ~ GS + CLIM + GS:CLIM + LF + GS:LF + ENDEM) identified in phylogenetic logistic regressions using 100 different phylogenies as input and a polarizing threshold (compared to the Red List) for dichotomizing the response variable into a non-threatened grouping (comprising Least Concern species) and a threatened grouping (comprising Vulnerable, Endangered, Critically Endangered, Extinct in the Wild species), while excluding Near Threatened species. Plots show the coefficient frequency distributions of all the model terms across all 100 of the conducted analyses. These coefficients are summarized in Table **S6** along with their 95% confidence intervals from bootstrap analysis; they were also used to estimate the solid curves in Fig. **S21** for predicting the probability of threat in angiosperms based on this model. Abbreviations: GS = genome size, CLIM = climate zone, LF = life form, ENDEM = endemism (proxy for range size).

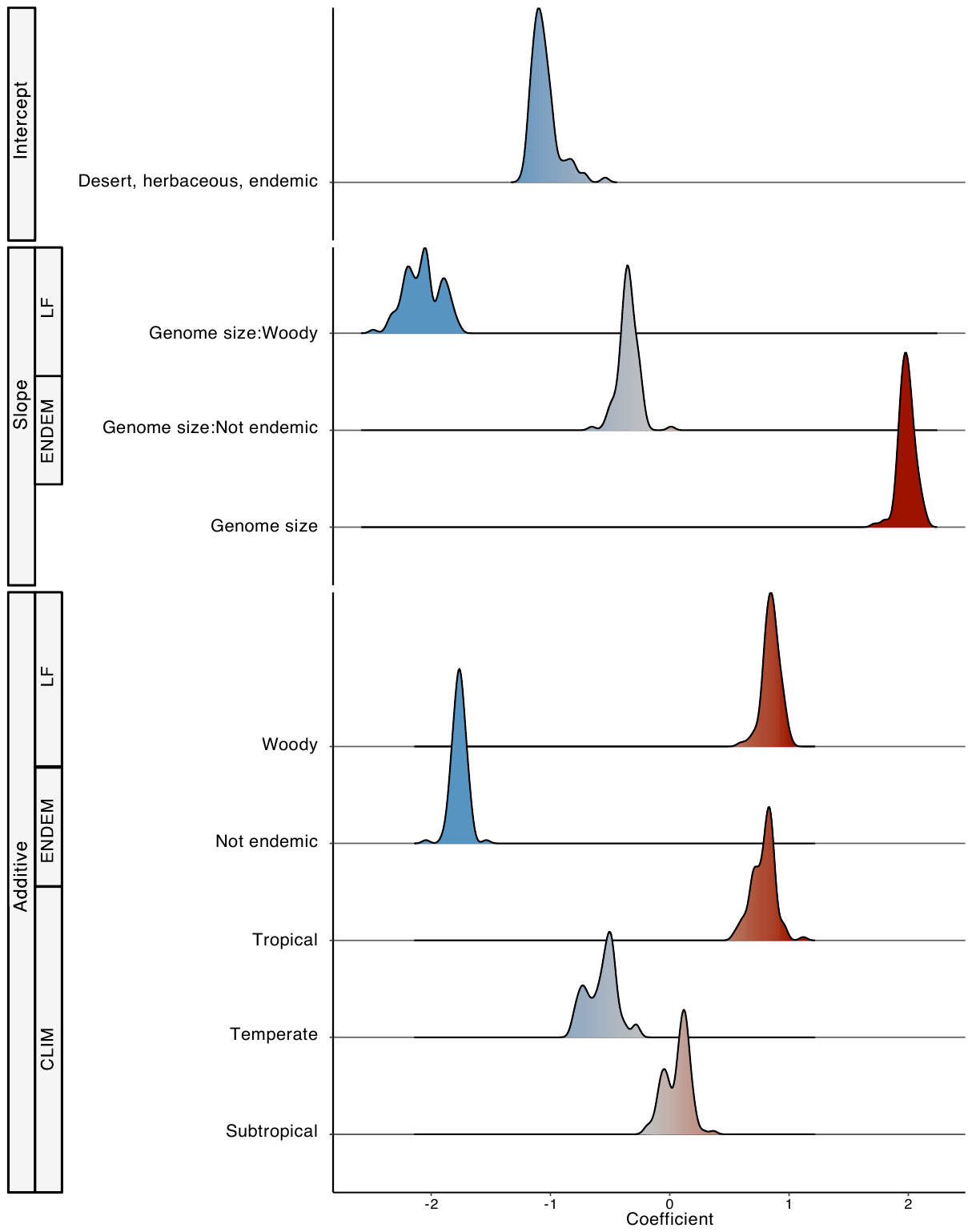

**Figure S19.** Distribution of coefficients for a third of three best models (i.e., extinction risk ~ GS + CLIM + LF + GS:LF + ENDEM + GS:ENDEM) identified in phylogenetic logistic regressions using 100 different phylogenies as input and a polarizing threshold (compared to the Red List) for dichotomizing the response variable into a non-threatened grouping (comprising Least Concern species) and a threatened grouping (comprising Vulnerable, Endangered, Critically Endangered, Extinct in the Wild species), while excluding Near Threatened species. Plots show the coefficient frequency distributions of all the model terms across all 100 of the conducted analyses. These coefficients are summarized in Table **S6** along with their 95% confidence intervals from bootstrap analysis; they were also used to estimate the solid curves in Fig. **S22** for predicting the probability of threat in angiosperms based on this model. Abbreviations: GS = genome size, CLIM = climate zone, LF = life form, ENDEM = endemism (proxy for range size).

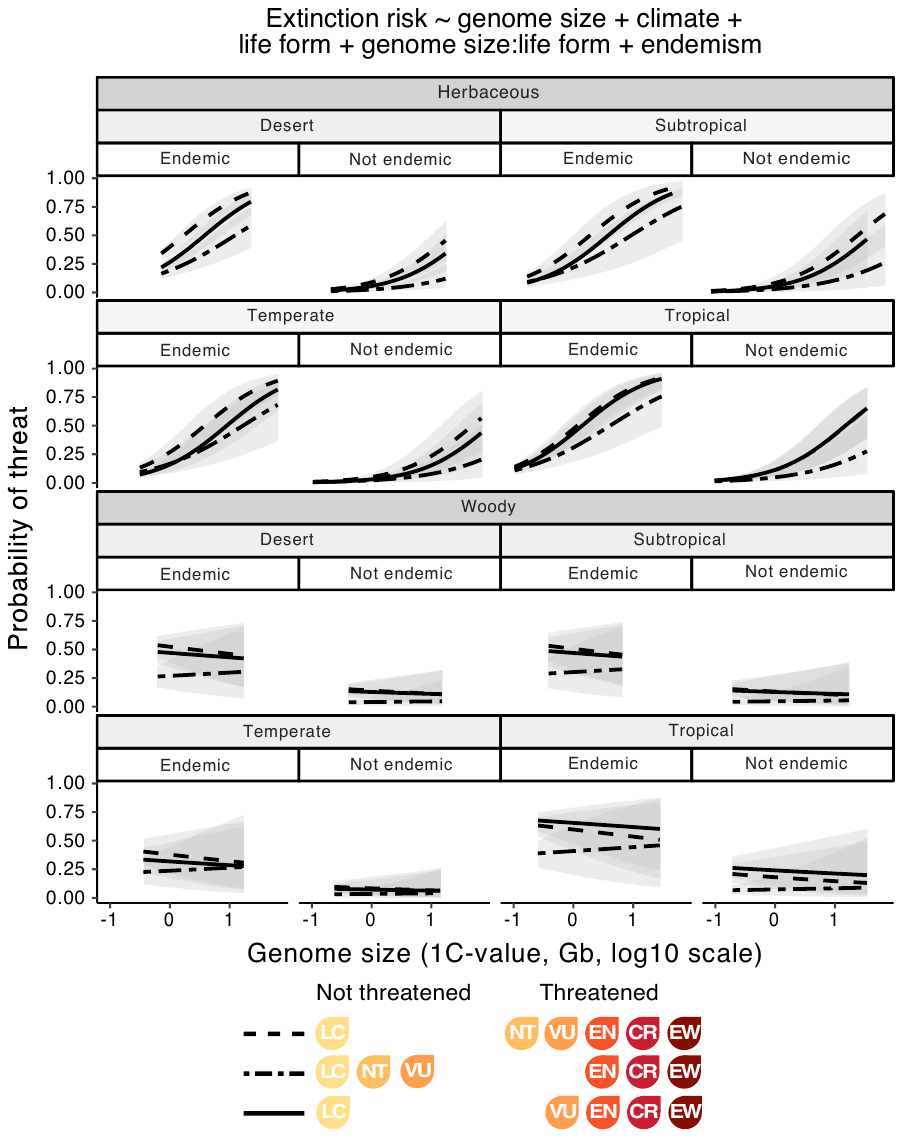

**Figure S20**. Phylogenetic logistic regression curves predicting the probability of threat in angiosperms as a function of genome size, life form, climate zone and endemism, showing a best model selected in common across analyses where extinction risk was dichotomized using comparison threat thresholds that were lower (dashed line), higher (double-dashed line), or polarizing (solid line) relative to the Red List definition of threat (used as a point of reference across analyses here). Curves represent mean coefficients summarized across the 100 different phylogenies used as input for each variant threat threshold and the shaded areas indicate 95% confidence intervals from bootstrap analysis. The start and end points of the curves and confidence intervals indicate the minimum and maximum genome sizes represented across the different variable partitions (also given in Table **S3**).

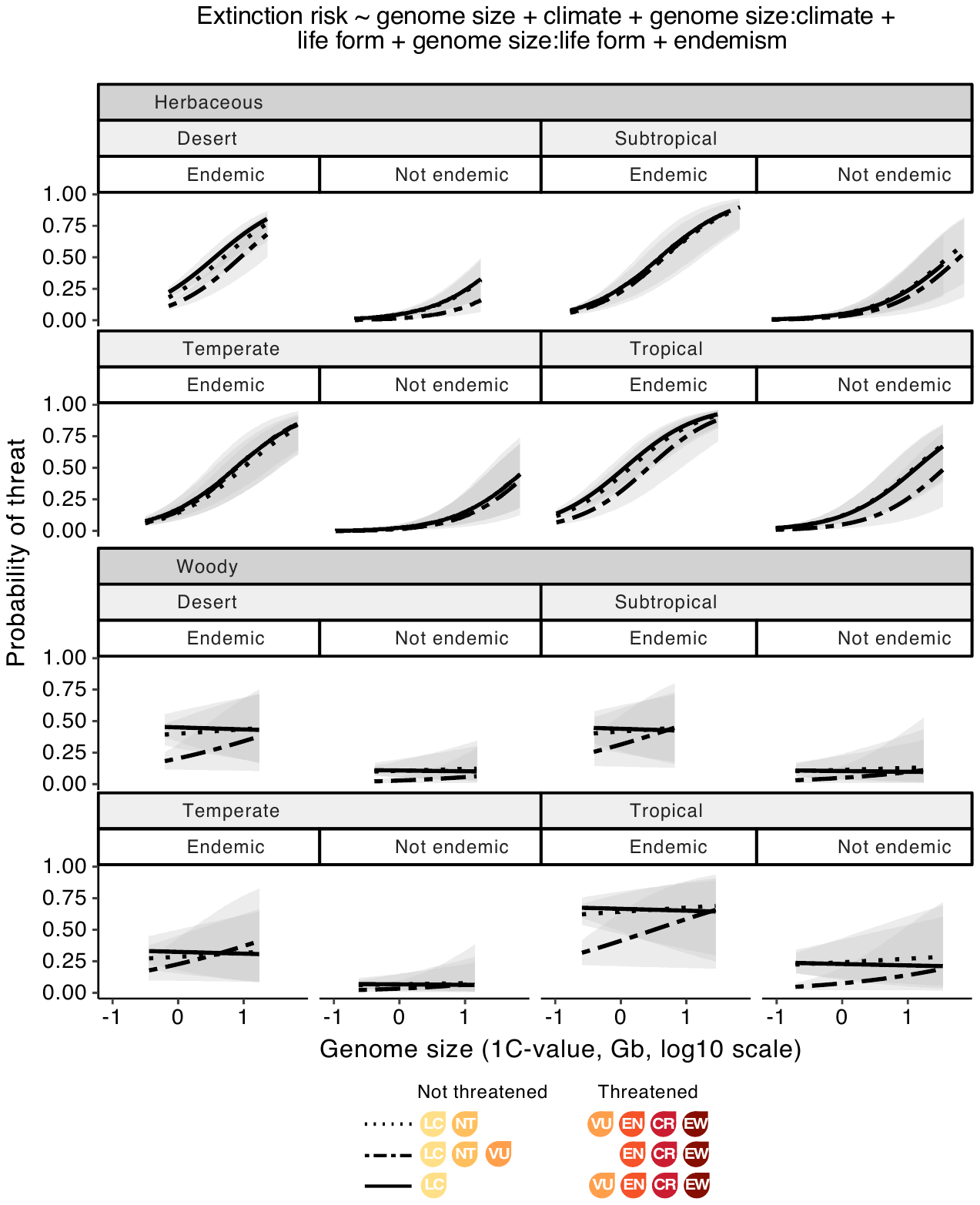

**Figure S21**. Phylogenetic logistic regression curves predicting the probability of threat in angiosperms as a function of genome size, life form, climate zone and endemism, showing a best model selected in common across analyses where extinction risk was dichotomized using the Red List definition of threat as a point of reference (dotted line) or using comparison threat thresholds that were higher (double-dashed line), or polarizing (solid line). Curves represent mean coefficients summarized across the 100 different phylogenies used as input for each variant threat threshold and the shaded areas indicate 95% confidence intervals from bootstrap analysis. The start and end points of the curves and confidence intervals indicate the minimum and maximum genome sizes represented across the different variable partitions (also given in Table **S3**).

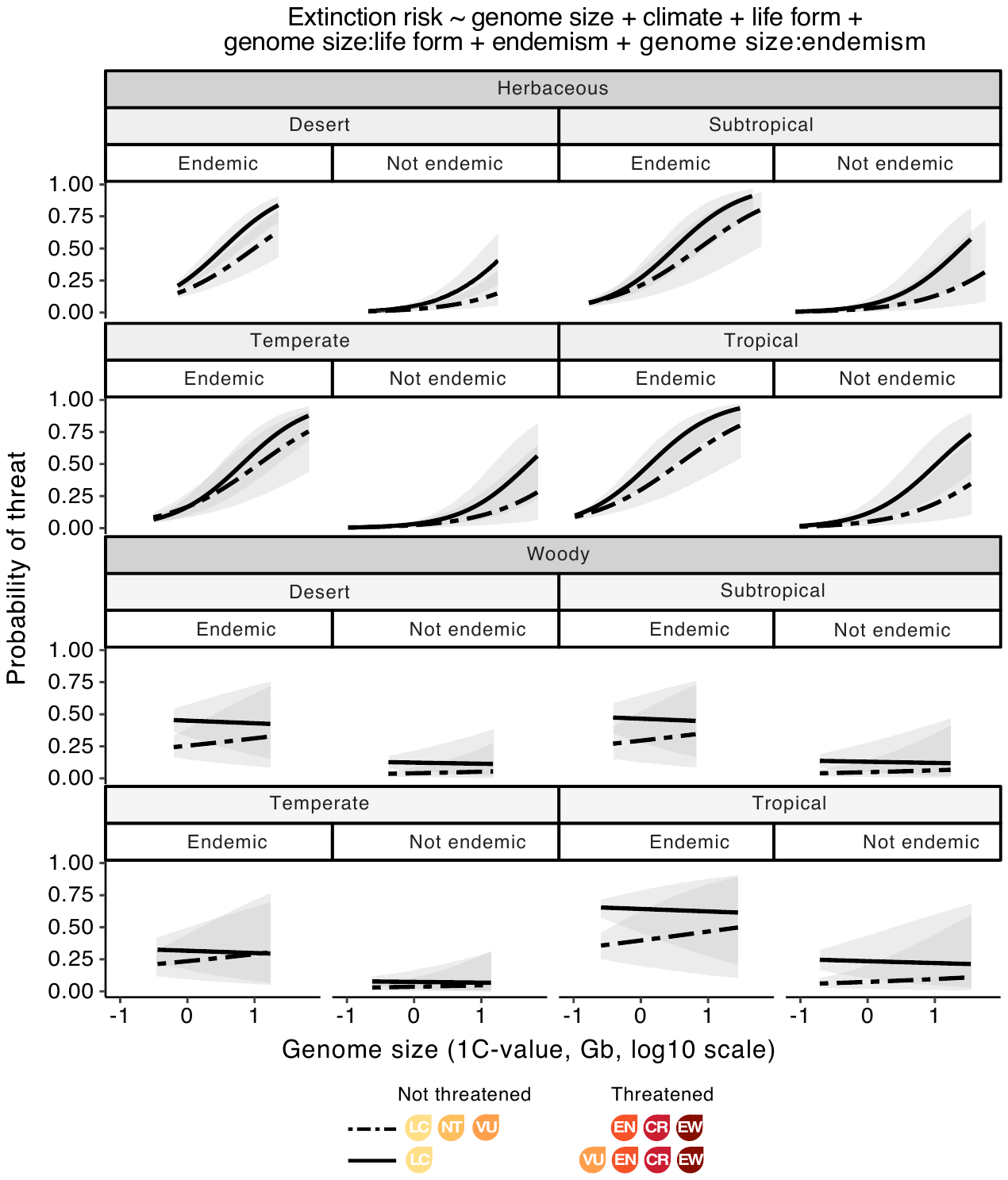

**Figure S22**. Phylogenetic logistic regression curves predicting the probability of threat in angiosperms as a function of genome size, life form, climate zone and endemism, showing a best model selected in common across analyses where extinction risk was dichotomized using comparison threat thresholds that were higher (double-dashed line), or polarizing (solid line) relative to the Red List definition of threat (used as a point of reference across analyses here). Curves represent mean coefficients summarized across the 100 different phylogenies used as input for each variant threat threshold and the shaded areas indicate 95% confidence intervals from bootstrap analysis. The start and end points of the curves and confidence intervals indicate the minimum and maximum genome sizes represented across the different variable partitions (also given in Table **S3**).

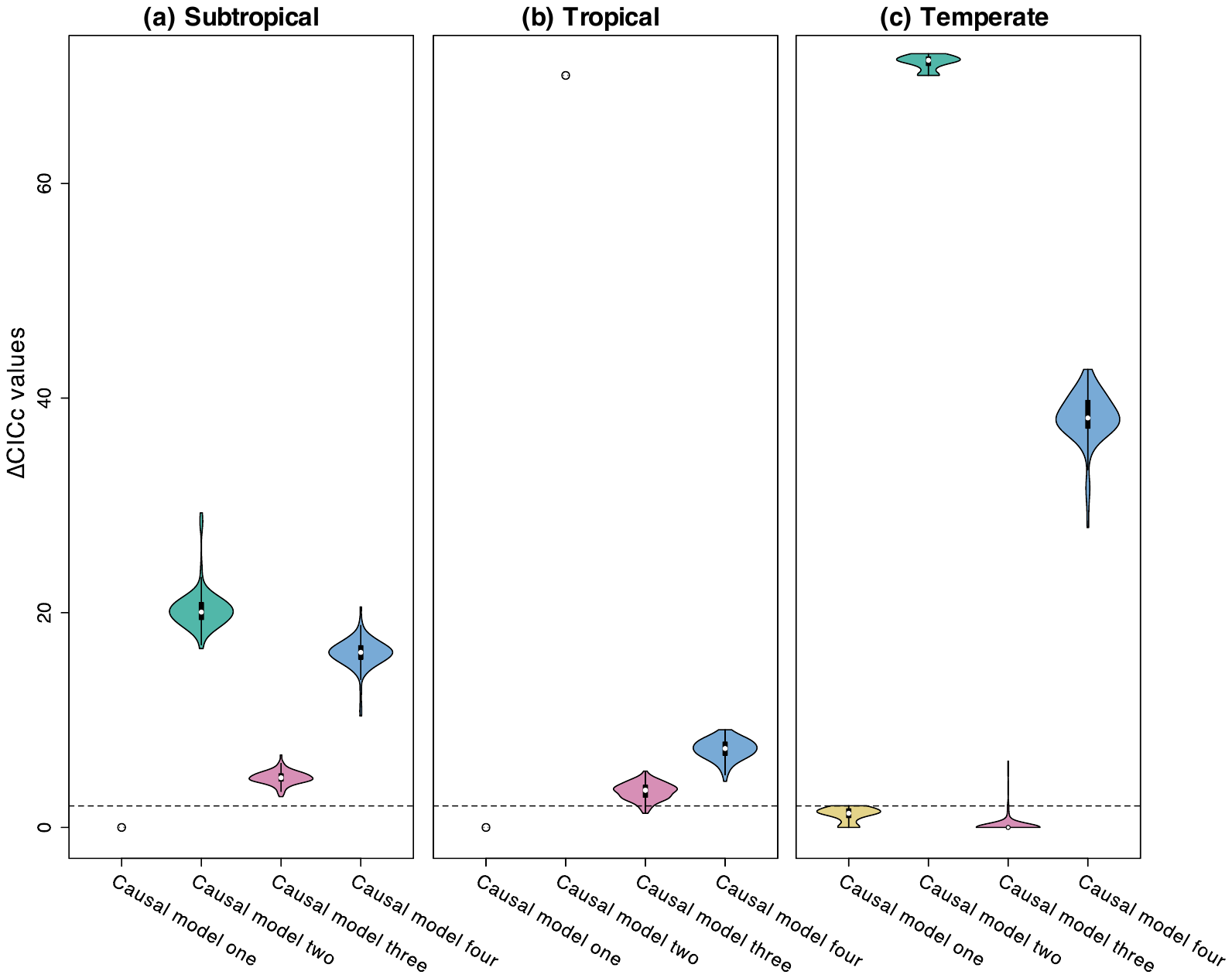

**Figure S23**. Support for four competing causal models (depicted in Fig. **S4**) based on ∆CICc values from confirmatory phylogenetic path analysis for (a) subtropical, (b) tropical and (c) temperate species using the Red List definition of threat as a point of reference for grouping Least Concern and Near Threatened species as non-threatened, and Vulnerable, Endangered, Critically Endangered and Extinct in the Wild species as threatened (desert species are not shown as analyses for this subset did not converge). Values were estimated by ranking hypotheses within each of the 100 variant phylogenies used as input in analyses, and then summarized by hypothesis. The dashed line indicates a ∆CICc of 2, as models with a median ∆CICc below this threshold were retained.

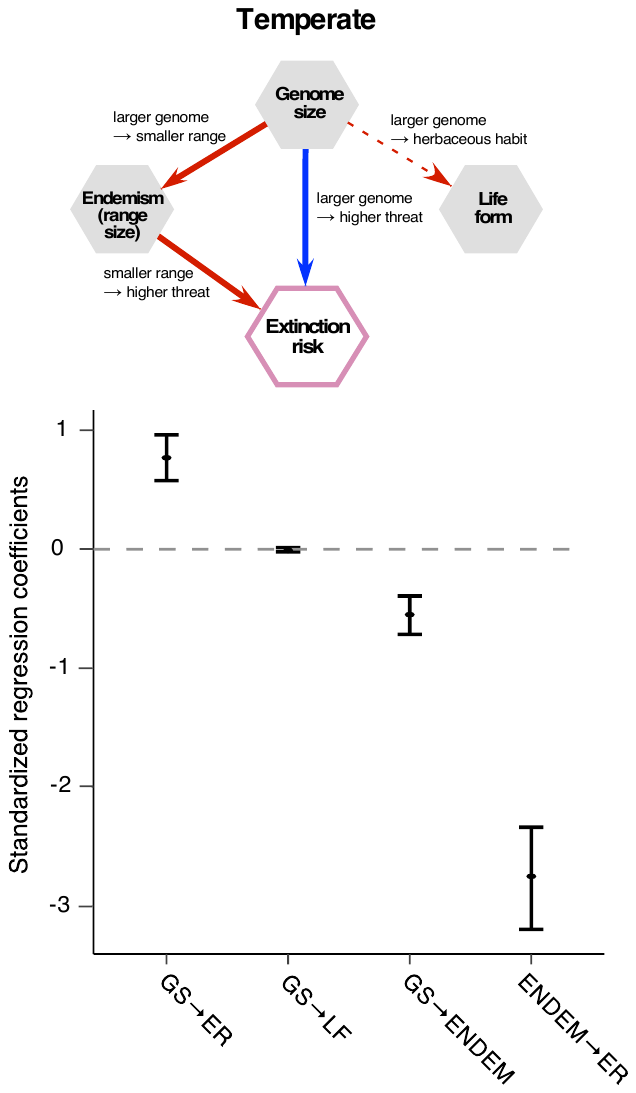

**Figure S24**. Schematic and coefficients for causal model three, one of two best models obtained from confirmatory phylogenetic path analyses for temperate species based on ∆CICc model selection (Fig. **S23**). An additional best causal model for this climate subset (i.e., causal model one) is shown in Fig. **4**. Positive links between variables are indicated by blue arrows in the model schematic and negative links by red arrows; solid and dashed arrows indicate significant and non-significant links, respectively. Plots show the mean coefficients and 95% confidence intervals (the latter obtained from bootstrap analyses) for relationships present in the model above, after summarising across the 100 different phylogenies used as input in individual path analyses for temperate species. Confidence intervals fully above or below the dashed line (coefficient = 0) indicate significant coefficients. Abbreviations: GS = genome size, ER = extinction risk, LF = life form, ENDEM = endemism (proxy for range size).
