## Supplementary Table 2 for "Genome size is positively correlated with extinction risk in herbaceous angiosperms"

**Table S2.** Representation of the 3,250 species in the genome size dataset across Red List Categories and sample size when partitioned by life form, climatic zone, and endemism.

**__________________________________________________________________**

| **Red List Category** | **No. of**  **species** | **Life form** | **Climate** | **Endemism** |
| --- | --- | --- | --- | --- |
| **Least Concern** | 2,388 |  |  |  |
|  | 8 | Herbaceous | Desert | Endemic |
|  | 25 | Herbaceous | Desert | Not endemic |
|  | 30 | Herbaceous | Subtropical | Endemic |
|  | 146 | Herbaceous | Subtropical | Not endemic |
|  | 21 | Herbaceous | Temperate | Endemic |
|  | 659 | Herbaceous | Temperate | Not endemic |
|  | 26 | Herbaceous | Tropical | Endemic |
|  | 271 | Herbaceous | Tropical | Not endemic |
|  | 13 | Woody | Desert | Endemic |
|  | 47 | Woody | Desert | Not endemic |
|  | 50 | Woody | Subtropical | Endemic |
|  | 141 | Woody | Subtropical | Not endemic |
|  | 36 | Woody | Temperate | Endemic |
|  | 434 | Woody | Temperate | Not endemic |
|  | 91 | Woody | Tropical | Endemic |
|  | 390 | Woody | Tropical | Not endemic |
| **Near Threatened** | 159 |  |  |  |
|  | 2 | Herbaceous | Desert | Endemic |
|  | 1 | Herbaceous | Desert | Not endemic |
|  | 5 | Herbaceous | Subtropical | Endemic |
|  | 10 | Herbaceous | Subtropical | Not endemic |
|  | 8 | Herbaceous | Temperate | Endemic |
|  | 19 | Herbaceous | Temperate | Not endemic |
|  | 5 | Herbaceous | Tropical | Endemic |
|  | 8 | Herbaceous | Tropical | Not endemic |
|  | 5 | Woody | Desert | Endemic |
|  | 5 | Woody | Desert | Not endemic |
|  | 6 | Woody | Subtropical | Endemic |
|  | 14 | Woody | Subtropical | Not endemic |
|  | 3 | Woody | Temperate | Endemic |
|  | 6 | Woody | Temperate | Not endemic |
|  | 25 | Woody | Tropical | Endemic |
|  | 37 | Woody | Tropical | Not endemic |
| **Vulnerable** | 243 |  |  |  |
|  | 1 | Herbaceous | Desert | Endemic |
|  | 2 | Herbaceous | Desert | Not endemic |
|  | 5 | Herbaceous | Subtropical | Endemic |
|  | 11 | Herbaceous | Subtropical | Not endemic |
|  | 11 | Herbaceous | Temperate | Endemic |
|  | 14 | Herbaceous | Temperate | Not endemic |
|  | 10 | Herbaceous | Tropical | Endemic |
|  | 13 | Herbaceous | Tropical | Not endemic |
|  | 4 | Woody | Desert | Endemic |
|  | 4 | Woody | Desert | Not endemic |
|  | 11 | Woody | Subtropical | Endemic |
|  | 9 | Woody | Subtropical | Not endemic |
|  | 13 | Woody | Temperate | Endemic |
|  | 14 | Woody | Temperate | Not endemic |
|  | 60 | Woody | Tropical | Endemic |
|  | 61 | Woody | Tropical | Not endemic |
| **Endangered** | 287 |  |  |  |
|  | 6 | Herbaceous | Desert | Endemic |
|  | N/A | Herbaceous | Desert | Not endemic |
|  | 9 | Herbaceous | Subtropical | Endemic |
|  | 8 | Herbaceous | Subtropical | Not endemic |
|  | 18 | Herbaceous | Temperate | Endemic |
|  | 11 | Herbaceous | Temperate | Not endemic |
|  | 24 | Herbaceous | Tropical | Endemic |
|  | 26 | Herbaceous | Tropical | Not endemic |
|  | 6 | Woody | Desert | Endemic |
|  | 3 | Woody | Desert | Not endemic |
|  | 16 | Woody | Subtropical | Endemic |
|  | 9 | Woody | Subtropical | Not endemic |
|  | 20 | Woody | Temperate | Endemic |
|  | 10 | Woody | Temperate | Not endemic |
|  | 79 | Woody | Tropical | Endemic |
|  | 42 | Woody | Tropical | Not endemic |
| **Critically Endangered** | 166 |  |  |  |
|  | 2 | Herbaceous | Desert | Endemic |
|  | N/A | Herbaceous | Desert | Not endemic |
|  | 6 | Herbaceous | Subtropical | Endemic |
|  | 3 | Herbaceous | Subtropical | Not endemic |
|  | 9 | Herbaceous | Temperate | Endemic |
|  | 4 | Herbaceous | Temperate | Not endemic |
|  | 40 | Herbaceous | Tropical | Endemic |
|  | 6 | Herbaceous | Tropical | Not endemic |
|  | 2 | Woody | Desert | Endemic |
|  | 2 | Woody | Desert | Not endemic |
|  | 12 | Woody | Subtropical | Endemic |
|  | 2 | Woody | Subtropical | Not endemic |
|  | 11 | Woody | Temperate | Endemic |
|  | 6 | Woody | Temperate | Not endemic |
|  | 42 | Woody | Tropical | Endemic |
|  | 19 | Woody | Tropical | Not endemic |
| **Extinct in the Wild** | 7 |  |  |  |
|  | N/A | Herbaceous | Desert | Endemic |
|  | N/A | Herbaceous | Desert | Not endemic |
|  | N/A | Herbaceous | Subtropical | Endemic |
|  | N/A | Herbaceous | Subtropical | Not endemic |
|  | 1 | Herbaceous | Temperate | Endemic |
|  | 1 | Herbaceous | Temperate | Not endemic |
|  | 1 | Herbaceous | Tropical | Endemic |
|  | N/A | Herbaceous | Tropical | Not endemic |
|  | N/A | Woody | Desert | Endemic |
|  | N/A | Woody | Desert | Not endemic |
|  | 2 | Woody | Subtropical | Endemic |
|  | N/A | Woody | Subtropical | Not endemic |
|  | 1 | Woody | Temperate | Endemic |
|  | N/A | Woody | Temperate | Not endemic |
|  | N/A | Woody | Tropical | Endemic |
|  | 1 | Woody | Tropical | Not endemic |
