## Supplementary Table 3 for "Genome size is positively correlated with extinction risk in herbaceous angiosperms"

**Table S3**. Genome size summary statistics for the 3,250-species dataset analysed in 27 phylogenetic logistic regression models and confirmatory phylogenetic path analyses. Estimates are presented across data partitions for life form, climatic zone, endemism, major angiosperm clades, and the four varying threat thresholds used here to model extinction risk as a binary response variable by grouping Red List Categories into non-threatened and threatened groupings. Note that the polarizing threat threshold excluded the Near Threatened category and therefore comprised 150 fewer species than the remaining three thresholds. Abbreviations: LC = Least Concern, NT = Near Threatened, VU = Vulnerable, EN = Endangered, CR = Critically Endangered, EW = Extinct in the Wild.

|  | Number of species | Genome size (Gb/1C) | | | | | | |
| --- | --- | --- | --- | --- | --- | --- | --- | --- |
|  |  | Minimum | Maximum | Mean | Median | Mode | Range  (max.–min.) | Fold change (max./min.) |
| **Genome size dataset** | 3,250 | 0.08 | 73.01 | 3.54 | 1.08 | 0.59 | 72.93 | 924.18 |
| **Extinction risk using point of reference threat threshold** |  |  |  |  |  |  |  |  |
| Non-threatened: LC, NT | 2,547 | 0.08 | 73.01 | 3.11 | 1.01 | 0.69 | 72.93 | 924.18 |
| Threatened: VU, EN, CR, EW | 703 | 0.21 | 69.54 | 5.10 | 1.40 | 0.59 | 69.33 | 324.96 |
| **Extinction risk using lower**  **threat threshold** |  |  |  |  |  |  |  |  |
| Non-threatened: LC | 2,388 | 0.08 | 73.01 | 2.98 | 1.00 | 0.69 | 72.93 | 924.18 |
| Threatened: NT, VU, EN, CR, EW | 862 | 0.21 | 72.81 | 5.09 | 1.45 | 0.59 | 72.60 | 340.25 |
| **Extinction risk using higher**  **threat threshold** |  |  |  |  |  |  |  |  |
| Non-threatened: LC, NT, VU | 2,790 | 0.08 | 73.01 | 3.21 | 1.03 | 0.58 | 72.93 | 924.18 |
| Threatened: EN, CR, EW | 460 | 0.21 | 69.54 | 5.56 | 1.47 | 0.33 | 69.33 | 324.96 |
| **Extinction risk using polarizing threat threshold** |  |  |  |  |  |  |  |  |
| Non-threatened: LC | 2,388 | 0.08 | 73.01 | 2.98 | 1.00 | 0.69 | 72.93 | 924.18 |
| Threatened: VU, EN, CR, EW | 703 | 0.21 | 69.54 | 5.10 | 1.40 | 0.58 | 69.33 | 324.96 |
| **Life form** |  |  |  |  |  |  |  |  |
| Herbaceous | 1,486 | 0.08 | 73.01 | 5.73 | 1.37 | 0.59 | 72.93 | 924.18 |
| Woody | 1,764 | 0.18 | 35.23 | 1.70 | 0.98 | 0.69 | 35.05 | 194.32 |
| **Climate** |  |  |  |  |  |  |  |  |
| Desert | 138 | 0.21 | 23.42 | 4.66 | 1.88 | 1.57 | 23.22 | 114.04 |
| Temperate | 1,330 | 0.10 | 73.01 | 4.51 | 1.12 | 0.39 | 72.91 | 715.78 |
| Subtropical | 505 | 0.08 | 72.81 | 3.68 | 1.27 | 1.47 | 72.73 | 921.70 |
| Tropical | 1,277 | 0.1 | 35.23 | 2.36 | 0.92 | 0.59 | 35.13 | 363.21 |
| **Range size (endemism)** |  |  |  |  |  |  |  |  |
| Endemic | 756 | 0.10 | 68.06 | 3.73 | 1.41 | 1.47 | 67.96 | 654.43 |
| Not endemic | 2,494 | 0.08 | 73.01 | 3.48 | 0.99 | 0.69 | 72.93 | 924.18 |
| **Higher order classification** |  |  |  |  |  |  |  |  |
| Monocots | 1,121 | 0.16 | 73.01 | 7.03 | 2.41 | 0.39 | 72.85 | 455.87 |
| Eudicots  Early diverging angiosperm | 2,033  96 | 0.08  0.39 | 34.73  14.36 | 1.66  3.01 | 0.89  2.15 | 0.69  0.98 | 34.66  13.97 | 439.67  36.63 |
