## Supplementary Table 4 for "Genome size is positively correlated with extinction risk in herbaceous angiosperms"

**Table S4.** Comparative representation of partitions in the 3,250-species dataset assembled here, relative to the 202,753 angiosperms in the World Checklist of Vascular Plants (WCVP) with available life form, endemism, and climate zone information. The proportion of species in each partition is reported for both datasets. The last column indicates the proportional representation of species in our dataset relative to the WCVP, where 1 = same number of species expected in a random sample of 3,250 angiosperms; 2 = double the number of species expected; and 0.5 = half the number of species expected.

| **Life form** | **Endemism** | **Climatic zone** | **No. of species in WCVP** | **% of species in WCVP** | **No. of species in our dataset** | **% of species in our dataset** | **Factor of representation** |
| --- | --- | --- | --- | --- | --- | --- | --- |
| Herbaceous | Endemic | Desert | 1927 | 0.95 | 19 | 0.58 | 0.62 |
| Herbaceous | Endemic | Subtropical | 11149 | 5.50 | 55 | 1.69 | 0.31 |
| Herbaceous | Endemic | Temperate | 13896 | 6.85 | 68 | 2.09 | 0.31 |
| Herbaceous | Endemic | Tropical | 23587 | 11.63 | 106 | 3.26 | 0.28 |
| Herbaceous | Non-endemic | Desert | 2868 | 1.41 | 28 | 0.86 | 0.61 |
| Herbaceous | Non-endemic | Subtropical | 9587 | 4.73 | 178 | 5.48 | 1.16 |
| Herbaceous | Non-endemic | Temperate | 22816 | 11.25 | 708 | 21.78 | 1.94 |
| Herbaceous | Non-endemic | Tropical | 13877 | 6.84 | 324 | 9.97 | 1.46 |
| Woody | Endemic | Desert | 4596 | 2.27 | 30 | 0.92 | 0.41 |
| Woody | Endemic | Subtropical | 11454 | 5.65 | 97 | 2.98 | 0.53 |
| Woody | Endemic | Temperate | 5739 | 2.83 | 84 | 2.58 | 0.91 |
| Woody | Endemic | Tropical | 34282 | 16.91 | 297 | 9.14 | 0.54 |
| Woody | Non-endemic | Desert | 3547 | 1.75 | 61 | 1.88 | 1.07 |
| Woody | Non-endemic | Subtropical | 6790 | 3.35 | 175 | 5.38 | 1.61 |
| Woody | Non-endemic | Temperate | 7328 | 3.61 | 470 | 14.46 | 4 |
| Woody | Non-endemic | Tropical | 29310 | 14.46 | 550 | 16.92 | 1.17 |
