## Supplementary Table 5 for "Genome size is positively correlated with extinction risk in herbaceous angiosperms"

**Table S5.** Description, selection, and phylogenetic signal of the 27 models tested in phylogenetic logistic regressions using three varying threat thresholds that aggregated (a) LC species into a non-threatened grouping and NT, VU, EN, CR, EW species into a threatened grouping; (b) LC, NT, VU species into a non-threatened grouping and EN, CR, EW species into a threatened grouping; and (c) LC species into a non-threatened grouping and VU, EN, CR, EW species into a threatened grouping (i.e., excluding NT species). These threat thresholds were compared to a point of reference, summarized in Table **1**. Models represent all combinations possible when adding life form, climatic zone and endemism as explanatory variables to a baseline model with genome size as the sole predictor. For each threat threshold, models are arranged in decreasing order of best fit based on median ∆AICc values, obtained by first ranking models within each of the 100 variant phylogenies used as input in analyses, and then summarized by model across all trees. Models found to be the best (i.e., ∆AICc = 0) in at least one phylogeny are depicted in Figs. **SXXX**. Mean phylogenetic signal (α) was obtained by summarizing across the 100 separate analyses conducted for each model and then used to calculate –log(α). Abbreviations: ER = extinction risk, GS = genome size, CLIM = climatic zone, LF = life form, ENDEM = endemism (proxy for range size).

_____________________________________________________________________________________________

**Model**	**∆AICc**

**___________________________________________**

**Min.**	**Max.**	**Median**	**Mean**	**Mean**

**α**	–**log(α)**

**_____________________________________________________________________________________________**

**(a)**

1. ER ~ GS + CLIM + LF + GS:LF + ENDEM	0.000	253.180	0.000	7.564	-2.023

2. ER ~ GS + CLIM + LF + GS:LF + ENDEM +	0.000	76.413	1.429	7.863	-2.062

GS:ENDEM

3. ER ~ GS + CLIM + GS:CLIM + LF + GS:LF + ENDEM	0.000	52.513	18.785	7.676	-2.038

4. ER ~ GS + CLIM + GS:CLIM + LF + GS:LF +	0.000	66.642	32.358	7.677	-2.038

ENDEM + GS:ENDEM

5. ER ~ GS + LF + GS:LF + ENDEM + GS:ENDEM	4.383	63.381	40.000	6.494	-1.871

6. ER ~ GS + LF + GS:LF + ENDEM	6.062	74.803	41.230	6.438	-1.862

7. ER ~ GS + CLIM + GS:CLIM + LF + ENDEM	10.024	94.414	57.253	8.291	-2.115

8. ER ~ GS + CLIM + GS:CLIM + ENDEM	16.362	118.726	57.464	7.866	-2.063

9. ER ~ GS + CLIM + GS:CLIM + LF + ENDEM +	13.356	101.958	60.037	8.216	-2.106

GS:ENDEM

10. ER ~ GS + ENDEM + GS:ENDEM + CLIM + GS:CLIM	10.473	86.467	60.167	7.908	-2.068

11. ER ~ GS + CLIM + LF + ENDEM	19.328	123.396	73.029	8.732	-2.167

12. ER ~ GS + CLIM + ENDEM + GS:ENDEM	37.526	112.722	77.543	8.567	-2.148

13. ER ~ GS + CLIM + LF + ENDEM + GS:ENDEM	23.180	151.938	78.668	9.048	-2.203

14. ER ~ GS + CLIM + ENDEM	44.888	114.793	87.782	8.703	-2.164

15. ER ~ GS + LF + ENDEM	76.537	151.826	113.559	7.142	-1.966

16. ER ~ GS + ENDEM	68.918	269.922	116.084	6.962	-1.941

17. ER ~ GS + ENDEM + GS:ENDEM	68.364	154.047	118.101	7.175	-1.971

18. ER ~ GS + LF + ENDEM + GS:ENDEM	83.312	156.006	119.442	7.398	-2.001

19. ER ~ GS + CLIM + LF	287.143	410.742	346.172	8.312	-2.118

20. ER ~ GS + CLIM	284.997	469.334	346.232	8.156	-2.099

21. ER ~ GS + CLIM + GS:CLIM + LF	281.077	405.947	349.183	8.589	-2.151

22. ER ~ CLIM + GS:CLIM	300.303	434.561	359.769	8.815	-2.177

23. ER ~ GS + CLIM + GS:CLIM + LF + GS:LF	300.384	749.828	367.105	9.494	-2.251

24. ER ~ GS + CLIM + LF + GS:LF	307.161	657.863	370.804	10.162	-2.319

25. ER ~ GS + LF	366.000	478.955	432.019	8.028	-2.083

26. ER ~ GS	372.775	506.658	451.243	8.003	-2.080

27. ER ~ GS + LF + GS:LF	387.834	546.206	475.192	9.267	-2.227

**(b)**

1. ER ~ GS + CLIM + LF + GS:LF + ENDEM +	0.000	14.529	0.354	17.333	-2.853

GS:ENDEM

2. ER ~ GS + CLIM + LF + GS:LF + ENDEM	0.000	16.014	1.041	17.220	-2.846

3. ER ~ GS + CLIM + GS:CLIM + LF + GS:LF +	0.000	250.593	1.409	16.612	-2.810

ENDEM + GS:ENDEM

4. ER ~ GS + CLIM + GS:CLIM + LF + GS:LF + ENDEM	0.000	185.449	2.212	16.922	-2.829

5. ER ~ GS + CLIM + GS:CLIM + ENDEM	5.326	25.539	12.202	15.594	-2.747

6. ER ~ GS + CLIM + GS:CLIM + LF + ENDEM	6.950	25.999	12.387	15.344	-2.731

7. ER ~ GS + ENDEM + GS:ENDEM + CLIM + GS:CLIM	6.347	27.531	13.431	15.872	-2.765

8. ER ~ GS + CLIM + GS:CLIM + LF + ENDEM +	6.984	33.688	14.056	15.812	-2.761

GS:ENDEM

9. ER ~ GS + LF + GS:LF + ENDEM + GS:ENDEM	3.286	31.645	14.695	14.711	-2.689

10. ER ~ GS + LF + GS:LF + ENDEM	4.565	32.788	16.476	14.491	-2.674

11. ER ~ GS + CLIM + ENDEM	10.096	34.702	20.474	16.865	-2.825

12. ER ~ GS + CLIM + ENDEM + GS:ENDEM	14.348	35.373	20.497	17.183	-2.844

13. ER ~ GS + CLIM + LF + ENDEM	14.567	35.837	21.361	16.609	-2.810

14. ER ~ GS + CLIM + LF + ENDEM + GS:ENDEM	14.903	36.363	21.568	16.874	-2.826

15. ER ~ GS + LF + ENDEM	20.355	50.554	34.369	14.716	-2.689

16. ER ~ GS + LF + ENDEM + GS:ENDEM	21.898	52.380	35.833	14.777	-2.693

17. ER ~ GS + ENDEM	25.270	66.737	36.310	14.920	-2.703

18. ER ~ GS + ENDEM + GS:ENDEM	26.321	53.384	37.764	14.954	-2.705

19. ER ~ GS + LF	287.664	493.381	325.958	13.561	-2.607

20. ER ~ GS + CLIM + LF + GS:LF	300.577	369.770	326.387	15.240	-2.724

21. ER ~ GS + CLIM + GS:CLIM + LF + GS:LF	297.718	523.336	327.957	14.798	-2.694

22. ER ~ GS + CLIM + GS:CLIM + LF	303.993	380.745	340.322	15.453	-2.738

23. ER ~ GS	303.263	386.426	341.843	14.052	-2.643

24. ER ~ GS + CLIM + LF	267.446	397.439	356.346	17.802	-2.879

25. ER ~ CLIM + GS:CLIM	278.437	499.195	367.838	15.371	-2.733

26. ER ~ GS + CLIM	268.649	419.200	376.938	18.008	-2.891

27. ER ~ GS + LF + GS:LF	289.127	622.996	455.434	15.929	-2.768

**(c)**

1. ER ~ GS + CLIM + GS:CLIM + LF + GS:LF + ENDEM	0.000	43.139	1.551	6.966	-1.941

2. ER ~ GS + CLIM + LF + GS:LF + ENDEM	0.000	32.048	2.836	7.401	-2.002

3. ER ~ GS + CLIM + LF + GS:LF + ENDEM +	0.000	94.881	3.254	7.308	-1.989

GS:ENDEM

4. ER ~ GS + CLIM + GS:CLIM + LF + GS:LF +	0.000	185.747	4.067	6.639	-1.893

ENDEM + GS:ENDEM

5. ER ~ GS + LF + GS:LF + ENDEM + GS:ENDEM	2.656	61.610	27.636	6.545	-1.879

6. ER ~ GS + CLIM + GS:CLIM + ENDEM	2.288	188.408	28.687	7.816	-2.056

7. ER ~ GS + LF + GS:LF + ENDEM	3.329	72.280	29.077	6.600	-1.887

8. ER ~ GS + CLIM + GS:CLIM + LF + ENDEM +	7.877	80.838	30.541	8.026	-2.083

GS:ENDEM

9. ER ~ GS + CLIM + GS:CLIM + LF + ENDEM	8.752	146.192	30.886	8.077	-2.089

10. ER ~ GS + ENDEM + GS:ENDEM + CLIM + GS:CLIM	10.448	237.854	32.531	7.835	-2.059

11. ER ~ GS + CLIM + ENDEM + GS:ENDEM	17.099	124.298	43.615	7.938	-2.072

12. ER ~ GS + CLIM + LF + ENDEM + GS:ENDEM	16.923	88.981	44.217	8.316	-2.118

13. ER ~ GS + CLIM + ENDEM	20.274	107.318	46.186	8.491	-2.139

14. ER ~ GS + CLIM + LF + ENDEM	24.738	90.841	46.370	8.481	-2.138

15. ER ~ GS + LF + ENDEM	48.662	131.451	81.269	8.191	-2.103

16. ER ~ GS + LF + ENDEM + GS:ENDEM	52.958	124.423	82.874	7.978	-2.077

17. ER ~ GS + ENDEM + GS:ENDEM	59.382	190.848	87.550	7.062	-1.955

18. ER ~ GS + ENDEM	58.357	141.675	88.743	7.606	-2.029

19. ER ~ GS + CLIM	304.069	410.422	339.568	8.385	-2.126

20. ER ~ GS + CLIM + LF	302.530	519.426	340.352	8.322	-2.119

21. ER ~ GS + CLIM + LF + GS:LF	317.177	412.465	356.743	9.468	-2.248

22. ER ~ GS + CLIM + GS:CLIM + LF + GS:LF	315.692	410.125	357.362	9.022	-2.200

23. ER ~ GS + CLIM + GS:CLIM + LF	298.849	558.229	367.280	9.060	-2.204

24. ER ~ CLIM + GS:CLIM	301.659	407.307	371.252	8.971	-2.194

25. ER ~ GS + LF	369.359	569.246	403.902	7.764	-2.049

26. ER ~ GS	395.108	480.857	439.739	8.398	-2.128

27. ER ~ GS + LF + GS:LF	367.711	518.348	441.625	8.654	-2.158
