## Supplementary Table 6 for "Genome size is positively correlated with extinction risk in herbaceous angiosperms"

**Table S6.** Consensus coefficients for best models identified in phylogenetic logistic regressions based on four varying threat thresholds to dichotomize extinction risk, showing (a, b) two retained best models when using the Red List definition of threat as a threshold (shown in Fig. **2**), (c) one retained best model when using a lower threshold (shown in Fig. **S19**), (d-f) three retained best models when using a higher threshold (shown in Figs. **S19–S21**), and (g-i) three retained best models when using a polarizing threshold that excludes Near Threatened species (shown in Figs. **S19–S21**). Values represent mean coefficients and 95% confidence intervals (from bootstrapping) across the 100 separate analyses conducted for each model using different phylogenies as input. Categories assigned as the reference for each of the categorical and binary explanatory variables are summarized in the intercept estimates. Abbreviations: ER = extinction risk, GS = genome size, CLIM = climatic zone, LF = life form, ENDEM = endemism (proxy for range size).

| **(a) Model: ER ~ GS + CLIM + GS:CLIM + LF + GS:LF + ENDEM + GS:ENDEM** | | | | |
| --- | --- | --- | --- | --- |
|  | **Model terms** | **Category** | **Estimate** | **95% confidence intervals** |
| **Intercept** |  | Desert, herbaceous, endemic | -1.272 | -1.494, -0.988 |
| **Slope** | GS | Desert, herbaceous, endemic | 1.927 | 1.563, 2.141 |
|  | GS:CLIM | Subtropical | 0.075 | -0.241, 0.439 |
|  |  | Tropical | -0.596 | -0.854, -0.247 |
|  |  | Temperate | 0.036 | -0.249, 0.388 |
|  | GS:LF | Woody | -1.712 | -2.043, -1.305 |
|  | GS:ENDEM | Not endemic | -0.365 | -0.671, 0.002 |
| **Additive** | CLIM | Subtropical | 0.116 | -0.165, 0.382 |
|  |  | Tropical | 0.985 | 0.75, 1.203 |
|  |  | Temperate | -0.462 | -0.746, -0.215 |
|  | LF | Woody | 0.831 | 0.579, 1.051 |
|  | ENDEM | Not endemic | -1.69 | -1.897, -1.502 |
| **(b) Model: ER ~ GS + CLIM + GS:CLIM + LF + GS:LF + ENDEM** | | | | |
| **Intercept** |  | Desert, herbaceous, endemic | -1.246 | -1.487, -0.979 |
| **Slope** | GS | Desert, herbaceous, endemic | 1.794 | 1.499, 1.97 |
|  | GS:CLIM | Subtropical | -0.01 | -0.314, 0.356 |
|  |  | Tropical | -0.666 | -0.909, -0.303 |
|  |  | Temperate | -0.068 | -0.33, 0.287 |
|  | GS:LF | Woody | -1.648 | -1.992, -1.267 |
| **Additive** | CLIM | Subtropical | 0.072 | -0.198, 0.351 |
|  |  | Tropical | 0.989 | 0.763, 1.212 |
|  |  | Temperate | -0.514 | -0.795, -0.257 |
|  | LF | Woody | 0.843 | 0.601, 1.066 |
|  | ENDEM | Not endemic | -1.728 | -1.914, -1.541 |
| **(c) Model: ER ~ GS + CLIM + LF + GS:LF + ENDEM** | | | |  |
| **Intercept** |  | Desert, herbaceous, endemic | -0.406 | -0.708, -0.178 |
| **Slope** | GS | Desert, herbaceous, endemic | 1.73 | 1.51,1.98 |
|  | GS:LF | Woody | -1.98 | -2.36, -1.61 |
| **Additive** | CLIM | Subtropical | -0.0741 | -0.297, 0.216 |
|  |  | Tropical | 0.296 | 0.0971, 0.563 |
|  |  | Temperate | -0.597 | -0.808, -0.324 |
|  | LF | Woody | 0.504 | 0.279, 0.743 |
|  | ENDEM | Not endemic | -1.91 | -2.11, -1.71 |
| **(d) Model: ER ~ GS + CLIM + LF + GS:LF + ENDEM** | | | |  |
| **Intercept** |  | Desert, herbaceous, endemic | -1.43 | -1.76, -1.07 |
| **Slope** | GS | Desert, herbaceous, endemic | 1.31 | 0.991, 1.59 |
|  | GS:LF | Woody | -1.17 | -1.61, -0.646 |
| **Additive** | CLIM | Subtropical | 0.161 | -0.209, 0.503 |
|  |  | Tropical | 0.628 | 0.306, 0.936 |
|  |  | Temperate | -0.172 | -0.533, 0.158 |
|  | LF | Woody | 0.435 | 0.107, 0.72 |
|  | ENDEM | Not endemic | -2.17 | -2.42, -1.93 |
| **(e) Model: ER ~ GS + CLIM + GS:CLIM + LF + GS:LF + ENDEM** | | | | |
| **Intercept** |  | Desert, herbaceous, endemic | -1.79 | -2.1, -1.45 |
| **Slope** | GS | Desert, herbaceous, endemic | 1.88 | 1.57, 2.17 |
|  | GS:CLIM | Subtropical | -0.794 | -1.25, -0.346 |
|  |  | Tropical | -0.596 | -0.977, -0.189 |
|  |  | Temperate | -0.468 | -0.874, -0.0793 |
|  | GS:LF | Woody | -1.19 | -1.65, -0.675 |
| **Additive** | CLIM | Subtropical | 0.577 | 0.208, 0.931 |
|  |  | Tropical | 1 | 0.695, 1.3 |
|  |  | Temperate | 0.129 | -0.242, 0.473 |
|  | LF | Woody | 0.435 | 0.122, 0.727 |
|  | ENDEM | Not endemic | -2.16 | -2.41, -1.92 |
| **(f) Model: ER ~ GS + CLIM + LF + GS:LF + ENDEM + GS:ENDEM** | | | | |
| **Intercept** |  | Desert, herbaceous, endemic | -1.51 | -1.83, -1.15 |
| **Slope** | GS | Desert, herbaceous, endemic | 1.52 | 1.15, 1.86 |
|  | GS:LF | Woody | -1.23 | -1.67, -0.733 |
|  | GS:ENDEM | Not endemic | -0.304 | -0.732, 0.101 |
| **Additive** | CLIM | Subtropical | 0.199 | -0.171, 0.542 |
|  |  | Tropical | 0.655 | 0.333, 0.96 |
|  |  | Temperate | -0.106 | -0.469, 0.22 |
|  | LF | Woody | 0.433 | 0.118, 0.723 |
|  | ENDEM | Not endemic | -2.11 | -2.39, -1.84 |
| **(g) Model: ER ~ GS + CLIM + LF + GS:LF + ENDEM** | | | |  |
| **Intercept** |  | Desert, herbaceous, endemic | -1.01 | -1.26, -0.743 |
| **Slope** | GS | Desert, herbaceous, endemic | 1.75 | 1.49, 1.97 |
|  | GS:LF | Woody | -1.9 | -2.23, -1.53 |
| **Additive** | CLIM | Subtropical | -0.00309 | -0.265, 0.254 |
|  |  | Tropical | 0.761 | 0.528, 0.989 |
|  |  | Temperate | -0.644 | -0.896, -0.395 |
|  | LF | Woody | 0.897 | 0.647, 1.11 |
|  | ENDEM | Not endemic | -1.8 | -1.99, -1.61 |
| **(h) Model: ER ~ GS + CLIM + GS:CLIM + LF + GS:LF + ENDEM** | | | | |
| **Intercept** |  | Desert, herbaceous, endemic | -0.994 | -1.22, -0.693 |
| **Slope** | GS | Desert, herbaceous, endemic | 1.77 | 1.47, 1.93 |
|  | GS:CLIM | Subtropical | 0.195 | -0.0868, 0.593 |
|  |  | Tropical | -0.522 | -0.744, -0.14 |
|  |  | Temperate | -0.00961 | -0.223, 0.361 |
|  | GS:LF | Woody | -1.84 | -2.19, -1.46 |
| **Additive** | CLIM | Subtropical | -0.0448 | -0.349, 0.202 |
|  |  | Tropical | 0.888 | 0.626, 1.09 |
|  |  | Temperate | -0.536 | -0.843, -0.307 |
|  | LF | Woody | 0.795 | 0.552, 1.03 |
|  | ENDEM | Not endemic | -1.91 | -2.1, -1.71 |
| **(i) Model: ER ~ GS + CLIM + LF + GS:LF + ENDEM + GS:ENDEM** | | | | |
| **Intercept** |  | Desert, herbaceous, endemic | -1.05 | -1.29, -0.754 |
| **Slope** | GS | Desert, herbaceous, endemic | 1.99 | 1.61, 2.29 |
|  | GS:LF | Woody | -2.07 | -2.41, -1.63 |
|  | GS:ENDEM | Not endemic | -0.35 | -0.645, 0.0252 |
| **Additive** | CLIM | Subtropical | 0.0613 | -0.219, 0.316 |
|  |  | Tropical | 0.786 | 0.538, 1.02 |
|  |  | Temperate | -0.57 | -0.839, -0.322 |
|  | LF | Woody | 0.85 | 0.591, 1.07 |
|  | ENDEM | Not endemic | -1.77 | -1.98, -1.58 |
