## Supplementary Table 8 for "Genome size is positively correlated with extinction risk in herbaceous angiosperms"

**Table S8.** List of angiosperms with a potentially heightened risk of extinction, focusing on large-genomed herbaceous species that lack an extinction risk assessment in the current release of the Red List (IUCN 2022); additional information on endemism (proxy for range size) and climatic niche is provided. A family-level summary of these species is depicted in Fig. **5**. Species are arranged alphabetically across and within families.

| **Species name** | **1C-value (Gb)** | **Endemism (range size)** | **Climate zone** |
| --- | --- | --- | --- |
| **Alismataceae** |  |  |  |
| *Aquarius floribundus* | 12.838 | Not endemic | Tropical |
| *Aquarius macrophyllus* | 12.838 | Not endemic | Tropical |
| *Aquarius paniculatus* | 13.426 | Not endemic | Tropical |
| *Aquarius subulatus* | 14.308 | Not endemic | Tropical |
| *Aquarius uruguayensis* | 14.014 | Not endemic | Subtropical |
| *Sagittaria lancifolia* | 13.034 | Not endemic | Tropical |
| **Alstroemeriaceae** | |  |  |
| *Alstroemeria angustifolia* | 25.382 | Not endemic | Subtropical |
| *Alstroemeria aurea* | 20.681 | Not endemic | Temperate |
| *Alstroemeria brasiliensis* | 24.598 | Endemic | Tropical |
| *Alstroemeria caryophyllaea* | 27.636 | Endemic | Subtropical |
| *Alstroemeria hookeri* | 25.428 | Not endemic | Subtropical |
| *Alstroemeria inodora* | 24.794 | Not endemic | Tropical |
| *Alstroemeria ligtu* | 38.631 | Not endemic | Subtropical |
| *Alstroemeria magnifica* | 17.934 | Endemic | Subtropical |
| *Alstroemeria pelegrina* | 20.8415 | Not endemic | Subtropical |
| *Alstroemeria philippii* | 20.384 | Endemic | Desert |
| *Alstroemeria psittacina* | 24.5 | Not endemic | Subtropical |
| *Alstroemeria pulchra* | 19.609 | Endemic | Subtropical |
| **Amaryllidaceae** | |  |  |
| *Agapanthus coddii* | 11.858 | Endemic | Subtropical |
| *Agapanthus praecox* | 12.234 | Not endemic | Subtropical |
| *Allium aflatunense* | 21.4179 | Not endemic | Temperate |
| *Allium akaka* | 13.328 | Endemic | Temperate |
| *Allium alexeianum* | 15.4987 | Not endemic | Temperate |
| *Allium altissimum* | 21.5061 | Not endemic | Temperate |
| *Allium angulosum* | 14.798 | Not endemic | Temperate |
| *Allium aroides* | 18.6641 | Not endemic | Temperate |
| *Allium atropurpureum* | 25.5976 | Not endemic | Temperate |
| *Allium backhousianum* | 21.707 | Endemic | Temperate |
| *Allium bakhtiaricum* | 20.2272 | Endemic | Temperate |
| *Allium barsczewskii* | 15.484 | Not endemic | Temperate |
| *Allium breviscapum* | 22.5155 | Endemic | Temperate |
| *Allium bucharicum* | 13.916 | Not endemic | Temperate |
| *Allium caesium* | 12.642 | Not endemic | Temperate |
| *Allium canadense* | 43.218 | Not endemic | Temperate |
| *Allium cardiostemon* | 19.0267 | Not endemic | Temperate |
| *Allium carolinianum* | 12.936 | Not endemic | Temperate |
| *Allium caspium* | 15.582 | Not endemic | Temperate |
| *Allium cepa* | 17.542 | Endemic | Temperate |
| *Allium cernuum* | 16.758 | Not endemic | Temperate |
| *Allium chitralicum* | 33.6581 | Not endemic | Temperate |
| *Allium chloranthum* | 15.288 | Endemic | Subtropical |
| *Allium chychkanense* | 16.6502 | Endemic | Temperate |
| *Allium clathratum* | 11.956 | Not endemic | Temperate |
| *Allium coloratum* | 17.726 | Not endemic | Temperate |
| *Allium costatovaginatum* | 16.8854 | Not endemic | Temperate |
| *Allium cristophii* | 22.5204 | Not endemic | Temperate |
| *Allium cupani* | 23.422 | Not endemic | Temperate |
| *Allium cupuliferum* | 20.0067 | Not endemic | Temperate |
| *Allium cyrilli* | 19.6 | Not endemic | Subtropical |
| *Allium darwasicum* | 16.611 | Not endemic | Temperate |
| *Allium delicatulum* | 13.622 | Not endemic | Temperate |
| *Allium denudatum* | 13.818 | Not endemic | Temperate |
| *Allium derderianum* | 21.2758 | Endemic | Temperate |
| *Allium dictyoprasum* | 16.562 | Not endemic | Temperate |
| *Allium dodecadontum* | 18.3309 | Endemic | Temperate |
| *Allium drummondii* | 22.246 | Not endemic | Temperate |
| *Allium elburzense* | 25.1174 | Endemic | Temperate |
| *Allium ellisii* | 21.9177 | Endemic | Temperate |
| *Allium ericetorum* | 15.778 | Not endemic | Temperate |
| *Allium farreri* | 27.538 | Not endemic | Temperate |
| *Allium fetisowii* | 12.8674 | Not endemic | Temperate |
| *Allium filidens* | 24.206 | Not endemic | Temperate |
| *Allium flavellum* | 11.858 | Not endemic | Temperate |
| *Allium flavum* | 11.9691711 | Not endemic | Temperate |
| *Allium fuscoviolaceum* | 20.188 | Not endemic | Temperate |
| *Allium galanthum* | 11.956 | Not endemic | Temperate |
| *Allium giganteum* | 20.1684 | Not endemic | Temperate |
| *Allium gomphrenoides* | 17.052 | Endemic | Subtropical |
| *Allium grande* | 19.4726 | Not endemic | Temperate |
| *Allium gunibicum* | 15.288 | Endemic | Temperate |
| *Allium gypsaceum* | 16.5718 | Not endemic | Temperate |
| *Allium hamedanense* | 22.6772 | Endemic | Temperate |
| *Allium heldreichii* | 15.778 | Endemic | Temperate |
| *Allium hissaricum* | 14.406 | Endemic | Temperate |
| *Allium hollandicum* | 21.5306 | Endemic | Temperate |
| *Allium hookeri* | 15.484 | Not endemic | Temperate |
| *Allium hymenorhizum* | 12.544 | Not endemic | Temperate |
| *Allium iliense* | 17.542 | Endemic | Temperate |
| *Allium insubricum* | 20.678 | Endemic | Temperate |
| *Allium intradarvazicum* | 18.5808 | Endemic | Temperate |
| *Allium isakulii* | 20.0018 | Not endemic | Temperate |
| *Allium jesdianum* | 11.76 | Not endemic | Temperate |
| *Allium junceum* | 13.328 | Not endemic | Subtropical |
| *Allium karataviense* | 19.4432 | Not endemic | Temperate |
| *Allium karsianum* | 14.014 | Not endemic | Temperate |
| *Allium koelzii* | 26.9059 | Endemic | Temperate |
| *Allium komarowii* | 19.9234 | Not endemic | Temperate |
| *Allium kuhsorkhense* | 22.7605 | Endemic | Temperate |
| *Allium kunthianum* | 17.248 | Not endemic | Temperate |
| *Allium kurssanovii* | 15.68 | Not endemic | Temperate |
| *Allium leucocephalum* | 12.936 | Not endemic | Temperate |
| *Allium lineare* | 12.838 | Not endemic | Temperate |
| *Allium lipskyanum* | 16.4738 | Not endemic | Temperate |
| *Allium longifolium* | 21.952 | Not endemic | Subtropical |
| *Allium longiradiatum* | 15.386 | Not endemic | Temperate |
| *Allium lusitanicum* | 21.3769943 | Not endemic | Temperate |
| *Allium macleanii* | 20.8887 | Not endemic | Temperate |
| *Allium macrochaetum* | 13.839 | Not endemic | Temperate |
| *Allium mairei* | 13.818 | Not endemic | Temperate |
| *Allium majus* | 17.9144 | Endemic | Temperate |
| *Allium margaritae* | 15.484 | Not endemic | Temperate |
| *Allium materculae* | 20.9426 | Not endemic | Temperate |
| *Allium montibaicalense* | 14.994 | Endemic | Temperate |
| *Allium moschatum* | 23.422 | Not endemic | Temperate |
| *Allium narcissiflorum* | 21.168 | Not endemic | Temperate |
| *Allium neapolitanum* | 15.288 | Not endemic | Subtropical |
| *Allium neriniflorum* | 21.756 | Not endemic | Temperate |
| *Allium nevskianum* | 17.934 | Not endemic | Temperate |
| *Allium nigrum* | 26.8569 | Not endemic | Temperate |
| *Allium nutans* | 22.148 | Not endemic | Temperate |
| *Allium obliquum* | 12.936 | Not endemic | Temperate |
| *Allium ochotense* | 40.964 | Not endemic | Temperate |
| *Allium oleraceum* | 20.8201 | Not endemic | Temperate |
| *Allium oreophilum* | 19.012 | Not endemic | Temperate |
| *Allium orientale* | 18.175 | Not endemic | Temperate |
| *Allium oschaninii* | 17.836 | Not endemic | Temperate |
| *Allium paradoxum* | 26.264 | Not endemic | Temperate |
| *Allium protensum* | 15.9201 | Not endemic | Temperate |
| *Allium pseudobodeanum* | 25.3232 | Endemic | Temperate |
| *Allium pseudotelmatum* | 31.1493 | Endemic | Temperate |
| *Allium pskemense* | 17.248 | Not endemic | Temperate |
| *Allium regelii* | 20.9867 | Not endemic | Temperate |
| *Allium robustum* | 15.7486 | Not endemic | Temperate |
| *Allium rosenbachianum* | 20.18592 | Not endemic | Temperate |
| *Allium rosenorum* | 18.9532 | Not endemic | Temperate |
| *Allium rotundum* | 15.288 | Not endemic | Temperate |
| *Allium rupestre* | 18.522 | Not endemic | Temperate |
| *Allium saposhnikovii* | 29.4735 | Not endemic | Temperate |
| *Allium saralicum* | 20.972 | Not endemic | Temperate |
| *Allium sarawschanicum* | 16.758 | Not endemic | Temperate |
| *Allium sardoum* | 21.462 | Not endemic | Subtropical |
| *Allium sativum* | 15.876 | Not endemic | Temperate |
| *Allium saxatile* | 13.524 | Not endemic | Temperate |
| *Allium schachimardanicum* | 16.6306 | Not endemic | Temperate |
| *Allium schubertii* | 15.484 | Not endemic | Subtropical |
| *Allium scorodoprasum* | 23.128 | Not endemic | Temperate |
| *Allium senescens* | 21.168 | Not endemic | Temperate |
| *Allium severtzovioides* | 15.7241 | Not endemic | Temperate |
| *Allium sewerzowii* | 27.6605 | Not endemic | Temperate |
| *Allium shelkovnikovii* | 19.3207 | Endemic | Temperate |
| *Allium siculum* | 34.7116 | Not endemic | Temperate |
| *Allium splendens* | 23.814 | Not endemic | Temperate |
| *Allium stellatum* | 21.56 | Not endemic | Temperate |
| *Allium stipitatum* | 21.0406 | Not endemic | Temperate |
| *Allium subhirsutum* | 17.444 | Not endemic | Subtropical |
| *Allium subvillosum* | 42.336 | Not endemic | Subtropical |
| *Allium suworowii* | 18.4338 | Not endemic | Temperate |
| *Allium taeniopetalum* | 15.0038 | Not endemic | Temperate |
| *Allium tenuissimum* | 12.74 | Not endemic | Temperate |
| *Allium thunbergii* | 25.872 | Not endemic | Temperate |
| *Allium trautvetterianum* | 20.2468 | Endemic | Temperate |
| *Allium tricoccum* | 34.1922 | Not endemic | Temperate |
| *Allium trifoliatum* | 15.974 | Not endemic | Subtropical |
| *Allium tschimganicum* | 17.8654 | Not endemic | Temperate |
| *Allium tuberosum* | 31.4482 | Not endemic | Temperate |
| *Allium tulipifolium* | 15.7584 | Not endemic | Temperate |
| *Allium tuncelianum* | 13.818 | Endemic | Temperate |
| *Allium tuvinicum* | 24.598 | Not endemic | Temperate |
| *Allium ubipetrense* | 21.3346 | Endemic | Temperate |
| *Allium umbilicatum* | 12.936 | Not endemic | Temperate |
| *Allium unifolium* | 15.974 | Not endemic | Temperate |
| *Allium verticillatum* | 16.4101 | Not endemic | Temperate |
| *Allium victorialis* | 18.62 | Not endemic | Temperate |
| *Allium vineale* | 19.1425 | Not endemic | Temperate |
| *Allium vvedenskyanum* | 16.3856 | Not endemic | Temperate |
| *Allium wallichii* | 15.876 | Not endemic | Temperate |
| *Allium winklerianum* | 16.1063 | Not endemic | Temperate |
| *Allium zebdanense* | 12.348 | Not endemic | Temperate |
| *Amaryllis belladonna* | 18.7621 | Endemic | Subtropical |
| *Apodolirion lanceolatum* | 22.055 | Endemic | Subtropical |
| *Apodolirion macowanii* | 20.505 | Endemic | Subtropical |
| *Brunsvigia josephinae* | 14.365 | Endemic | Subtropical |
| *Clivia gardenii* | 18.326 | Not endemic | Desert |
| *Clivia miniata* | 17.248 | Not endemic | Subtropical |
| *Clivia nobilis* | 16.8952 | Endemic | Subtropical |
| *Crinum asiaticum* | 24.695 | Not endemic | Tropical |
| *Crinum bulbispermum* | 19.698 | Not endemic | Subtropical |
| *Crinum moorei* | 27.0255 | Not endemic | Subtropical |
| *Cryptostephanus haemanthoides* | 25.2497 | Not endemic | Tropical |
| *Cyrtanthus elatus* | 14.455 | Endemic | Subtropical |
| *Cyrtanthus montanus* | 13.545 | Endemic | Subtropical |
| *Galanthus alpinus* | 30.478 | Not endemic | Temperate |
| *Galanthus cilicicus* | 32.193 | Endemic | Temperate |
| *Galanthus elwesii* | 27.097 | Not endemic | Temperate |
| *Galanthus fosteri* | 26.95 | Not endemic | Temperate |
| *Galanthus gracilis* | 26.803 | Not endemic | Temperate |
| *Galanthus krasnovii* | 44.247 | Not endemic | Temperate |
| *Galanthus lagodechianus* | 80.507 | Not endemic | Temperate |
| *Galanthus platyphyllus* | 44.296 | Not endemic | Temperate |
| *Galanthus rizehensis* | 23.765 | Not endemic | Temperate |
| *Galanthus samothracicus* | 25.43126 | Endemic | Temperate |
| *Galanthus transcaucasicus* | 40.131 | Not endemic | Temperate |
| *Galanthus woronowii* | 27.587 | Not endemic | Temperate |
| *Gethyllis villosa* | 22.344 | Endemic | Subtropical |
| *Gilliesia atropurpurea* | 19.3155 | Endemic | Subtropical |
| *Gilliesia graminea* | 18.71892 | Not endemic | Subtropical |
| *Gilliesia miersioides* | 18.00498 | Endemic | Subtropical |
| *Gilliesia montana* | 19.4622 | Endemic | Subtropical |
| *Haemanthus albiflos* | 32.5565 | Not endemic | Subtropical |
| *Haemanthus coccineus* | 22.16 | Not endemic | Subtropical |
| *Haemanthus sanguineus* | 25.815 | Endemic | Subtropical |
| *Hippeastrum argentinum* | 15.0332 | Not endemic | Subtropical |
| *Hippeastrum aulicum* | 16.2288 | Not endemic | Tropical |
| *Hippeastrum blossfeldiae* | 22.5596 | Not endemic | Tropical |
| *Hippeastrum correiense* | 14.2345 | Endemic | Tropical |
| *Hippeastrum cybister* | 27.6115 | Not endemic | Desert |
| *Hippeastrum elegans* | 16.3415 | Not endemic | Tropical |
| *Hippeastrum evansiae* | 15.6408 | Endemic | Subtropical |
| *Hippeastrum machupijchense* | 16.7482 | Endemic | Subtropical |
| *Hippeastrum morelianum* | 13.1761 | Endemic | Tropical |
| *Hippeastrum papilio* | 14.23 | Endemic | Subtropical |
| *Hippeastrum parodii* | 14.7 | Not endemic | Subtropical |
| *Hippeastrum psittacinum* | 15.3517 | Not endemic | Tropical |
| *Hippeastrum puniceum* | 18.9581 | Not endemic | Tropical |
| *Hippeastrum reginae* | 25.8671 | Not endemic | Tropical |
| *Hippeastrum reticulatum* | 14.063 | Not endemic | Tropical |
| *Hippeastrum scopulorum* | 28.7679 | Endemic | Subtropical |
| *Hippeastrum striatum* | 13.7102 | Not endemic | Tropical |
| *Ipheion sessile* | 20.33262 | Not endemic | Subtropical |
| *Latace andina* | 18.25437 | Not endemic | Subtropical |
| *Leucocoryne appendiculata* | 26.88522 | Endemic | Desert |
| *Leucocoryne coquimbensis* | 27.25686 | Endemic | Subtropical |
| *Leucocoryne incrassata* | 50.83644 | Endemic | Desert |
| *Leucocoryne ixioides* | 55.80468 | Endemic | Subtropical |
| *Leucocoryne macropetala* | 57.59442 | Not endemic | Desert |
| *Leucocoryne narcissoides* | 51.30588 | Endemic | Desert |
| *Leucocoryne odorata* | 54.97338 | Endemic | Subtropical |
| *Leucocoryne pauciflora* | 27.21774 | Endemic | Subtropical |
| *Leucocoryne purpurea* | 42.16158 | Endemic | Subtropical |
| *Leucocoryne vittata* | 28.3131 | Endemic | Subtropical |
| *Lycoris aurea* | 23.961 | Not endemic | Subtropical |
| *Lycoris chinensis* | 31.682 | Not endemic | Temperate |
| *Lycoris haywardii* | 23.416 | Not endemic | Subtropical |
| *Lycoris longituba* | 30.675 | Endemic | Subtropical |
| *Lycoris radiata* | 19.755 | Not endemic | Subtropical |
| *Lycoris sprengeri* | 23.8 | Not endemic | Subtropical |
| *Lycoris straminea* | 26.354 | Endemic | Subtropical |
| *Miersia chilensis* | 25.87788 | Endemic | Subtropical |
| *Miersia humilis* | 30.38646 | Endemic | Subtropical |
| *Narcissus abscissus* | 12.936 | Not endemic | Temperate |
| *Narcissus broussonetii* | 18.326 | Endemic | Subtropical |
| *Narcissus cuneiflorus* | 11.858 | Not endemic | Temperate |
| *Narcissus flavus* | 25.088 | Not endemic | Temperate |
| *Narcissus hispanicus* | 12.642 | Not endemic | Temperate |
| *Narcissus jonquilla* | 16.072 | Not endemic | Temperate |
| *Narcissus moleroi* | 12.789 | Endemic | Temperate |
| *Narcissus papyraceus* | 16.513 | Not endemic | Subtropical |
| *Narcissus romieuxii* | 14.112 | Endemic | Temperate |
| *Narcissus rupicola* | 13.034 | Not endemic | Temperate |
| *Narcissus tazetta* | 14.847 | Not endemic | Subtropical |
| *Nerine bowdenii* | 17.297 | Not endemic | Subtropical |
| *Nerine gracilis* | 12.054 | Not endemic | Subtropical |
| *Nerine humilis* | 12.397 | Endemic | Subtropical |
| *Nerine krigei* | 15.876 | Not endemic | Subtropical |
| *Nerine laticoma* | 13.105 | Not endemic | Desert |
| *Nerine pudica* | 12.838 | Endemic | Subtropical |
| *Nerine pusilla* | 13.72 | Endemic | Desert |
| *Nerine ridleyi* | 12.838 | Endemic | Subtropical |
| *Nerine sarniensis* | 12.691 | Endemic | Subtropical |
| *Nerine undulata* | 13.916 | Endemic | Subtropical |
| *Nothoscordum andicola* | 31.57962 | Not endemic | Subtropical |
| *Nothoscordum bivalve* | 13.50129 | Not endemic | Temperate |
| *Nothoscordum bonariense* | 34.70433 | Not endemic | Subtropical |
| *Nothoscordum dialystemon* | 27.90723 | Not endemic | Subtropical |
| *Nothoscordum felipponei* | 15.07098 | Not endemic | Subtropical |
| *Nothoscordum gaudichaudianum* | 14.53797 | Not endemic | Subtropical |
| *Nothoscordum gracile* | 41.3415 | Not endemic | Subtropical |
| *Nothoscordum hirtellum* | 17.54043 | Not endemic | Subtropical |
| *Nothoscordum montevidense* | 45.92688 | Not endemic | Subtropical |
| *Nothoscordum nudicaule* | 23.03 | Not endemic | Temperate |
| *Nothoscordum vittatum* | 14.28858 | Not endemic | Subtropical |
| *Pancratium maritimum* | 24.3334 | Not endemic | Subtropical |
| *Phycella scarlatina* | 14.9634 | Endemic | Subtropical |
| *Scadoxus multiflorus* | 43.316 | Not endemic | Subtropical |
| *Sprekelia formosissima* | 64.1365 | Not endemic | Tropical |
| *Sternbergia candida* | 30.31311 | Endemic | Temperate |
| *Sternbergia clusiana* | 31.64808 | Not endemic | Temperate |
| *Sternbergia colchiciflora* | 19.5118 | Not endemic | Temperate |
| *Sternbergia lutea* | 23.481 | Not endemic | Temperate |
| *Sternbergia vernalis* | 30.22998 | Not endemic | Temperate |
| *Tristagma bivalve* | 18.90474 | Endemic | Subtropical |
| *Tristagma circinatum* | 14.94384 | Not endemic | Temperate |
| *Tristagma gracile* | 16.90962 | Endemic | Subtropical |
| *Tristagma graminifolium* | 17.38395 | Endemic | Temperate |
| *Tristagma nivale* | 16.39617 | Not endemic | Temperate |
| *Tristagma patagonicum* | 16.21524 | Not endemic | Temperate |
| *Tristagma porrifolium* | 34.67988 | Endemic | Subtropical |
| *Tristagma violaceum* | 32.50872 | Endemic | Subtropical |
| *Tulbaghia simmleri* | 19.01232 | Endemic | Subtropical |
| *Tulbaghia violacea* | 18.155 | Not endemic | Subtropical |
| *Urceolina amazonica* | 11.984 | Endemic | Tropical |
| *Zephyranthes candida* | 18.62 | Not endemic | Subtropical |
| *Zephyranthes citrina* | 15.435 | Not endemic | Subtropical |
| *Zephyranthes minuta* | 14.41083 | Not endemic | NA |
| **Araceae** |  |  |  |
| *Amorphophallus johnsonii* | 15.484 | Not endemic | Tropical |
| *Anthurium grande* | 13.25205 | Endemic | Tropical |
| *Arisarum vulgare* | 14.2665 | Not endemic | Subtropical |
| *Arum italicum* | 12.3815 | Not endemic | Temperate |
| *Orontium aquaticum* | 14.7 | Not endemic | Temperate |
| *Zamioculcas zamiifolia* | 23.569 | Not endemic | Tropical |
| **Asparagaceae** |  |  |  |
| *Albuca concordiana* | 17.64 | Not endemic | Subtropical |
| *Aspidistra elatior* | 17.542 | Endemic | Subtropical |
| *Bellevalia desertorum* | 24.001 | Not endemic | Subtropical |
| *Bellevalia edirnensis* | 26.066145 | Not endemic | Subtropical |
| *Bellevalia longistyla* | 32.046 | Not endemic | Temperate |
| *Bellevalia sitiaca* | 19.15413 | Endemic | Subtropical |
| *Bellevalia tauri* | 13.328 | Endemic | Subtropical |
| *Bellevalia trifoliata* | 20.874 | Not endemic | Temperate |
| *Convallaria majalis* | 17.3105 | Not endemic | Temperate |
| *Dichelostemma ida-maia* | 17.787 | Not endemic | Temperate |
| *Dipcadi ursulae* | 14.2737 | Endemic | Tropical |
| *Drimia coromandeliana* | 19.992 | Endemic | Tropical |
| *Eucomis bicolor* | 12.593 | Not endemic | Subtropical |
| *Eucomis comosa* | 23.765 | Not endemic | Subtropical |
| *Eucomis humilis* | 23.422 | Not endemic | Subtropical |
| *Eucomis montana* | 23.863 | Not endemic | Subtropical |
| *Eucomis pallidiflora* | 22.736 | Not endemic | Subtropical |
| *Eucomis regia* | 15.337 | Endemic | Subtropical |
| *Fessia bisotunensis* | 22.246 | Endemic | Temperate |
| *Fessia furseorum* | 16.66 | Endemic | Temperate |
| *Fessia gorganica* | 15.582 | Endemic | Temperate |
| *Fessia greilhuberi* | 12.642 | Endemic | Temperate |
| *Fessia puschkinioides* | 12.152 | Not endemic | Temperate |
| *Hosta hypoleuca* | 12.495 | Endemic | Temperate |
| *Hosta kiyosumiensis* | 11.907 | Endemic | Temperate |
| *Hosta plantaginea* | 12.103 | Not endemic | Temperate |
| *Hosta ventricosa* | 19.208 | Not endemic | Temperate |
| *Hyacinthella nervosa* | 13.8572 | Not endemic | Subtropical |
| *Hyacinthoides hispanica* | 23.6676 | Not endemic | Temperate |
| *Hyacinthoides non-scripta* | 20.734 | Not endemic | Temperate |
| *Hyacinthus orientalis* | 20.874 | Not endemic | Temperate |
| *Liriope spicata* | 12.544 | Not endemic | Subtropical |
| *Maianthemum bifolium* | 14.994 | Not endemic | Temperate |
| *Maianthemum canadense* | 14.847 | Not endemic | Temperate |
| *Maianthemum dilatatum* | 16.366 | Not endemic | Temperate |
| *Maianthemum japonicum* | 20.139 | Not endemic | Temperate |
| *Maianthemum racemosum* | 16.611 | Not endemic | Temperate |
| *Maianthemum stellatum* | 13.034 | Not endemic | Temperate |
| *Ornithogalum alpigenum* | 12.0783 | Endemic | Temperate |
| *Ornithogalum arabicum* | 17.983 | Not endemic | Subtropical |
| *Ornithogalum exscapum* | 30.037 | Not endemic | Subtropical |
| *Ornithogalum montanum* | 30.478 | Not endemic | Temperate |
| *Ornithogalum narbonense* | 32.781 | Not endemic | Temperate |
| *Ornithogalum nutans* | 21.511 | Not endemic | Temperate |
| *Ornithogalum pamphylicum* | 13.6431 | Endemic | Temperate |
| *Ornithogalum pyrenaicum* | 15.239 | Not endemic | Temperate |
| *Ornithogalum sigmoideum* | 17.297 | Not endemic | Temperate |
| *Ornithogalum umbellatum* | 24.353 | Not endemic | Temperate |
| *Ornithogalum wiedemannii* | 13.7898 | Not endemic | Temperate |
| *Peliosanthes macrostegia* | 12.05385 | Not endemic | Subtropical |
| *Peliosanthes ophiopogonoides* | 14.35704 | Endemic | Subtropical |
| *Peliosanthes sinica* | 12.59175 | Not endemic | Subtropical |
| *Peliosanthes yunnanensis* | 12.00495 | Not endemic | Subtropical |
| *Polygonatum biflorum* | 20.384 | Not endemic | Temperate |
| *Polygonatum franchetii* | 30.4595 | Not endemic | Temperate |
| *Polygonatum multiflorum* | 15.043 | Not endemic | Temperate |
| *Reineckea carnea* | 23.422 | Not endemic | Temperate |
| *Rohdea japonica* | 52.323 | Not endemic | Temperate |
| *Scilla amoena* | 23.226 | Endemic | Temperate |
| *Scilla bithynica* | 22.396 | Not endemic | Temperate |
| *Scilla cilicica* | 28.028 | Not endemic | Subtropical |
| *Scilla ingridiae* | 23.422 | Endemic | Temperate |
| *Scilla mischtschenkoana* | 21.168 | Not endemic | Temperate |
| *Scilla peruviana* | 17.738 | Not endemic | Subtropical |
| *Scilla pneumonanthe* | 15.974 | Endemic | Temperate |
| *Scilla rosenii* | 23.324 | Not endemic | Temperate |
| *Scilla siberica* | 31.066 | Not endemic | Temperate |
| *Sowerbaea alliacea* | 12.446 | Endemic | Tropical |
| *Sowerbaea subtilis* | 16.562 | Endemic | Tropical |
| *Zagrosia persica* | 20.482 | Not endemic | Temperate |
| **Asphodelaceae** | |  |  |
| *Aloe ammophila* | 19.502 | Endemic | Desert |
| *Aloe amudatensis* | 16.66 | Not endemic | Tropical |
| *Aloe bergeriana* | 13.328 | Not endemic | Subtropical |
| *Aloe chabaudii* | 17.934 | Not endemic | Tropical |
| *Aloe claviflora* | 16.513 | Not endemic | Desert |
| *Aloe cryptopoda* | 14.161 | Not endemic | Tropical |
| *Aloe davyana* | 14.474 | Not endemic | Subtropical |
| *Aloe ecklonis* | 13.524 | Not endemic | Subtropical |
| *Aloe forbesii* | 16.464 | Endemic | Desert |
| *Aloe glauca* | 15.68 | Endemic | Subtropical |
| *Aloe globuligemma* | 16.611 | Not endemic | Tropical |
| *Aloe haworthioides* | 14.749 | Endemic | Desert |
| *Aloe humilis* | 16.528 | Endemic | Subtropical |
| *Aloe jeppeae* | 14.963 | Not endemic | Subtropical |
| *Aloe krapohliana* | 17.311 | Endemic | Desert |
| *Aloe linearifolia* | 12.936 | Not endemic | Subtropical |
| *Aloe longistyla* | 15.582 | Endemic | Subtropical |
| *Aloe macrosiphon* | 17.934 | Not endemic | Tropical |
| *Aloe melanacantha* | 12.299 | Endemic | Desert |
| *Aloe micracantha* | 13.916 | Not endemic | Subtropical |
| *Aloe parvula* | 16.562 | Endemic | Tropical |
| *Aloe petricola* | 15.092 | Endemic | Desert |
| *Aloe polyphylla* | 13.377 | Endemic | Subtropical |
| *Aloe prinslooi* | 17.444 | Endemic | Subtropical |
| *Aloe prostrata* | 20.139 | Endemic | Tropical |
| *Aloe rauhii* | 15.337 | Endemic | Desert |
| *Aloe vera* | 16.072 | Endemic | Desert |
| *Aloe welwitschii* | 13.328 | Endemic | Tropical |
| *Aristaloe aristata* | 13.4855 | Not endemic | Subtropical |
| *Astroloba bullulata* | 14.896 | Endemic | Subtropical |
| *Astroloba congesta* | 15.435 | Endemic | Subtropical |
| *Bulbine asphodeloides* | 14.504 | Not endemic | Desert |
| *Bulbine bulbosa* | 13.426 | Not endemic | Subtropical |
| *Bulbine frutescens* | 14.651 | Not endemic | Desert |
| *Bulbine glauca* | 17.689 | Not endemic | Subtropical |
| *Bulbine praemorsa* | 12.21325 | Not endemic | Subtropical |
| *Bulbinella latifolia* | 12.152 | Endemic | Subtropical |
| *Gasteria acinacifolia* | 18.963 | Endemic | Subtropical |
| *Gasteria batesiana* | 21.076 | Not endemic | Desert |
| *Gasteria baylissiana* | 17.689 | Endemic | Subtropical |
| *Gasteria brachyphylla* | 17.213 | Endemic | Desert |
| *Gasteria carinata* | 18.68 | Endemic | Subtropical |
| *Gasteria croucheri* | 18.914 | Not endemic | Subtropical |
| *Gasteria disticha* | 17.493 | Endemic | Subtropical |
| *Gasteria ellaphieae* | 17.199 | Endemic | Subtropical |
| *Gasteria excelsa* | 18.326 | Endemic | Subtropical |
| *Gasteria glauca* | 17.395 | Endemic | Subtropical |
| *Gasteria glomerata* | 17.395 | Endemic | Subtropical |
| *Gasteria nitida* | 15.582 | Endemic | Subtropical |
| *Gasteria pillansii* | 17.897 | Not endemic | Desert |
| *Gasteria polita* | 17.346 | Endemic | Subtropical |
| *Gasteria pulchra* | 15.582 | Endemic | Subtropical |
| *Gasteria vlokii* | 17.346 | Endemic | Subtropical |
| *Haworthia angustifolia* | 11.907 | Endemic | Subtropical |
| *Haworthia arachnoidea* | 11.8825 | Endemic | Subtropical |
| *Haworthia magnifica* | 11.90226 | Endemic | Subtropical |
| *Haworthia mucronata* | 11.8335 | Endemic | Subtropical |
| *Haworthia nortieri* | 12.005 | Endemic | Subtropical |
| *Haworthia pubescens* | 12.005 | Endemic | Subtropical |
| *Haworthia semiviva* | 11.956 | Endemic | Desert |
| *Haworthiopsis attenuata* | 12.8607 | Endemic | Subtropical |
| *Haworthiopsis bruynsii* | 14.357 | Endemic | Subtropical |
| *Haworthiopsis koelmaniorum* | 14.847 | Endemic | Desert |
| *Haworthiopsis limifolia* | 16.3326 | Not endemic | Subtropical |
| *Haworthiopsis longiana* | 12.936 | Endemic | Subtropical |
| *Kniphofia uvaria* | 13.456 | Endemic | Temperate |
| **Asteraceae** |  |  |  |
| *Artemisia copa* | 15.4085 | Not endemic | NA |
| *Artemisia macrantha* | 13.02207 | Not endemic | Temperate |
| *Echinacea pallida* | 13.328 | Not endemic | Temperate |
| *Helianthus agrestis* | 12.74 | Not endemic | NA |
| *Kleinia cephalophora* | 14.896 | Not endemic | NA |
| *Kleinia grantii* | 14.2461667 | Not endemic | NA |
| *Leucanthemopsis alpina* | 12.3137 | Not endemic | NA |
| *Leucanthemum heterophyllum* | 20.776 | Not endemic | NA |
| *Leucanthemum illyricum* | 15.827 | Endemic | NA |
| *Leucanthemum pachyphyllum* | 21.952 | Not endemic | Temperate |
| *Leucanthemum subglaucum* | 24.304 | Endemic | Temperate |
| *Rudbeckia laciniata* | 14.9646 | Not endemic | Temperate |
| *Santolina chamaecyparissus* | 14.3962 | Not endemic | NA |
| **Berberidaceae** |  |  |  |
| *Anchusa ochroleuca* | 12.4206 | Not endemic | Temperate |
| *Gymnospermium scipetarum* | 14.4256 | Not endemic | Temperate |
| *Podophyllum hexandrum* | 14.381 | Not endemic | Temperate |
| *Podophyllum peltatum* | 26.117 | Not endemic | Temperate |
| **Colchicaceae** |  |  |  |
| *Disporum sessile* | 18.228 | Not endemic | Temperate |
| *Disporum uniflorum* | 24.516 | Not endemic | Temperate |
| *Uvularia grandiflora* | 15.6985 | Not endemic | Temperate |
| *Uvularia perfoliata* | 12.74 | Not endemic | Temperate |
| **Commelinaceae** | |  |  |
| *Callisia rosea* | 37.926 | Not endemic | Subtropical |
| *Dichorisandra thyrsiflora* | 16.66 | Not endemic | Tropical |
| *Gibasis matudae* | 17.444 | Not endemic | Tropical |
| *Gibasis pulchella* | 20.678 | Not endemic | Subtropical |
| *Tradescantia ambigua* | 30.38 | Not endemic | Tropical |
| *Tradescantia andrieuxii* | 27.538 | Not endemic | Tropical |
| *Tradescantia buckleyi* | 22.932 | Not endemic | Desert |
| *Tradescantia ernestiana* | 19.992 | Not endemic | Temperate |
| *Tradescantia gigantea* | 29.4 | Not endemic | Subtropical |
| *Tradescantia hirsuticaulis* | 21.168 | Not endemic | Temperate |
| *Tradescantia hirta* | 23.422 | Endemic | Tropical |
| *Tradescantia llamasii* | 15.386 | Not endemic | Tropical |
| *Tradescantia longipes* | 40.964 | Not endemic | Temperate |
| *Tradescantia nuevoleonensis* | 12.348 | Endemic | Desert |
| *Tradescantia ohiensis* | 20.188 | Not endemic | Temperate |
| *Tradescantia poelliae* | 13.524 | Not endemic | Tropical |
| *Tradescantia sillamontana* | 17.738 | Endemic | Subtropical |
| *Tradescantia subaspera* | 39.102 | Not endemic | Temperate |
| *Tradescantia tharpii* | 38.318 | Not endemic | Temperate |
| *Tradescantia virginiana* | 42.532 | Not endemic | Temperate |
| **Convolvulaceae** | |  |  |
| *Cuscuta exaltata* | 20.5114 | Not endemic | Subtropical |
| *Cuscuta indecora* | 32.1146 | Not endemic | Temperate |
| *Cuscuta japonica* | 25.584 | Not endemic | Temperate |
| *Cuscuta lupuliformis* | 22.0157 | Not endemic | Temperate |
| *Cuscuta monogyna* | 32.45 | Not endemic | Temperate |
| **Cynomoriaceae** | |  |  |
| *Cynomorium coccineum* | 13.35459 | Not endemic | Subtropical |
| **Ericaceae** |  |  |  |
| *Monotropa uniflora* | 29.302 | Not endemic | Temperate |
| **Fabaceae** |  |  |  |
| *Lathyrus vestitus* | 14.308 | Not endemic | Subtropical |
| *Vicia faba* | 13.034 | Not endemic | Temperate |
| **Iridaceae** |  |  |  |
| *Iris danfordiae* | 15.827 | Endemic | Subtropical |
| *Iris florentina* | 12.14676 | Not endemic | Subtropical |
| *Iris graeberiana* | 12.446 | Endemic | Temperate |
| *Iris hoogiana* | 14.82159 | Not endemic | Temperate |
| *Iris lineata* | 14.55753 | Not endemic | Temperate |
| *Iris stolonifera* | 14.78736 | Not endemic | Temperate |
| *Iris tuberosa* | 13.524 | Not endemic | Subtropical |
| *Moraea atropunctata* | 14.014 | Endemic | Subtropical |
| *Moraea bifida* | 12.446 | Endemic | Subtropical |
| *Moraea calcicola* | 15.484 | Endemic | Subtropical |
| *Moraea flaccida* | 17.542 | Endemic | Subtropical |
| *Moraea radians* | 13.426 | Endemic | Subtropical |
| *Moraea tulbaghensis* | 30.772 | Endemic | Subtropical |
| *Moraea villosa* | 30.772 | Endemic | Subtropical |
| **Liliaceae** |  |  |  |
| *Amana edulis* | 21.0455 | Not endemic | Temperate |
| *Amana erythronioides* | 24.843 | Not endemic | Temperate |
| *Cardiocrinum giganteum* | 38.612 | Not endemic | Temperate |
| *Clintonia borealis* | 18.522 | Not endemic | Temperate |
| *Erythronium albidum* | 74.725 | Not endemic | Temperate |
| *Erythronium americanum* | 73.6715 | Not endemic | Temperate |
| *Erythronium californicum* | 32.91 | Endemic | Temperate |
| *Erythronium dens-canis* | 24.4902 | Not endemic | Temperate |
| *Erythronium japonicum* | 31.115 | Not endemic | Temperate |
| *Fritillaria acmopetala* | 64.925 | Not endemic | Subtropical |
| *Fritillaria affinis* | 45.031 | Not endemic | Temperate |
| *Fritillaria alfredae* | 55.844 | Not endemic | Subtropical |
| *Fritillaria assyriaca* | 75.999 | Not endemic | Temperate |
| *Fritillaria aurea* | 80.262 | Endemic | Temperate |
| *Fritillaria bucharica* | 44.2078 | Not endemic | Temperate |
| *Fritillaria camschatcensis* | 55.0025 | Not endemic | Temperate |
| *Fritillaria crassifolia* | 65.219 | Not endemic | Temperate |
| *Fritillaria davidii* | 33.32 | Endemic | Temperate |
| *Fritillaria eastwoodiae* | 47.6525 | Not endemic | Temperate |
| *Fritillaria elwesii* | 50.7738 | Not endemic | Subtropical |
| *Fritillaria fleischeriana* | 75.558 | Endemic | Temperate |
| *Fritillaria gibbosa* | 41.9048 | Not endemic | Temperate |
| *Fritillaria glauca* | 47.8975 | Not endemic | Temperate |
| *Fritillaria graeca* | 64.67 | Not endemic | Subtropical |
| *Fritillaria gussichiae* | 48.461 | Not endemic | Temperate |
| *Fritillaria imperialis* | 45.0898 | Not endemic | Temperate |
| *Fritillaria koidzumiana* | 85.4168 | Endemic | Temperate |
| *Fritillaria kurdica* | 79.968 | Not endemic | Temperate |
| *Fritillaria liliacea* | 43.904 | Endemic | Temperate |
| *Fritillaria maximowiczii* | 33.6042 | Not endemic | Temperate |
| *Fritillaria melananthera* | 52.332 | Endemic | Temperate |
| *Fritillaria meleagris* | 46.354 | Not endemic | Temperate |
| *Fritillaria montana* | 53.4966 | Not endemic | Temperate |
| *Fritillaria olivieri* | 57.379 | Endemic | Temperate |
| *Fritillaria pallidiflora* | 42.826 | Not endemic | Temperate |
| *Fritillaria persica* | 37.0048 | Not endemic | Subtropical |
| *Fritillaria pinardii* | 64.239 | Not endemic | Temperate |
| *Fritillaria pluriflora* | 40.6994 | Endemic | Temperate |
| *Fritillaria pudica* | 65.0475 | Not endemic | Temperate |
| *Fritillaria pyrenaica* | 39.347 | Not endemic | Temperate |
| *Fritillaria raddeana* | 41.7284 | Not endemic | Temperate |
| *Fritillaria ruthenica* | 50.078 | Not endemic | Temperate |
| *Fritillaria sewerzowii* | 43.561 | Not endemic | Temperate |
| *Fritillaria thunbergii* | 38.0975 | Not endemic | Temperate |
| *Fritillaria tubaeformis* | 44.01 | Not endemic | Temperate |
| *Fritillaria uva-vulpis* | 87.416 | Not endemic | Temperate |
| *Fritillaria verticillata* | 40.8072 | Not endemic | Temperate |
| *Gagea lutea* | 19.355 | Not endemic | Temperate |
| *Gagea pratensis* | 16.072 | Not endemic | Temperate |
| *Gagea spathacea* | 22.883 | Not endemic | Temperate |
| *Lilium albanicum* | 33.0652 | Not endemic | Temperate |
| *Lilium amabile* | 13.426 | Not endemic | Temperate |
| *Lilium apertum* | 22.62114 | Not endemic | Temperate |
| *Lilium auratum* | 54.22032 | Endemic | Temperate |
| *Lilium bakerianum* | 28.93413 | Not endemic | Temperate |
| *Lilium basilissum* | 22.71894 | Not endemic | Temperate |
| *Lilium bosniacum* | 33.2367 | Endemic | Temperate |
| *Lilium brownii* | 30.55272 | Not endemic | Temperate |
| *Lilium bulbiferum* | 35.2443203 | Not endemic | Temperate |
| *Lilium callosum* | 64.09323 | Not endemic | Temperate |
| *Lilium canadense* | 46.942 | Not endemic | Temperate |
| *Lilium candidum* | 37.2204 | Not endemic | Temperate |
| *Lilium carniolicum* | 33.0113 | Not endemic | Temperate |
| *Lilium catesbaei* | 68.23995 | Not endemic | Temperate |
| *Lilium cernuum* | 39.03198 | Not endemic | Temperate |
| *Lilium columbianum* | 82.09332 | Not endemic | Temperate |
| *Lilium concolor* | 31.54539 | Not endemic | Temperate |
| *Lilium davidii* | 38.0828 | Not endemic | Temperate |
| *Lilium distichum* | 45.43788 | Not endemic | Temperate |
| *Lilium duchartrei* | 48.86088 | Not endemic | Temperate |
| *Lilium fargesii* | 35.88771 | Not endemic | Temperate |
| *Lilium formosanum* | 35.868 | Endemic | Temperate |
| *Lilium grayi* | 82.43562 | Not endemic | Temperate |
| *Lilium hansonii* | 49.39878 | Endemic | Temperate |
| *Lilium henrici* | 33.3498 | Endemic | Temperate |
| *Lilium henryi* | 41.5716 | Not endemic | Temperate |
| *Lilium jankae* | 32.7761 | Not endemic | Temperate |
| *Lilium japonicum* | 68.95878 | Endemic | Temperate |
| *Lilium lancifolium* | 61.83894 | Not endemic | Temperate |
| *Lilium lankongense* | 42.3474 | Not endemic | Temperate |
| *Lilium leichtlinii* | 45.54546 | Not endemic | Temperate |
| *Lilium leucanthum* | 38.4354 | Not endemic | Temperate |
| *Lilium longiflorum* | 34.496 | Not endemic | Subtropical |
| *Lilium mackliniae* | 46.73862 | Endemic | Temperate |
| *Lilium maculatum* | 39.609 | Endemic | Temperate |
| *Lilium martagon* | 36.4462 | Not endemic | Temperate |
| *Lilium meleagrina* | 25.0368 | Not endemic | Temperate |
| *Lilium michauxii* | 62.86584 | Not endemic | Temperate |
| *Lilium michiganense* | 47.187 | Not endemic | Temperate |
| *Lilium monadelphum* | 50.21541 | Not endemic | Temperate |
| *Lilium nanum* | 37.877 | Not endemic | NA |
| *Lilium nepalense* | 32.60163 | Not endemic | Temperate |
| *Lilium nobilissimum* | 36.3327 | Endemic | Temperate |
| *Lilium occidentale* | 75.57984 | Not endemic | Temperate |
| *Lilium pardalinum* | 61.01742 | Not endemic | Temperate |
| *Lilium pardanthinum* | 25.3791 | Endemic | Temperate |
| *Lilium parryi* | 63.30594 | Not endemic | Temperate |
| *Lilium pensylvanicum* | 41.77038 | Not endemic | Temperate |
| *Lilium philadelphicum* | 57.71178 | Not endemic | Temperate |
| *Lilium primulinum* | 35.72145 | Not endemic | Temperate |
| *Lilium pumilum* | 46.6578 | Not endemic | Temperate |
| *Lilium pyrenaicum* | 38.0534 | Not endemic | Temperate |
| *Lilium regale* | 47.65305 | Endemic | Temperate |
| *Lilium rosthornii* | 31.22754 | Endemic | Temperate |
| *Lilium rubellum* | 63.35484 | Endemic | Temperate |
| *Lilium saluenense* | 27.16884 | Not endemic | Temperate |
| *Lilium sargentiae* | 58.68489 | Endemic | Temperate |
| *Lilium sealyi* | 22.08324 | Not endemic | Temperate |
| *Lilium souliei* | 21.94632 | Not endemic | Temperate |
| *Lilium speciosum* | 54.22521 | Not endemic | Temperate |
| *Lilium sulphureum* | 54.86091 | Not endemic | Temperate |
| *Lilium superbum* | 78.44049 | Not endemic | Temperate |
| *Lilium taliense* | 27.0906 | Endemic | Temperate |
| *Lilium tsingtauense* | 59.95629 | Not endemic | Temperate |
| *Lilium wallichianum* | 42.56256 | Not endemic | Temperate |
| *Lilium wardii* | 55.38414 | Not endemic | Temperate |
| *Medeola virginiana* | 13.916 | Not endemic | Temperate |
| *Notholirion bulbuliferum* | 24.6764 | Not endemic | Temperate |
| *Notholirion thomsonianum* | 36.6814 | Not endemic | Temperate |
| *Tulipa agenensis* | 28.371 | Not endemic | Subtropical |
| *Tulipa alberti* | 25.676 | Not endemic | Temperate |
| *Tulipa aleppensis* | 46.697 | Not endemic | Subtropical |
| *Tulipa altaica* | 22.001 | Not endemic | Temperate |
| *Tulipa anisophylla* | 21.266 | Not endemic | Temperate |
| *Tulipa armena* | 25.382 | Not endemic | Temperate |
| *Tulipa biflora* | 21.07 | Not endemic | Temperate |
| *Tulipa bifloriformis* | 29.057 | Not endemic | Temperate |
| *Tulipa borszczowii* | 20.342 | Not endemic | Temperate |
| *Tulipa butkovii* | 26.411 | Endemic | Temperate |
| *Tulipa carinata* | 26.558 | Not endemic | Temperate |
| *Tulipa clusiana* | 15.729 | Not endemic | Temperate |
| *Tulipa dasystemon* | 25.235 | Not endemic | Temperate |
| *Tulipa dubia* | 26.803 | Not endemic | Temperate |
| *Tulipa ferganica* | 21.952 | Not endemic | Temperate |
| *Tulipa fosteriana* | 25.725 | Not endemic | Temperate |
| *Tulipa gesneriana* | 26.068 | Endemic | Temperate |
| *Tulipa greigii* | 25.284 | Not endemic | Temperate |
| *Tulipa heteropetala* | 19.159 | Not endemic | Temperate |
| *Tulipa heterophylla* | 18.375 | Not endemic | Temperate |
| *Tulipa heweri* | 23.373 | Endemic | NA |
| *Tulipa hissarica* | 20.972 | Not endemic | Temperate |
| *Tulipa hoogiana* | 23.912 | Not endemic | Temperate |
| *Tulipa humilis* | 21.56 | Not endemic | Temperate |
| *Tulipa iliensis* | 20.09 | Not endemic | Temperate |
| *Tulipa ingens* | 26.215 | Not endemic | Temperate |
| *Tulipa intermedia* | 20.2446 | Endemic | Temperate |
| *Tulipa jacquesii* | 25.3791 | Endemic | Temperate |
| *Tulipa julia* | 30.184 | Not endemic | Temperate |
| *Tulipa kaufmanniana* | 22.148 | Not endemic | Temperate |
| *Tulipa kolbintsevii* | 23.52 | Endemic | Temperate |
| *Tulipa kolpakowskiana* | 20.384 | Not endemic | Temperate |
| *Tulipa korolkowii* | 18.914 | Not endemic | Temperate |
| *Tulipa kuschkensis* | 26.117 | Not endemic | Temperate |
| *Tulipa lanata* | 25.725 | Not endemic | Temperate |
| *Tulipa lehmanniana* | 19.502 | Not endemic | Temperate |
| *Tulipa lemmersii* | 17.836 | Endemic | Temperate |
| *Tulipa linifolia* | 12.152 | Not endemic | Temperate |
| *Tulipa montana* | 14.8225 | Not endemic | Temperate |
| *Tulipa orithyioides* | 28.567 | Not endemic | Temperate |
| *Tulipa orphanidea* | 24.108 | Not endemic | Temperate |
| *Tulipa ostrowskiana* | 38.073 | Not endemic | Temperate |
| *Tulipa persica* | 25.676 | Endemic | Temperate |
| *Tulipa praestans* | 22.05 | Endemic | Temperate |
| *Tulipa regelii* | 25.676 | Endemic | Temperate |
| *Tulipa saxatilis* | 26.754 | Not endemic | Subtropical |
| *Tulipa scharipovii* | 18.2886 | Not endemic | Temperate |
| *Tulipa schmidtii* | 28.371 | Not endemic | Temperate |
| *Tulipa sosnowskyi* | 31.066 | Endemic | Temperate |
| *Tulipa sprengeri* | 31.654 | Endemic | Temperate |
| *Tulipa suaveolens* | 30.135 | Not endemic | Temperate |
| *Tulipa sylvestris* | 21.756 | Not endemic | NA |
| *Tulipa systola* | 25.823 | Not endemic | Subtropical |
| *Tulipa tetraphylla* | 19.649 | Not endemic | Temperate |
| *Tulipa turkestanica* | 42.63 | Not endemic | Temperate |
| *Tulipa undulatifolia* | 23.52 | Not endemic | Temperate |
| *Tulipa uniflora* | 18.767 | Not endemic | Temperate |
| *Tulipa urumiensis* | 23.226 | Not endemic | Temperate |
| *Tulipa vvedenskyi* | 25.186 | Not endemic | Temperate |
| **Loranthaceae** |  |  |  |
| *Amylotheca dictyophleba* | 13.426 | Not endemic | Tropical |
| **Melanthiaceae** |  |  |  |
| *Paris incompleta* | 41.4001 | Not endemic | Temperate |
| *Paris japonica* | 149.1854 | Endemic | Temperate |
| *Paris mairei* | 54.7869 | Endemic | Temperate |
| *Paris quadrifolia* | 49.5096 | Not endemic | Temperate |
| *Paris tetraphylla* | 39.9301 | Not endemic | Temperate |
| *Paris thibetica* | 51.4647 | Not endemic | Temperate |
| *Paris verticillata* | 30.5809 | Not endemic | Temperate |
| *Trillium apetalon* | 93.1 | Not endemic | Temperate |
| *Trillium chloropetalum* | 49.9016 | Endemic | Temperate |
| *Trillium maculatum* | 52.773 | Not endemic | Temperate |
| *Trillium smallii* | 83.741 | Not endemic | Temperate |
| *Trillium taiwanense* | 73.2893 | Endemic | Temperate |
| **Orchidaceae** |  |  |  |
| *Anacamptis pyramidalis* | 12.0736 | Not endemic | Temperate |
| *Calanthe tricarinata* | 12.936 | Not endemic | Temperate |
| *Calypso bulbosa* | 12.8680546 | Not endemic | Temperate |
| *Cephalanthera damasonium* | 16.072 | Not endemic | Temperate |
| *Cephalanthera kotschyana* | 17.302287 | Not endemic | Temperate |
| *Cephalanthera longifolia* | 16.464 | Not endemic | Temperate |
| *Cephalanthera rubra* | 15.3272 | Not endemic | Temperate |
| *Corallorhiza maculata* | 15.68 | Not endemic | Temperate |
| *Corallorhiza odontorhiza* | 15.778 | Not endemic | Temperate |
| *Epipactis atrorubens* | 12.999 | Not endemic | Temperate |
| *Epipactis bugacensis* | 12.101 | Not endemic | Temperate |
| *Epipactis helleborine* | 11.98 | Not endemic | Temperate |
| *Epipactis kleinii* | 14.736 | Not endemic | Temperate |
| *Epipactis mairei* | 13.9133 | Not endemic | Temperate |
| *Epipactis microphylla* | 13.578 | Not endemic | Temperate |
| *Epipactis persica* | 13.365 | Not endemic | Subtropical |
| *Habenaria pectinata* | 15.19 | Not endemic | Temperate |
| *Himantoglossum hircinum* | 12.5184 | Not endemic | Temperate |
| *Limodorum abortivum* | 19.0505 | Not endemic | Temperate |
| *Liparis mamillata* | 27.3172925 | Endemic | Subtropical |
| *Liparis purpureoviridis* | 54.8777 | Not endemic | Tropical |
| *Neotinea lactea* | 15.0227255 | Not endemic | Subtropical |
| *Neottia ovata* | 16.317 | Not endemic | Temperate |
| *Orchis anthropophora* | 14.4744 | Not endemic | Temperate |
| *Pterygodium catholicum* | 15.6010999 | Endemic | Subtropical |
| *Rhyncholaelia digbyana* | 13.39371 | Not endemic | Tropical |
| **Paeoniaceae** |  |  |  |
| *Paeonia anomala* | 17.052 | Not endemic | Temperate |
| *Paeonia arietina* | 24.402 | Not endemic | Temperate |
| *Paeonia broteri* | 20.58 | Not endemic | Temperate |
| *Paeonia californica* | 16.366 | Not endemic | Temperate |
| *Paeonia clusii* | 28.42 | Not endemic | Temperate |
| *Paeonia daurica* | 11.8041 | Not endemic | Temperate |
| *Paeonia intermedia* | 16.993 | Not endemic | Temperate |
| *Paeonia kesrouanensis* | 28.567 | Not endemic | Temperate |
| *Paeonia mascula* | 17.444 | Not endemic | Temperate |
| *Paeonia peregrina* | 25.97 | Not endemic | Temperate |
| *Paeonia tenuifolia* | 16.268 | Not endemic | Temperate |
| **Poaceae** |  |  |  |
| *Aegilops juvenalis* | 18.424 | Not endemic | Temperate |
| *Agropyron desertorum* | 12.936 | Not endemic | Temperate |
| *Anthosachne aprica* | 13.818 | Endemic | Temperate |
| *Anthosachne falcis* | 13.5093 | Endemic | Temperate |
| *Anthosachne kingiana* | 12.3284 | Not endemic | Subtropical |
| *Anthosachne solandri* | 13.9944 | Not endemic | Temperate |
| *Anthoxanthum amarum* | 19.36137 | Not endemic | Temperate |
| *Anthoxanthum brunonis* | 13.6269 | NA | Temperate |
| *Anthoxanthum fuscum* | 13.4995 | NA | Temperate |
| *Avena byzantina* | 13.426 | Not endemic | Temperate |
| *Avena sativa* | 12.593 | Not endemic | Temperate |
| *Bromus arizonicus* | 13.5191 | Not endemic | Desert |
| *Bromus grossus* | 13.622 | Not endemic | Temperate |
| *Bromus intermedius* | 12.642 | Not endemic | Subtropical |
| *Bromus pumpellianus* | 12.985 | Not endemic | Temperate |
| *Bromus racemosus* | 13.23 | Not endemic | Temperate |
| *Bromus ramosus* | 12.7629 | Not endemic | Temperate |
| *Bromus secalinus* | 13.72 | Not endemic | Temperate |
| *Bromus setifolius* | 15.974 | Not endemic | Temperate |
| *Elymus athericus* | 14.1321 | Not endemic | Temperate |
| *Elymus dahuricus* | 12.936 | Not endemic | Temperate |
| *Elymus pungens* | 14.7189 | Not endemic | Temperate |
| *Elymus tenuis* | 15.7241 | Not endemic | Temperate |
| *Helictochloa pratensis* | 17.64 | Not endemic | Temperate |
| *Hordeum procerum* | 13.3182 | Not endemic | Temperate |
| *Kengyilia alatavica* | 14.896 | Not endemic | NA |
| *Koeleria tristis* | 14.3227 | Endemic | Temperate |
| *Lachnagrostis ammobia* | 12.4852 | Endemic | Temperate |
| *Lachnagrostis elata* | 13.1369 | Not endemic | Temperate |
| *Lachnagrostis leptostachys* | 12.3529 | Endemic | Temperate |
| *Lachnagrostis pilosa* | 12.059 | Not endemic | Temperate |
| *Lachnagrostis uda* | 12.495 | Endemic | Temperate |
| *Leymus arenarius* | 20.825 | Not endemic | Temperate |
| *Lygeum spartum* | 13.524 | Not endemic | Subtropical |
| *Poa litorosa* | 15.9544 | Not endemic | Temperate |
| *Thinopyrum intermedium* | 12.74 | Not endemic | Temperate |
| *Thinopyrum obtusiflorum* | 22.127 | Not endemic | Temperate |
| *Triticum aestivum* | 16.944 | Not endemic | Temperate |
| **Ranunculaceae** | |  |  |
| *Actaea cimicifuga* | 11.79468 | Not endemic | Temperate |
| *Actaea dahurica* | 12.1272 | Not endemic | Temperate |
| *Actaea simplex* | 12.24456 | Not endemic | Temperate |
| *Adonis aestivalis* | 15.4208671 | Not endemic | Temperate |
| *Adonis annua* | 16.513 | Not endemic | Temperate |
| *Adonis flammea* | 14.58198 | Not endemic | Temperate |
| *Adonis vernalis* | 13.867 | Not endemic | Temperate |
| *Anemonastrum fasciculatum* | 20.776 | Not endemic | Temperate |
| *Anemonastrum flaccidum* | 28.077 | Not endemic | Temperate |
| *Anemonastrum tetrasepalum* | 22.246 | Not endemic | Temperate |
| *Anemone berlandieri* | 12.544 | Not endemic | Temperate |
| *Anemone hortensis* | 12.152 | Not endemic | Subtropical |
| *Anemonoides apennina* | 13.227 | Not endemic | Temperate |
| *Anemonoides blanda* | 13.5195 | Not endemic | Temperate |
| *Anemonoides nemorosa* | 17.6302 | Not endemic | Temperate |
| *Anemonoides quinquefolia* | 21.07 | Not endemic | Temperate |
| *Anemonoides ranunculoides* | 19.551 | Not endemic | Temperate |
| *Anemonoides trifolia* | 20.09 | Not endemic | Temperate |
| *Beesia deltophylla* | 13.08075 | Endemic | Temperate |
| *Eranthis lobulata* | 13.56486 | Endemic | Temperate |
| *Eranthis sibirica* | 27.11016 | Not endemic | Temperate |
| *Eranthis stellata* | 15.258756 | Not endemic | Temperate |
| *Eranthis tanhoensis* | 12.1259775 | Not endemic | Temperate |
| *Halerpestes ruthenica* | 11.858 | Not endemic | Temperate |
| *Helleborus atrorubens* | 14.259 | Endemic | Temperate |
| *Helleborus cyclophyllus* | 14.014 | Not endemic | Temperate |
| *Helleborus dumetorum* | 14.504 | Not endemic | Temperate |
| *Helleborus multifidus* | 13.426 | Not endemic | Temperate |
| *Helleborus niger* | 14.43344 | Not endemic | Temperate |
| *Helleborus odorus* | 13.328 | Not endemic | Temperate |
| *Helleborus orientalis* | 14.523 | Not endemic | Temperate |
| *Helleborus thibetanus* | 16.023 | Not endemic | Temperate |
| *Helleborus torquatus* | 13.818 | Not endemic | Temperate |
| *Helleborus vesicarius* | 13.328 | Endemic | Temperate |
| *Helleborus viridis* | 13.23 | Not endemic | Temperate |
| *Hepatica acutiloba* | 16.415 | Not endemic | Temperate |
| *Hepatica americana* | 17.15 | Not endemic | Temperate |
| *Hepatica asiatica* | 20.678 | Not endemic | Temperate |
| *Hepatica falconeri* | 14.798 | Not endemic | Temperate |
| *Hepatica henryi* | 32.242 | Not endemic | Temperate |
| *Hepatica insularis* | 20.433 | Endemic | Temperate |
| *Hepatica nobilis* | 15.174 | Not endemic | Temperate |
| *Hepatica transsilvanica* | 32.585 | Endemic | Temperate |
| *Pulsatilla rubra* | 13.21767 | Not endemic | Temperate |
| *Ranunculus carolinianus* | 16.366 | Not endemic | Temperate |
| *Ranunculus constantinopolitanus* | 11.858 | Not endemic | Temperate |
| *Ranunculus kochii* | 17.7018 | Not endemic | Temperate |
| *Ranunculus sulphureus* | 12.45483 | Not endemic | NA |
| **Viburnaceae** |  |  |  |
| *Adoxa moschatellina* | 14.014 | Not endemic | Temperate |
| **Zingiberaceae** |  |  |  |
| *Zingiber mioga* | 23.863 | Not endemic | Subtropical |
